# Genomic architecture of three newly isolated unclassified *Butyrivibrio* species elucidate their potential role in the rumen ecosystem

**DOI:** 10.1101/2021.10.08.463653

**Authors:** Kriti Sengupta, Sai Suresh Hivarkar, Nikola Palevich, Prem Prashant Chaudhary, Prashant K. Dhakephalkar, Sumit Singh Dagar

## Abstract

One cellulose-degrading strain CB08 and two xylan-degrading strains XB500-5 and X503 were isolated from buffalo rumen. All the strains were designated as putative novel species of *Butyrivibrio* based on phylogeny, phylogenomy, digital DNA-DNA hybridization, and average nucleotide identity with their closest type strains. The draft genome length of CB08 was ∼3.54 Mb, while X503 and XB500-5 genome sizes were ∼3.24 Mb and ∼3.27 Mb, respectively. Only 68.28% of total orthologous clusters were shared among three genomes, and 40-44% of genes were identified as hypothetical proteins. The presence of genes encoding diverse carbohydrate-active enzymes (CAZymes) exhibited the lignocellulolytic potential of these strains. Further, the genome annotations revealed the metabolic pathways for monosaccharide fermentation to acetate, butyrate, lactate, ethanol, and hydrogen. The presence of genes for chemotaxis, antibiotic resistance, antimicrobial activity, synthesis of vitamins, and essential fatty acid suggested the versatile metabolic nature of these *Butyrivibrio* strains in the rumen environment.

## 1. Introduction

Plants derived biomass contains vast amounts of fermentable sugars trapped in a lignocellulosic complex. These sugars, if efficiently liberated, can be fermented to several value-added products such as essential vitamins and fatty acids or biofuels like hydrogen and ethanol [1]. However, the major limitation of process development is the slow rate of plant fiber degradation, limiting the release of desired sugars [2]. The recent focus on the development of environmentally benign technologies has shifted the focus back to the exploration of microbial strains, which, on their own, can utilize biomass and produce desirable products. The rumen is a natural home to such competent plant polysaccharide-degrading and fermenting microbes, including anaerobic bacteria and fungi [3–5]. Various species of the genera *Ruminococcus, Fibrobacter, Eubacterium, Prevotella*, and *Butyrivibrio* are well-known fiber degraders in the rumen [6]. Amongst, *Butyrivibrio* spp. are metabolically diverse and able to ferment simple or complex sugars into industrially important compounds like butyric acid, bacteriocins, vitamins, and conjugated linoleic acid (CLA) [7,8]. The diverse nature of *Butyrivibrio* strains in terms of sugar utilization patterns, fermentation products, and genome sequences has been reported recently [9].

Only four species of *Butyrivibrio* are known to date, including *B. crossotus, B. fibrisolvens, B. hungatei*, and *B. proteoclasticus*, while many novel strains remain unclassified. *Butyrivibrio* are Gram-positive, curved, motile, and non-sporulating anaerobic bacteria, first described by Bryant and Small [10]. The genus *Butyrivibrio* is well studied for their diverse Carbohydrate-Active enZYmes (CAZymes) that hydrolyze plant-derived polymers such as cellulose, hemicellulose, xylan, and pectin [11–13]. Some of the *Butyrivibrio* spp. are also known for producing health-promoting compounds such as CLA and vitamins. The *B. fibrisolvens* MDT-5 is regarded as one of the fastest and highest CLA producers compared to other ruminal bacteria [7]. Besides, the members of this genus are also reported to produce antibacterial compounds like bacteriocins or lantibiotics, including a cyclic butyrivibriocin produced by *B. fibrisolvens* AR10 [14]. Evidently, the biotechnologically important features of *Butyrivibrio* not only make them relevant for improving the nutritional status of dairy or meat products but also a key candidate for use in various industrial processes.

The latest studies on the exploration of genome-scale analysis have further helped to decipher the genetic diversity, functional genomics, and metabolic gene architecture of several unclassified *Butyrivibrio* strains to understand their role in plant polysaccharides degradation and fermentation. Also, comparative genomics and pan-genome analysis studies have revealed the CAZymes patterns and core genes distribution among rumen *Butyrivibrio* strains [9,13]. Presently, the NCBI database contains around 50 genomes of unclassified *Butyrivibrio*, representing different geographical locations across the world; however, their true diversity remains understudied. Here, we report the genome sequencing, analysis, and genetic insights into the three putative novel *Butyrivibrio* species cultured from the rumen of Indian buffalo.

## 2. Materials and methods

### 2.1. Isolation & characterization of strains

Three strains designated as CB08, XB500-5, and X503 were cultured from the rumen digesta of slaughtered buffalo from Pune, India. The culturing process involved initial enrichment of rumen digesta in cellulose or xylan based bacterial culture media without or with nalidixic acid (final concentration, 500 µg/ml), followed by isolation using anaerobic serum roll bottle method at 37º C (pH 6.8 ± 0.1) [15,16]. Strain CB08 was obtained on cellulose medium without nalidixic acid, while strains XB500-5 and X503 were cultured on xylan medium with nalidixic acid. The cultures were grown on their respective substrates, and their fermentation metabolites like gases, alcohols, volatile fatty acids (VFAs), and other acids were determined after three days of incubation using gas chromatography [17–19].

### 2.2. DNA extraction and 16S rRNA-based phylogenetic analysis

The high-quality genomic DNA was isolated from all three strains using GenElute™ Bacterial Genomic DNA Kit (Sigma-Aldrich) as per the manufacturer’s instructions. The universal primer set (27f and 1492r) was used to amplify the 16S rRNA gene [20] and outsourced for Sanger sequencing to 1st BASE, Singapore. The obtained sequences were compiled, and sequence similarity searches were performed to find the closest neighbor. The phylogenetic tree analysis of the strains along with their nearest matches was performed by the maximum-likelihood method using the IQ-TREE web server (http://iqtree.cibiv.univie.ac.at/) [21]. For this, the compiled sequences were first aligned using ClustalW in BioEdit v7.2.5, and the aligned file was uploaded to the IQ-Tree web-server and analyzed with default parameters. The deposited sequences are available in the GenBank database (NCBI) with accession numbers MF361102, MF361116, and MG551272 for CB08, XB500-5, and X503, respectively.

### 2.3. Genome sequencing and assembly

The isolated DNA was submitted to Sandor Lifesciences, Hyderabad for whole-genome sequencing (WGS) using Illumina Hiseq PE150 (2×150 bp paired-end) platform. The assembly was performed on sample reads by SPAdes version 3.12 [22] using The Pathosystems Resource Integration Center (PATRIC, www.patricbrc.org) [23]. The assembled contigs of the three strains were quality checked using ContEst16S [24] and AmphoraNet [25], following the step-by-step guidelines on their respective web-servers. The assembled contigs of CB08, XB500-5 and X503 are available in the GenBank database (NCBI) with accession numbers RAIR00000000, RAIS00000000 and RAIQ00000000, respectively.

### 2.4. Genome-based phylogenetic analysis and in-silico species delineation

A genome-based phylogenetic tree of all the reported *Butyrivibrio* strains was obtained from the NCBI database (https://www.ncbi.nlm.nih.gov/genome/16211). The genomic similarity between our strains and the closest strains from the NCBI database were calculated using the Orthologous Average Nucleotide Identity Tool (OAT) provided by ChunLab (https://www.ezbiocloud.net/tools/orthoani). Additionally, the genomic similarity between the three strains was calculated using ANI calculators provided by ChunLab (https://www.ezbiocloud.net/tools/ani), Kostas lab (http://enve-omics.ce.gatech.edu/ani) and JGI-IMG (https://ani.jgi.doe.gov/html/calc.php). The Average Amino Acid Identity (AAI) was calculated using the AAI calculator provided by Kostas lab (http://enve-omics.ce.gatech.edu/aai/). To delineate new species, Genome-to-Genome Distance Calculator (http://ggdc.dsmz.de) and Type (Strain) Genome Server (https://tygs.dsmz.de) provided by German Collection of Microorganisms and Cell Cultures (DSMZ, Germany) was used. The nucleotide dot plots were obtained from JGI-IMG using the BLASTn algorithm [26] to compare the genomes of the three strains to obtain a synteny relationship between the genomes. Whereas, the progressive Mauve program in PATRIC was also used at default parameters to validate the deduced relationship. The circular views of the comparative genomes were obtained from the CGView Server (http://cgview.ca) and BRIG tool [27].

### 2.5. Genome annotations

The assembled contigs were annotated by NCBI Prokaryotic Genome Annotation Pipeline (PGAP) [28] and submitted to RAST server [29] for further annotations. The assembled contigs were also submitted to PATRIC 3.6.4 to report genome annotations (Supplementary file 1). The presence of plasmids and prophages were tested using PlasmidFinder [30] and PHAST [31], respectively. The Pfam search was performed to predict the protein families and clusters of orthologous groups (COGs) using WebMGA [32,33]. The family of regulatory proteins was obtained using P2RP (Predicted Prokaryotic Regulatory Proteins) [34], while transporter proteins were detected by the Transporter Classification Database (TCDB) [35]. Antibiotic resistance genes were identified by the CARD database [36], bacteriocin prediction was performed by BAGEL 3 [37], and antimicrobial peptide gene clusters were identified by antiSMASH web-server [38]. The CAZymes were predicted using the dbCAN web server [39] and compared with other type species of *Butyrivibrio* in the CAZy database [40]. KEGG Mapper online tool was used to visualize the metabolic pathways of annotated genes [41]. Genomic islands (GIs) in the genomes were predicted by IslandViewer 4 [42]. NCBI BLASTn was used to find the sequence similarity of the aerotolerant gene with other ruminal bacteria.

### 2.5 Analysis of orthologous genes

The online program OrthoVenn2 (https://orthovenn2.bioinfotoolkits.net/home) was used at the default settings to evaluate the genome-wide comparison and COGs. For the comparison with type strains, the protein sequences of *Butyrivibrio hungatei* DSM14810^T^ (accession number: FRDH00000000) and *Butyrivibrio proteoclasticus* B316^T^ (accession number: NC_014387) from the NCBI database were included for comparison of the COGs.

### 2.6. Gene family evolution

The proteomes of *Butyrivibrio* genomes were downloaded from NCBI and OrthoFinder v2.5.2 [43] was used to determine single-copy orthologous groups (OGs) in the *Butyrivibrio* proteomes. Cluster selection was based on default parameters for determination of species presence with a single protein in each gene cluster. The resulting extracted proteins were subjected to phylogenetic analysis with 1,000 bootstrap replicates with individual maximum likelihood (ML) inferences for each resulting trimmed gene-cluster generated to infer the species tree under a multiple-species coalescent model for full-coverage data. An evolutionary model was selected automatically for each cluster and performed using the MAFFT v7.017 algorithm within Geneious Prime v2021.2.1 [44].

## 3. Results and Discussion

### 3.1. Identification and characterization of strains

The 16S rRNA gene-based phylogeny revealed that the two xylan degrading strains XB500-5 and X503 showed a 97.50-97.56% match with *B. hungatei* JK 615^T^ (=DSM14810) (accession number: NR_025525), whereas cellulose degrading strain CB08 showed 96.29% and 96.21% sequence similarities with *B. proteoclasticus* B316^T^ (accession number: CP001810) and *B. hungatei* JK 615^T^, respectively. Since a cut-off value of 98.65% is accepted for species delineation, these results indicated the putative novel nature of our strains [45]. Amongst, the strains XB500-5 and X503 were similar by sharing 98.88% sequence identity, while both differed from strain CB08 and shared only 97.35% and 97.55% similarity, respectively. The phylogenetic tree analysis further showed the distinctness of our strains, as evident by the presence of individual clades near but separated from their respective closest matches (Fig. 1a). The tree also confirmed the diverse nature of strain CB08 from strains XB500-5 and X503, which despite their high sequence similarity, formed separate clades supported by high bootstrap values. The codon-based phylogenomic tree (Fig. 1b) too recognized the distinct and novel nature of all three strains, which we further examined using genome-based taxonomic analysis described in section 3.3.1.

**Figure 1.**
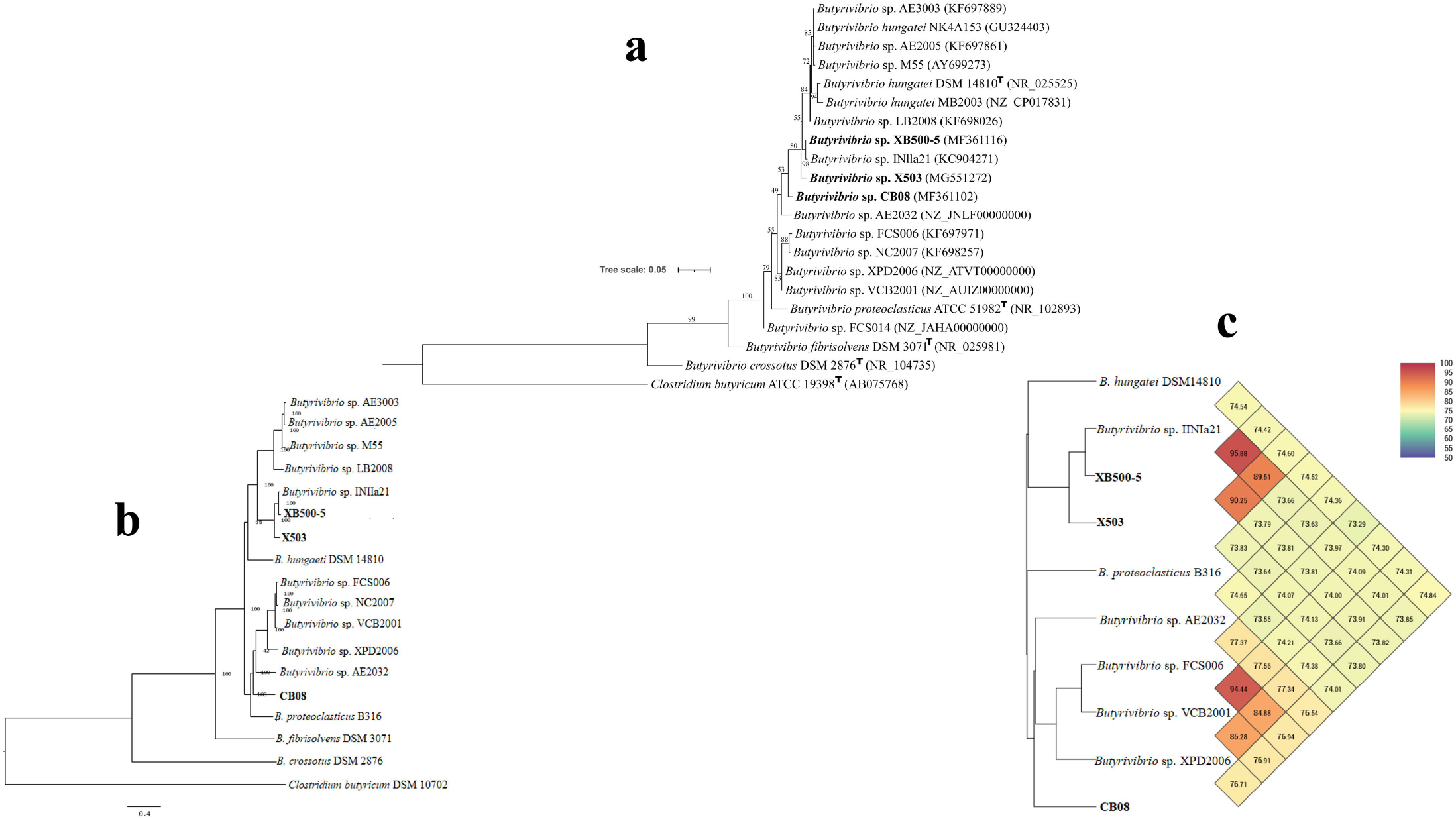
Phylogenetic tree based on 16S rRNA gene sequences (a) phylogenomic tree (b) codon tree and OrthoANI values (% nucleotide level genomic similarity) tree (c) showing the position of strains CB08, XB500-5 and X503 with other *Butyrivibrio* strains. (a) The tree was constructed using the Maximum-Likelihood (ML) method on the IQ-TREE web-server. *Clostridium butyricum* strain ATCC 19398 was used as an outgroup. Bootstrap values based on 1000 replicates are indicated at branching points. The GenBank accession number of each strain is listed in parentheses. Scale bar: 0.05 substitutions per nucleotide position. (b) The tree was generated using the codon tree method within the PATRIC server (https://patricbrc.org/app/PhylogeneticTree). A total of 100 single-copy genes found for 16 genomes, and nucleotide sequences were used for each gene. *Clostridium butyricum* DSM 10702 was used as an outgroup. Bootstrap values based on 500 replicates are indicated at branching points. Scale bar: 0.4 substitutions per nucleotide position.

### 3.2. Fermentation profiles of strains

All the strains produced various acids like formate, acetate, butyrate, propionate, valerate, succinate, and lactate; hydrogen and carbon dioxide as gases and ethanol. The maximum VFA concentrations (in mmol) were obtained in strain CB08 (13.49 ± 0.76), followed by strains XB500-5 (8.15 ± 0.19) and least in strain X503 (6.16 ± 0.13). Acetate was the major VFA in all three strains followed by butyrate, whereas propionate and valerate were produced in minor quantities. Amongst, strain CB08 produced the highest acetate (7.00 ± 1.70) and butyrate (3.86 ± 1.34) followed by strains XB500-5 (acetate: 3.64 ± 0.45, butyrate: 3.43 ± 0.27) and X503 (acetate: 3.21 ± 0.44, butyrate: 1.46 ± 0.03). Besides, all strains produced succinate in traces while minor quantities of formate were detected only in strains XB500-5 and X503. Only the strain CB08 produced lactate as a major acid (31.59 ± 1.11 mmol), implying its possible role in ruminal acidosis [46,47]. The differences in acetate-propionate-butyrate ratio and lactate production by strain CB08 showed its different characteristics over strains XB500-5 and X503. Higher acetate or lactate production instances have also been observed previously in different *Butyrivibrio* spp., but only under certain growth conditions [48]. Moreover, our strains also produced considerable quantities (in mmol) of ethanol (CB08: 23.56 ± 2.3, XB500-5: 34.58 ± 1.1, X503: 16.89 ± 0.3). It is reported that ethanol with concentrations of 0.2% has a moderate inhibitory effect on some *Butyrivibrio* strains, while concentrations of 3.3% completely inhibit the growth [49]. The final concentrations of ethanol produced by the strains CB08 (0.108 ± 0.011%), XB500-5 (0.159 ± 0.005%) and X503 (0.078 ± 0.002%) was way below the inhibitory threshold. While all strains produced carbon dioxide, hydrogen was produced only by the strains CB08 (74.85 ± 1.43 ml/g cellulose) and XB500-5 (46.79 ± 0.51 ml/g xylan). In different species of *Butyrivibrio, B. fibrisolvens* and *B. proteoclasticus* are well-known hydrogen producers, while *B. crossotus* and *B. hungatei* lack this ability [50,51]. The genetic architecture of ethanol and hydrogen production pathways in these strains is detailed in section 3.3.5.

### 3.3. Genomic features of draft genomes

The draft genome sizes of strains XB500-5, X503 and CB08 were 3,273,773 bp, 3,239,716 bp and 3,544,402 bp, respectively. The G+C content of both strains XB500-5 and X503 was 42.4% and 42.2%, respectively, while that of the strain CB08 was 43.7%. The circular genome views of all three genomes are represented in supplementary file 2a and BRIG circular view of gemone sequence comparision with their closest type strains are shown in supplementary file 2b. Their total number of genes, protein-coding genes, rRNA, tRNA, and other general features are listed in Table 1. All the assembled genomes had an L50 value of three, which informed that the longest three contigs together constituted half of the whole genome sequence length. The genomes were further screened for possible contamination with ContEst16S, which revealed the absence of foreign 16 rRNA gene in all the three genomes. Further, AmphoraNet scan detected the presence of 31 bacterial single-copy genes in all the three genomes indicating the completeness of the assembled genome (Supplementary file 3) None of our strains contained any plasmid, which is similar to most *Butyrivibrio* strains [8,13,52].

**Table 1.**
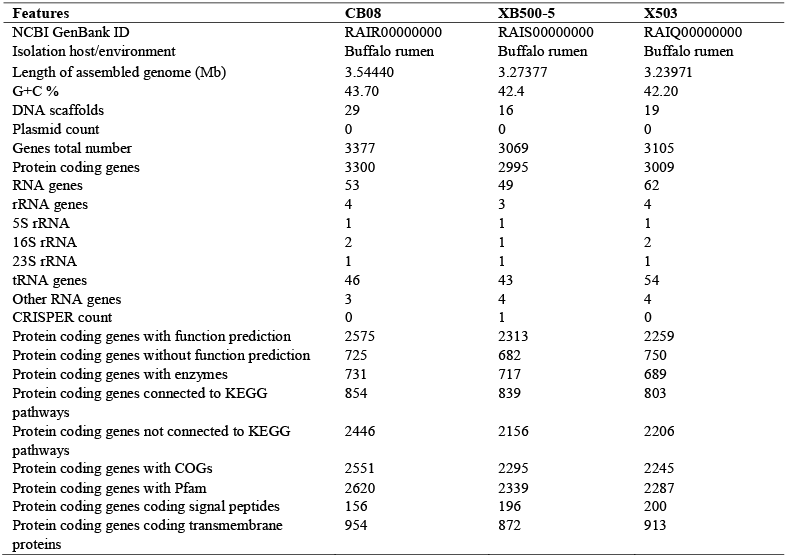
General genomic features of the genome sequence of strains CB08, XB500-5 and X503.

Moreover, a large percentage of genes were annotated as hypothetical proteins in the genomes of CB08 (43.8%), XB500-5 (40.8%), and X503 (42.6%), indicating their involvement in currently unknown functions. A collective genome comparative study of rumen *Butyrivibrio* performed by Palevich *et. al*. (2020) [9] suggested that only 2% collective genome represents the set of survival genes, including energy metabolism and cellular processes. Numerous hypothetical or uncharacterized genes in most rumen *Butrivibrio* probably reflect their unexplored possibility of survival and adaptation in different niches within the rumen.

#### 3.3.1. Genome-derived taxonomy

All strains were identified as novel species of *Butyrivibrio* based on 16S rRNA gene sequence homology, WGS-based ANI and dDDH values. The 16S rRNA gene sequences retrieved from WGS data were identical to the amplified sequences (section 3.1), reconfirming their purity, identity, and novelty. The ANI and dDDH values of the CB08 genome was determined as 76.88% and 15.4%, respectively, to its closest neighbour *B. proteoclasticus* B316^T^. The strains XB500-5 and X503 showed 76.91% and 76.83% ANI value, respectively, with their closest neighbour *B. hungatei* JK 615^T^ and both the strains showed 16.8% dDDH value with the type strain. The lower ANI (<95%) and dDDH (<70%) values also strengthened the novel species claim of these strains as per the existing guidelines [45,53]. The orthoANI values between type strain *Butyrivibrio hungatei* DSM14810 and *Butyrivibrio proteoclasticus* B316, along with the closest genome to our novel strains also supported the novelty amongst the strains (Fig. 1c). The XB500-5 and X503 genome showed approximately 76.5% ANI and 16% dDDH values with the CB08 genome, indicating that CB08 was the most distant among the three strains. Though the 16S rRNA gene similarity between strains XB500-5 and X503 was more than 98.7%, the genome-based ANI (90.62%) and dDDH (66%) values highlighted the distinct species nature of these strains. The detailed ANI and dDDH values between the studied genomes are mentioned in supplementary table S4a and S4b. These results also reflected the importance of genome sequencing in taxonomy studies for the claims of novel species. The TYGS based results of the genome alignment of our strains with their respective type strains also positioned all three strains as new *Butyrivibrio* species (Supplementary file 5). Additionally, the whole genome dot plots were prepared to visualize the gene synteny among the *Butyrivibrio* strains, indicating a significant diversity among the three strains (Fig. 2). The plot showed high genome relatedness between strains XB500-5 and X503, while CB08 showed weaker sequence similarity with the other two strains, this similarity was also confirmed by genome alignment using progressive Mauve (Supplementary file 6). The ANI and dDDH analysis also aligned with this outcome. Thus, the appropriate affiliations of these strains are delineated as new species of genus *Butyrivibrio*, family *Lachnospiraceae*, order *Clostridiales*, class *Clostridia* in phylum *Firmicutes* [10].

**Figure 2.**
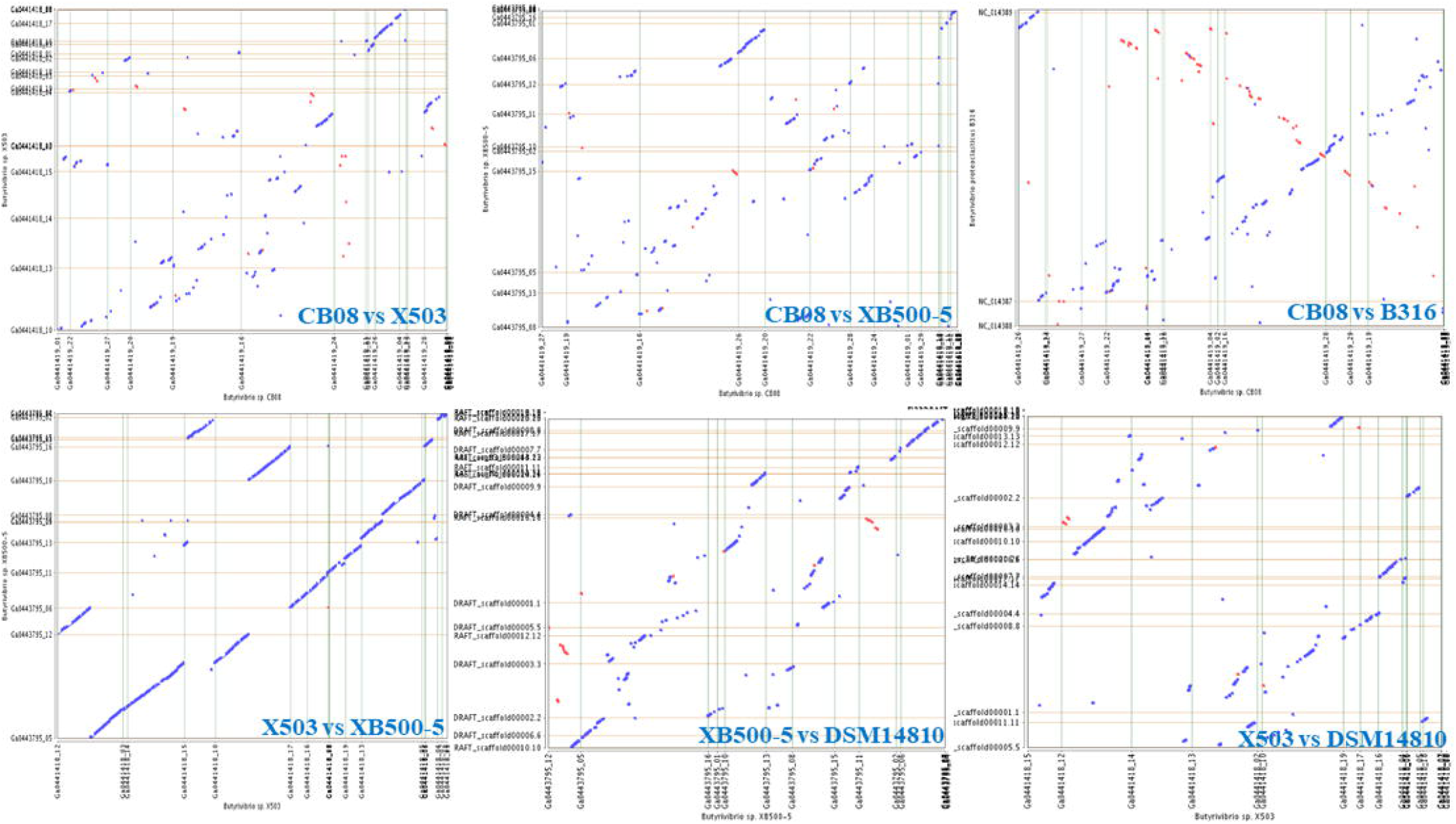
Phylogenetic species-tree reconstruction of homologous gene families and flower plot of unique, group-specific, and core gene families in the *Butyrivibrio* genomes. (a) The consensus tree is based on losses and gains of orthologous groups corresponding to the protein sequence alignments of the 20 *Butyrivibrio* genomes. The predicted protein-coding sequences of all genomes were subjected to an OrthoFinder analysis [44], and the resulting distance matrix was imported into Geneious Prime [45]. The functional distribution was visualized using maximum likelihood (ML) inferences. The tree is drawn to scale, with the branch lengths being in the same units as those of the functional distances used to infer the distribution tree. The bar represents the number of nucleotide substitutions per site. (b) The core genome is shown in the center circle. Each colored segment represents the number of gene families shared among the species groups, and the outer petals represent unique gene families for individual genomes.

#### 3.3.2. COG comparison

The strain CB08 contained the highest protein-coding sequences (3,233), of which 77% were designated to various COG categories. An almost equal number of protein-encoding genes were found in strains XB500-5 (2,957) and X503 (2,958), 75% of which were assigned to different COGs. The relative abundance of proteins assigned to each COG category was comparable to other type species of *Butyrivibrio* [*B. hungatei* NK4A153 (NZ_AUJY00000000), *B. proteoclasticus* B316 (NC_014387), *B. fibrisolvens* DSM3071 (NZ_FQXK00000000), and *B. crossotus* DSM2876 (NZ_ABWN00000000)]. A greater number of genes were predicted in COG categories E (Amino acid transport and metabolism), G (Carbohydrate metabolism and transport), J (translation, ribosomal structure and biogenesis), K (transcription), M (cell wall/membrane/envelope biogenesis), R (general function), S (function unknown), and T (signal transduction mechanism) in comparison to other COG categories. A similar observation was also made in the gene distribution of RAST subsystem annotations, where maximum number of genes were identified in energy metabolism, protein processing and cellular processes (Supplementary file 7). The predicted COG gene distribution pattern was found to be similar with close *Butyrivibrio* spp., except T category gene abundance percentage is exceptionally high in all three strains (8-10% of total) in contrast to other species [8,13,54].

The Venn diagram represented 2749 orthologous gene clusters distributed among the three strains (Fig. 3). A total of 1877 core orthologous gene clusters were conserved in all three genomes, representing that 68.28% of total orthologous clusters are shared among three genomes. The two xylanolytic strains, XB500-5 and X503, shared 2531 (92.07%) orthologous groups among themselves; whereas, this number was only 1947 (70.82%) and 1923 (69.95%) with cellulolytic strain CB08, respectively. The unique orthologous gene clusters identified in strains CB08 (72), XB500-5 (12), and X503 (18) were less than 1% of total orthologous clusters. However, species level uniqueness based on COGs phylogeny has been partially confirmed without complete genome sequence data of these strains.

**Figure 3.**
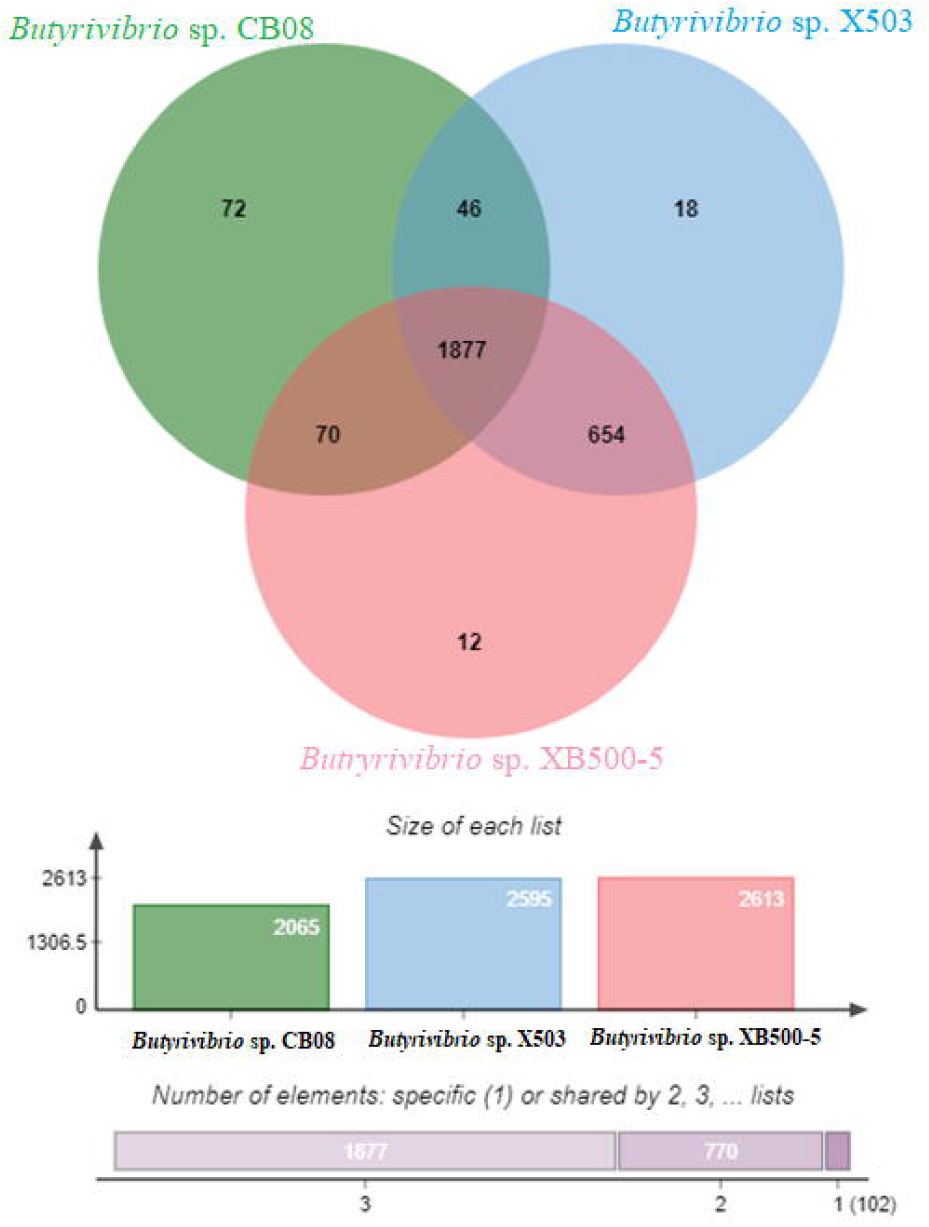
Dot plot representing the gene synteny among the strains CB08, XB500-5 and X503 and their closest type strains *Butyrivibrio proteoclasticus* B316 (=ATCC51982) and *Butyrivibrio hungatei* DSM 14810.

#### 3.3.3. Orthology and evolutionary relationships of Butyrivibrio strains

Comparative orthology profiling and genome alignments based on modelling of gene gain and loss among the *Butyrivibrio* strains [9,13,55] along with *C. butyricum* DSM10702 type strain [56] as an outlier (Fig. 4a), revealed consistent grouping in agreement with the phylogenetic inferences based on the full-length 16*S* rRNA gene sequence data. A total of 18,119 orthologous gene families were found in the *Butyrivibrio* genomes of which 437 represented the core genome set (Fig. 4b), that predominantly consisted of housekeeping and carbohydrate metabolism related genes. *B. fibrisolvens, B. crossotus* and *C. butyricum* genomes clearly separated from the other *Butyrivibrio* based on high numbers of unique and species-specific gene content. *Butyrivibrio* species clearly exhibit a high degree of plasticity at the genome level as reflected by the large size of the collective *Butyrivibrio* pan-genome and unique strain-specific genes. As such, the large reservoir of genetic diversity within this group of organisms further reinforce recent reports of the ability of this group to occupy different niches within the rumen environment [57–59].

**Figure 4.**
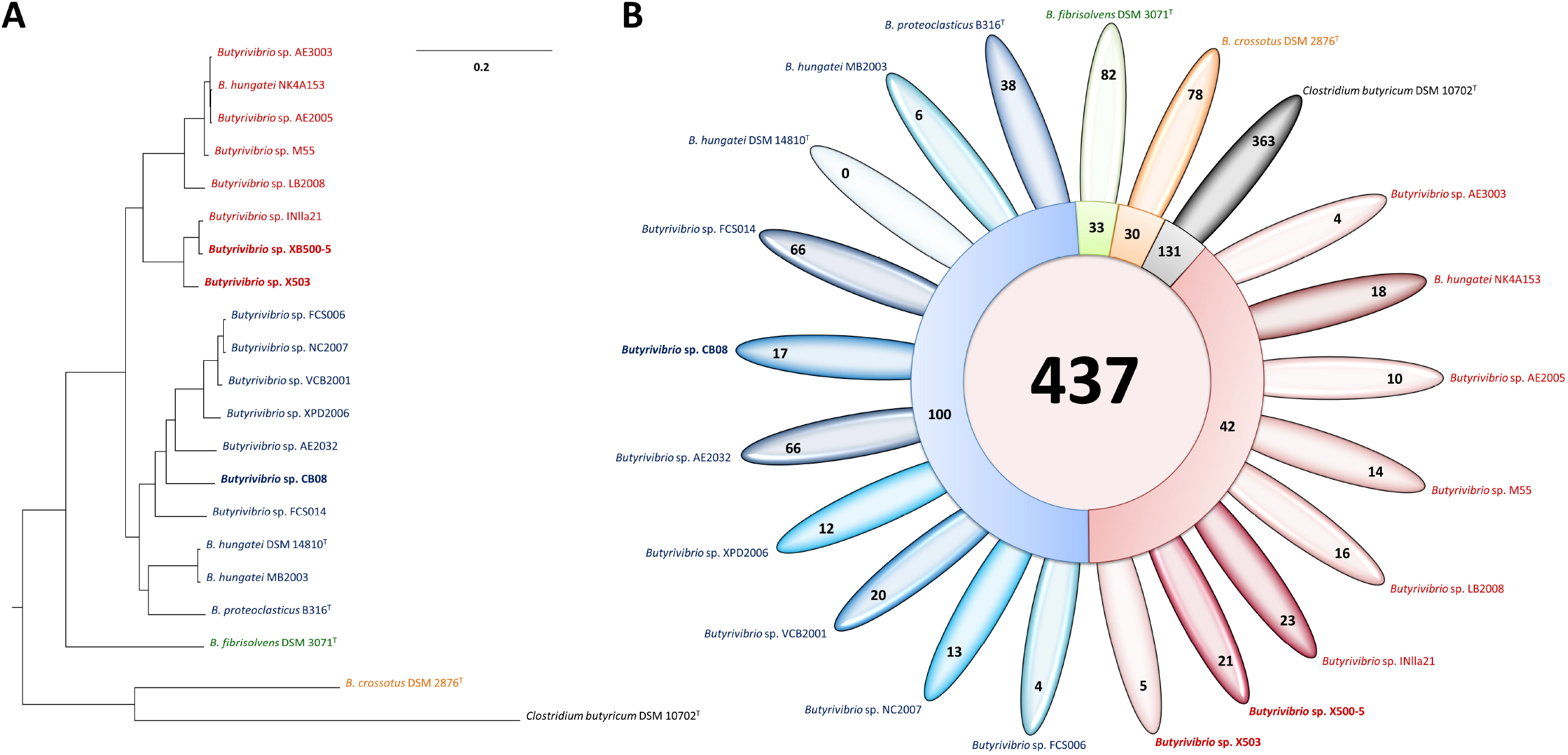
Venn diagram representing the shared orthologous gene clusters in strains CB08, XB500-5 and X503.

#### 3.3.4. Genetics of CAZymes

Different groups of CAZymes were detected in strains CB08, XB500-5, and X503; highest in the strain CB08 and most diverse in strains XB500-5 and X503 (Fig. 5). The comparative CAZyme profiles indicated functional differences of our strains to their respective type strains in polysaccharides degradation. A total of 124 CAZyme genes were identified in strain CB08, constituted by 58 glycoside hydrolases (GHs), 48 glycoside transferases (GTs), 1 polysaccharide lyases (PLs), 10 carbohydrate esterases (CEs), and 9 carbohydrate-binding modules (CBMs). The XB500-5 genome contained 105 total CAZyme families, consisting of 41 GHs and 47 GTs, whereas the X503 genome contained 100 CAZyme families, of which 35 and 48 were GHs and GTs, respectively. Both strains contained 2 PLs, 8 CEs, and 9 CBMs in their genomes (Supplementary file 8). The presence of 47-48 GTs, 1-2 PLs, 8-10 CEs, and 9 CBMs in all three genomes showed their functional similarities towards polysaccharide degradation. The significant abundance of GHs, GTs, CEs, and PLs in these strains displayed their ability to degrade different constituents of plant fiber, such as cellulose, hemicellulose, and pectin. Further, the presence of CBMs in our strains, mainly the CBM2, suggested their role in cellulosome mediated degradation by binding to the cellulose surface similar to other *Butyrivibrio* spp. [60,61]. These results also suggested that all strains could degrade both cellulose and xylan irrespective of substrates used for isolation.

**Figure 5.**
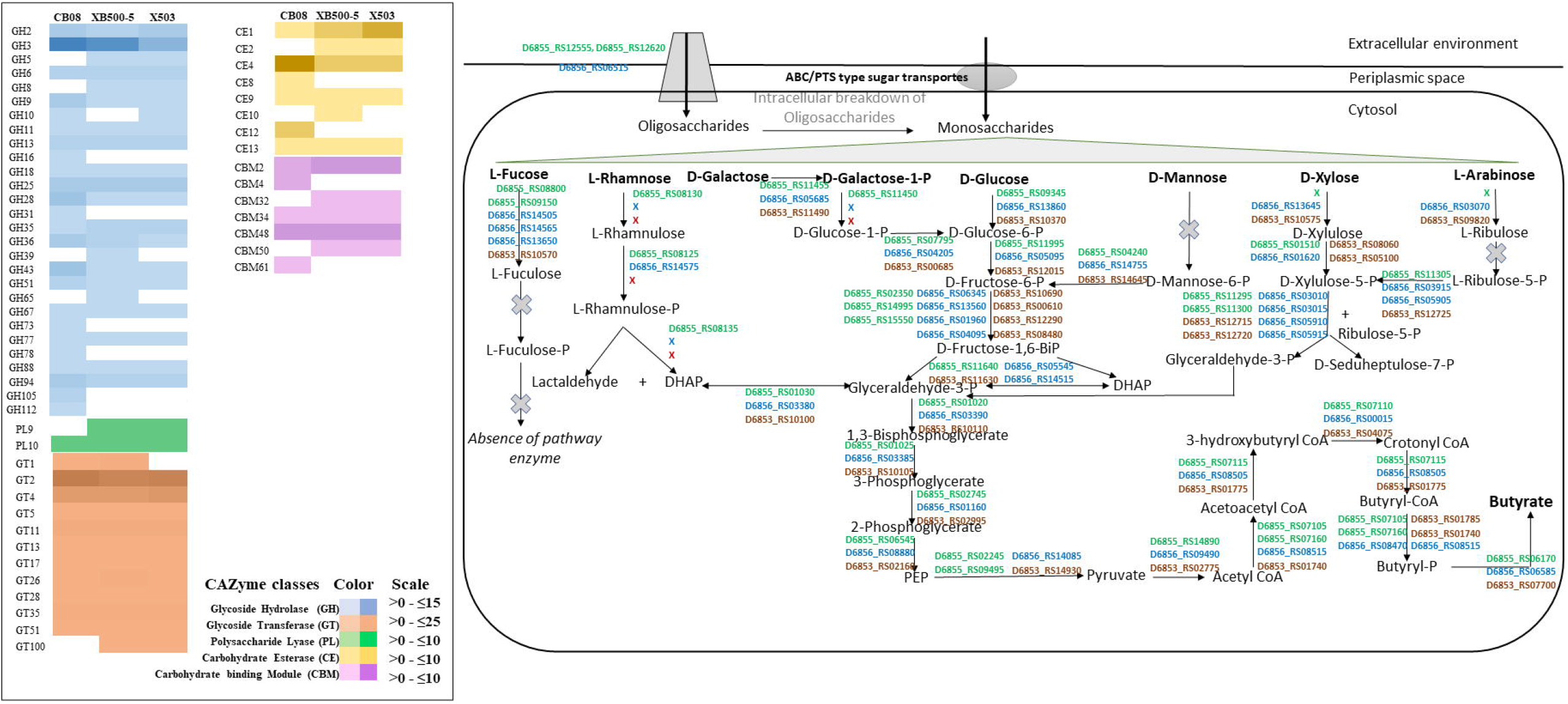
Distribution of gene copy numbers of each CAZymes family protein in strains CB0-8, XB500-5 and X503; gene abundance of CAZymes were predicted from the genome by dbCAN meta server (http://csbl.bmb.uga.edu/dbCAN/annotate.php) and compared with CAZymes gene profile of other *Butyrivibrio* spp. available in the CAZy database (http://www.cazy.org). Predicted pathway model for various sugar fermentation and butyric acid production by *Butyrivibrio* strains based on genetic evidence curated by the KEGG pathway database; NCBI locus of each gene is shown in green, blue and red for strains CB0-8, XB500-5 and X503, respectively. DHAP, dihydroxyacetone phosphate; CoA, Coenzyme A; P, phosphate.

The Pfam search of amino acid sequences of GHs suggested that most CAZymes were without signal peptides, indicating their intracellular nature similar to the previous report [9]. Only a few, like amylolytic GH13 in strain CB08, pectinolytic GH13, and hemicellulolytic GH43 in the other two strains contained signal peptides, showing the extracellular activity. The detailed analysis of GHs also suggested an abundance of GH3 in our strains over compared strains indicating their superior cellobiohydrolase and xylobiohydrolase activity. All the strains contained pectinolytic GH28 and xylanolytic GH11, GH43, GH51, GH67, GH120, which are well reported in *Butyrivibrio* [9,57].

#### 3.3.5. Genetics of fermentative metabolism

The complete pathway genes essential for D-glucose and D-galactose fermentation to acetate and butyrate were identified in all three strains. Besides, genomes of XB500-5 and X503 contained pathway genes required to convert D-xylose and L-arabinose into butyrate, whereas butyrate production from L-rhamnose was specific to CB08 (Fig. 5). The strains XB500-5 and X503 had the genetic construction of the pentose phosphate pathway to utilize arabinose and xylose in contrast to CB08. Such concurrent hexose and pentose sugar utilization have been known in ruminal *Butyrivibrio* [62,63]. The comparison of the butyrate pathway gene organization did not show any substantial difference. The genomes were further annotated with gene *nifJ* encoding pyruvate ferredoxin oxidoreductase (EC 1.2.7.1), which catalyzed the conversion of pyruvate to acetyl CoA. The acetyl CoA often accumulates as an active acetate before its further conversion into butyrate. The butyrate pathway from acetyl CoA usually comprises of acetoacetyl CoA acetyltransferase (EC 2.3.1.9), 3-hydroxybutyryl-CoA dehydrogenase (EC 1.1.1.157), crotonase (EC 4.2.1.17), acetate-CoA transferase (EC 2.8.3.8), and butyrate kinase (EC 2.7.2.7), all of which were detected in our strains. As a butyrate pathway intermediate in these strains, the acetate formation was similar to earlier reports in *Butyrivibrio* [9,13,64]. Another fate of pyruvate to lactate was identified in our strains, where the *ldh* gene encoding lactate dehydrogenase (EC 1.1.1.28) was found responsible for catalyzing the reversible conversion of pyruvate to significant quantities of lactate [65].

The presence of genes encoding pyruvate decarboxylase (PDC, EC 4.1.1.1) and alcohol dehydrogenase (ADH, EC 1.1.1.1) were identified in our strains, which suggested their ability to produce ethanol in two steps. Firstly, PDC catalyzes the decarboxylation of pyruvate to form acetaldehyde, which is then converted to ethanol by ADH in the second step [66,67]. In this process, an electron is donated by NADH and NAD^+^ is formed, which is then reused in glucose to pyruvate conversion. Ethanol production thus helps *Butyrivibrio* in maintaining the NADH: NAD^+^ ratio [68]. Furthermore, the accumulation of acidic metabolites, such as acetic, butyric, lactic, and succinic acids, leads to acidification of culture broth. Hence, *Butyrivibrio* must produce neutral metabolites such as ethanol [69]. Yet, very few reports have documented ethanol as a fermentation product in *Butyrivibrio* [70,71].

The hydrogen is produced as a result of dark fermentation in anaerobic microbes via [FeFe]-hydrogenase [EC 1.12.2.1] mediated reversible reduction of H^+^ to H_2_ coupled with NAD^+^ formation [72,73]. The hydrogenase gene cluster *hypABCDEF* was identified in all three genomes. Though the hydrogenase gene cluster was present in X503, it could not produce hydrogen from xylan under the studied culture conditions. Notably, very few *Butyrivibrio* species are reported to produce hydrogen, and no genetic information of the hydrogenase gene cluster is available [50,51].

#### 3.3.6. Transporters

The rumen environment is rich in mono-, di- or oligo-saccharides because of the action of inherent cellulolytic microflora, including bacteria and gut fungi [74,75]. Similarly, nutrients like amino acids and inhibitors like antibiotics are also present in rumen. Thus, competent carbohydrate transporters should not only influx available sugars and amino acids but also efflux antibiotics [76]. The major class of membrane transporting proteins *viz*., ABC sugar permeases, energy coupling factor transporters (ECF), phosphotransferase system (PTS), major facilitator superfamily (MFS), and oligosaccharide flippase family protein were identified in the genomes (Table 3). Genes for ABC transporters were identified in the strains with COG classes, COG0395, COG0577, COG0601, COG0765, COG0842, COG1129, COG1172, and COG1173, which play crucial roles in the transportation of sugars, amino acids, and multidrug efflux [77]. The genomes also contained abundant gene copies of PTS family proteins (COG1445, COG1762, COG1925, COG2190, COG2213, COG4668), which phosphorylates glucose, fructose, mannose, and other monosaccharides while transporting into the cell [78]. The MFS transporter (COG0659) involved in the transportation of osmolytes, organic ions, sugars, amino acids, and peptides [79], and drug/ metabolite transporter (DMT, COG0697) related to drug and secondary metabolite transportation [80] were also present in the genomes. As *Butyrivibrio* also produce extracellular CAZymes, protein secretion systems are requisite to translocate the enzymes out of the cell [8]. The gene homolog of type II/IV secretory protein (COG4959) was also identified in all the genomes, enabling the secretion of versatile proteins or enzymes in response to the surrounding environments [81]. The abundance of sugar transporter genes in the genomes reflected the remarkable persistency of these strains in lignocellulosic environments like herbivore rumen.

#### 3.3.7. Antimicrobial activity and antibiotic resistance

The rumen bacteria of phylum *Firmicutes*, including *Butyrivibrio*, have been reported to outcompete other microbes by producing different types of inhibitory peptide complexes called bacteriocins or lanthicin [82,83]. Antimicrobial peptides also play an essential role in maintaining the structure of the ruminal microbial community and improve gut health by inhibiting undesirable pathogens [84–86]. The DNA regions encoding secondary metabolites that act as antimicrobial peptides have been detected in all three genomes as ranthipeptide, RiPP type lanthipeptide class II, thiopeptide and beta lactone (Fig. 6a). CB08 genome contained two such regions, one to carry the core biosynthetic gene for ranthipeptide and the other for beta lactone. Ranthipeptide encoding regions were also detected in XB500-5 and X503. However, RiPP type lanthipeptide class II and thiopeptide is detected only in X503 and XB500-5 genome, respectively. Additionally, the gene encoding bacteriocin (COG2274) was detected in the annotated genome of CB08. The presence of genetic machinery for the synthesis of several antibacterial peptides in these strains showed their potential inhibitory action against competing bacteria [82,87].

**Figure 6.**
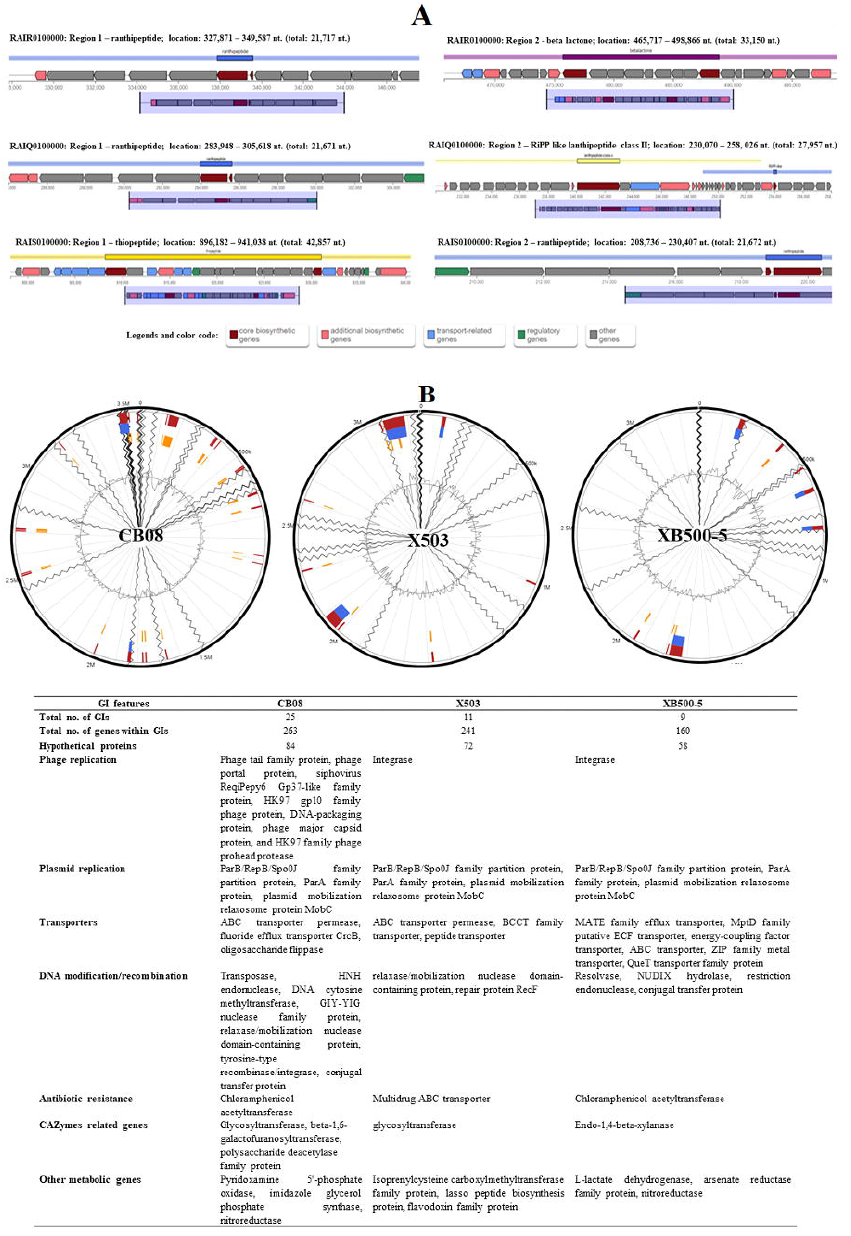
(a) Antimicrobial peptides predicted in strains CB0-8, XB500-5 and X503 using antiSMASH 6.0. Genome accession numbers: RAIR0100000 (CB08), RAIQ0100000 (X503) and RAIS0100000 (XB500-5); (b) Genome Island prediction in strains CB08, XB500-5 and X503. Color code for predicted regions red, blue and yellow signifies three different methods Integrated, Island-Path-DIMOB, and SIGI-HMM, respectively, programmed in IslandViewer 4.

The detailed analysis also discovered the genes associated with antibiotic resistance against nitroimidazole, erythromycin, and aminoglycosides such as tetracycline, kanamycin, streptomycin, and neomycin, which is in line with previous reports [88]. All three genomes contained genes *nimA*/*nimC* that encode a protein capable of inactivating the nitroimidazole, an antimetabolite antibiotic. The *nimA*/*nimC* genes were also observed for the first time in *B. fibrisolvens* DSM 3071^T^ (gene locus BUC27_RS09650) and *B. hungatei* NK4A153^T^ (gene locus G628_RS0103345) but absent in *B. proteoclasticus* B316^T^ and *B. crossotus* DSM 2876^T^. Aminoglycoside cleaving enzyme and erythromycin inactivating gene *ereA* were also annotated in all our strains.

For some quinolone antibiotics like nalidixic acid, the resistance is conferred through amino acid substitutions in parts of the *GyrA* (gyrase) and *ParC* (topoisomerase IV) proteins called quinolone resistance -determining regions (QRDRs) [89,90]. Cultivation of strains X503 and XB500-5 in the nalidixic acid medium indicates the probable presence of such mutations in these strains. Further, several reports do suggest widespread resistance against quinolones in *Butyrivibrio* strains [91–93].

#### 3.3.8. Biosynthesis of essential micronutrients

Micronutrients such as vitamins and essential fatty acids (EFAs) are produced by enteric or gut microorganisms, which enhance nutrient accessibility to the host and gut microflora [94]. Pathway genes for biosynthesis of vitamins such as biotin (vitamin B7), thiamine (vitamin B1), riboflavin (vitamin B2), pyridoxine (vitamin B6), folic acid (vitamin B9), and cobalamin (vitamin B12) were predicted in the genomes of all three strains (Table 3). The key genes for biotin pathway enzymes like biotin-(acetyl-CoA carboxylase) ligase (COG0340), biotin carboxylase (COG0439), biotin carboxyl carrier protein (COG0511), and thiamine synthesis pathways such as thiamine pyrophosphokinase (COG1564), thiamine pyrophosphate-requiring enzymes (COG0028), thiamine biosynthesis ATP pyrophosphatase (COG0301), thiamine monophosphate synthase (COG0352), hydroxymethylpyrimidine/ phosphomethylpyrimidine kinase (COG0351), thiamine biosynthesis protein ThiC (COG0422), thiamin transporter (COG1477), and thiamine-pyrophosphate-binding protein (COG4032) were identified. The COGs for biosynthesis of riboflavin [riboflavin kinase/ FMN adenylyltransferase (COG1853), ATP phosphoribosyltransferase (COG2038)], pyridoxin [pyridoxamine 5′-phosphate oxidase (COG0399)], folate [methenyltetrahydrofolate cyclohydrolase (COG0190), 5-formyltetrahydrofolate cyclo-ligase (COG0212), dihydrofolate reductase (COG0262), folylpolyglutamate synthase (COG0285), methylenetetrahydrofolate reductase (COG0685)] and cobalamin [cobalamin-5-phosphate synthase (COG0368), CbiQ permease (COG1120) and biosynthesis protein like CbiB, -D, -G, -K (COG1270, COG1903, COG2073, COG4822)] were also detected. The presence of the biosynthesis pathways of several vitamins in other rumen *Butyrivibrio* strains has also been reported recently [95]. The traditional cattle feedstuffs contain large amounts of polyunsaturated fatty acid (PUFAs), particularly linolenic acid (LNA, C18:3) and linoleic acid (LA, C18:2), which are toxic to rumen microorganisms. Therefore, several ruminal bacteria biohydrogenates PUFAs to EFAs such as stearic acid as a defence mechanism [96,97], including *Butyrivibrio* species like *B. fibrisolvens* DSM 3071^T^ and *B. hungatei* Su6 [7,13,98]. The presence of genes for linoleate isomerase complex (COG0059), enone reductase (COG0656), and 2-hydroxy-6-oxo-6-phenylhexa-2,4-dienoate hydrolase (COG1575) demonstrated the role of our strains in linoleic acid metabolism.

#### 3.3.9. Chemotaxis, regulons and stress response

Attachment of ruminal bacteria to the ingested feed is important for effective fermentation and assimilation of nutrients in the host. Plant-derived forages often contain chemoattractants like amino acids, sugars, and flavonoids which help in the signalling process to initiate the interactions between feed and bacteria [99–101]. The genome analysis of each strain suggested the presence of several chemotaxis proteins (Table 2) such as CheA, chemotaxis protein histidine kinase (COG0643); CheB, methyl-accepting chemotaxis protein (COG0840); CheC, an inhibitor of MCP methylation (COG1776); CheD, methylation stimulant of MCP proteins (COG1871); CheW, chemotaxis signal transduction protein (COG0835); CheY-like receiver domain (COG2201) and CheR, methylase of chemotaxis methyl-accepting proteins (COG1352). The membrane-bound protein (CheB) acts as a receptor for binding several chemical compounds that trigger chemotaxis signal transduction. The other chemotaxis proteins like CheA, -Y, -W, -Z, and -R are involved in flagellar movement regulation [102]. Chemotaxis in rumen protozoa and anaerobic fungi is well reported [103–105], but despite motile nature and flagellar presence in *Butyrivibrio* chemotaxis is not properly documented [8,106]. This study explains the genetic existence of chemotaxis response in *Butyrivibrio*, which indicated their chemotactic response to the feed. The P2RP analysis, which identifies regulons in the genomes, predicted several regulatory genes involved in two-component systems (TCS), transcription factors, and stress responses in our strains. Regulons for cell motility and quorum sensing (LysR, COG0583), osmoregulation (OmpR, COG0745), carbohydrate metabolism (AraC, COG2207, COG4977, COG1609) [107], polysaccharide degradation (GntR, COG1167, COG1802, COG2186) [108] and multi-drug resistance (MerR, COG0789; TetR, COG1309, COG3226) [109] were predicted. Moreover, the presence of response regulators like the ArsR family (COG0640) and LexA family (COG1974) in the strains indicated their ability to respond to stresses of acidic pH and DNA damage, respectively [110,111].

**Table 2.**
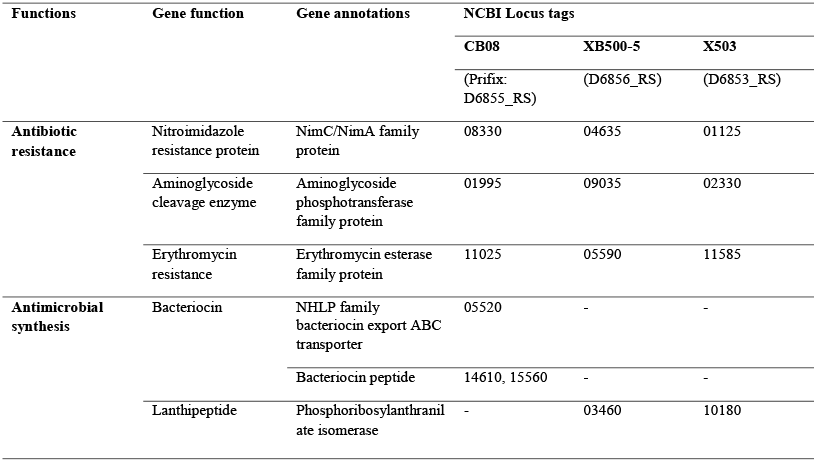
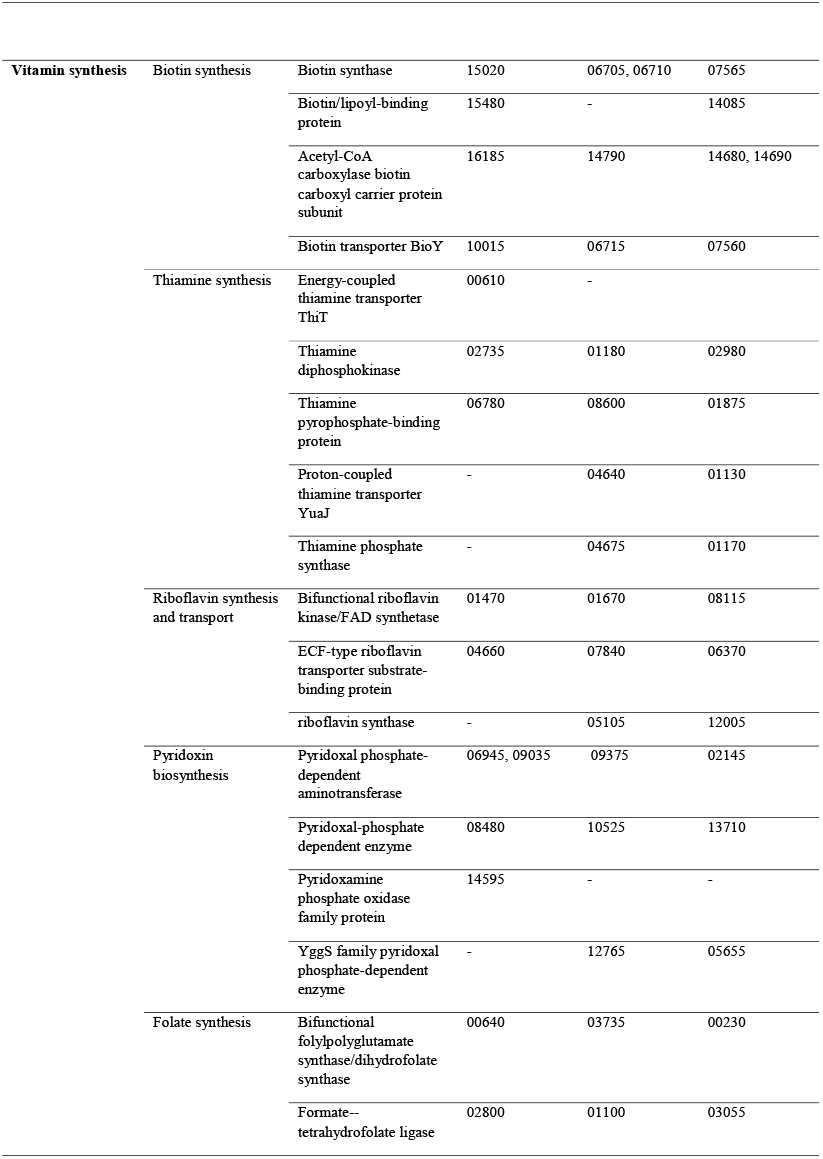
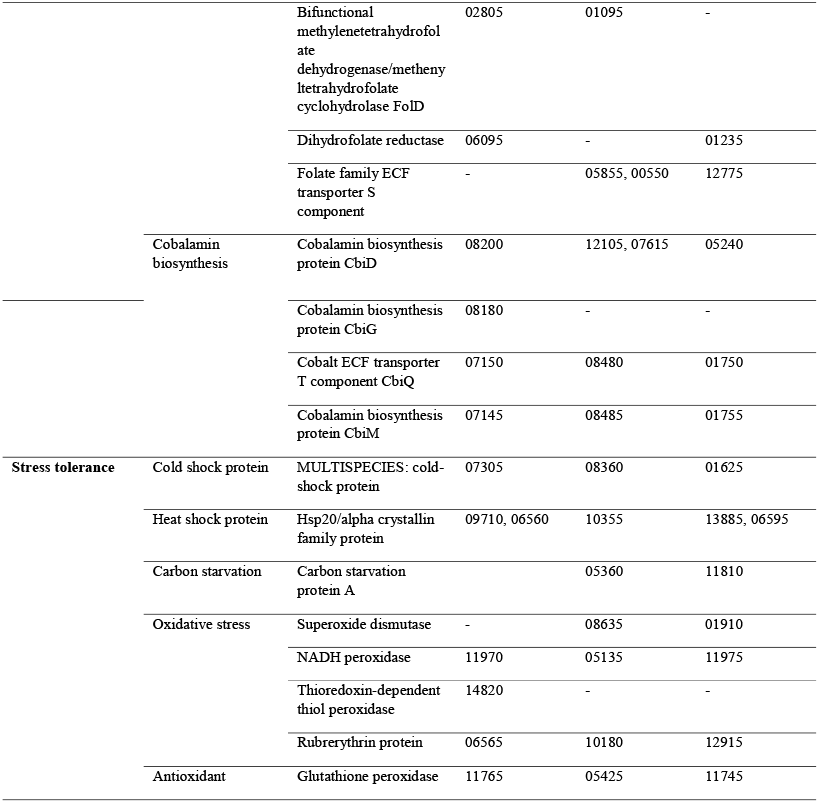
List of genes with functional annotation and gene locus tag identified for antibiotic resistance, vitamin synthesis and other ruminal stress tolerating genes in the genome of strains CB08, XB500-5 and X503.

The presence of stress regulatory genes in all our strains exhibited their ability to survive in adverse conditions like osmotic pressure, acidic environment, and nucleic acid damage. The other stress genes responsible for cold shock protein (COG1278) and heat shock protein (COG0071, COG0576, COG1188) were identified in all the strains, while carbon starvation protein (COG1702) was only detected in strains XB500-5 and X503. The comparative genome analysis of our strains with their respective type strains revealed the presence of oxidative stress tolerance genes for superoxide dismutase (COG0605), rubrerythrin (COG1592), and glutathione peroxidase (COG0386) only in strains XB500-5 and X503. These proteins are associated mainly with aerotolerant microbes, and their presence in *Butyrivibrio* has never have been detailed [112–114]. The presence of these genes in our strains shows their potential ability to tolerate brief air exposures.

#### 3.3.10. Genetics of horizontal gene transfer

Horizontal gene transfer (HGT) refers to the acquisition of foreign genes, mainly in genomic islands (GIs), that may or may not be from related partners and help microbes evolve under changing environmental conditions [115]. The Island Viewer 4 tool predicted GIs involved in functions such as carbohydrate transport and recombination of DNA (Fig. 6b). The maximum number of GIs were observed in strain CB08 (25; total length 232013 bp; genes, 263 including 84 hypothetical proteins) followed by strains X503 (11; total length 231137 bp, bp; genes, 241 including 72 hypothetical proteins) and XB500-5 (9; total length, 154067 bp; genes, 160 including 58 hypothetical proteins). The genes encoding mobile element and plasmid genes were also identified in our strains, which showed sequence similarities with other rumen bacteria. These results indicated that the strains might have undergone genetic recombination through lateral gene transfer from their nearest relatives (Supplementary table S9). A report on the plasmidome of rumen has established the mosaic nature of rumen derived plasmids by various bioinformatics tools and proposed the horizontal gene transfer events among rumen phyla *Firmicutes* [116]. Although these strains did not possess any plasmid, nontransferable incomplete plasmid genes were found scattered in their genomes, remarkably more in strain CB08. The most probable source of these genes was identified as *Clostridium* sp. SY8519 (AP012212), *B. proteoclasticus* B316 (CP001810), *B. hungatei* MB2003 (CP017831), and *B. fibrisolvens* 16/4 (FP929036). Moreover, the CB08 genome contained bacteriophage genes encoding holin, capsid, tail protein, and phage protease, suggesting the acquisition of foreign genes via transduction events (Supplementary file 10). This information is unique in terms of coining the genetic basis of lateral gene transfer in the *Butyrivibrio* species.

## 4. Conclusions

The genus *Butyrivibrio* is an indispensable member of the keystone functional groups in the rumen. The members of this genus are actively involved in several vital functions, including but not limited to fibrolysis and biohydrogenation. The genomic analysis of strains CB08, XB500-5, and X503 not only confirmed their distinct and novel nature but also elucidated their genetic machinery for cellulose, hemicelluloses, and pectin degradation. Similar to recognized members of this genus, acetate, butyrate, propionate, lactate, ethanol, and hydrogen were major fermentation products, the genes of which were successfully mapped in their genome. Furthermore, the presence of genes for chemotaxis, transporters, stress responses, antibiotic resistance, antimicrobial activity, vitamin, and EFAs synthesis in the genomes of these strains highlighted their genetic versatility. In addition to numerous similarities, few unique and specific characters like carbon starvation and aerotolerance in XB500-5 and X503 were also noticed. Interestingly, many genes encoding phage proteins and plasmid mobilization regions in CB08 indicated the integration of foreign genes in this strain. Moreover, complete hydrogenase gene cluster *hypABCDEF* in all strains is reported for the first time in this study. Therefore, these results point towards the inherent diversity and versatility of *Butyrivibrio* members and their role in the rumen ecosystem. Our study also highlights the need for genome-based comparative analyses to comprehensively understand the genetic heterogeneity of this group.

## Data accessibility statement

The 16S rRNA gene sequences are available in the GenBank database with accession numbers MF361102, MF361116, and MG551272 for CB08, XB500-5, and X503, respectively. Genome sequences of each *Butyrivibrio* species strain CB08, XB500-5, and X503 have been deposited at GenBank under the accession number RAIR00000000, RAIS00000000, and RAIQ00000000, respectively.

## Acknowledgements

The authors acknowledge the support of the Director, Agharkar Research Institute, for an institutional research grant (MIC-32). The SERB-National Post-Doctoral Fellowship (N-PDF) Fellowship (F. No. PDF/2019/002389) to KS is thankfully acknowledged. The SERB sponsored project (YSS/2015/000718) based Junior Research Fellowship to SSH is acknowledged.

## CRediT Author contributions

**Kriti Sengupta:** Data analysis, Writing-Original Draft, Data visualization, Writing-Review & Editing **Sai Suresh Hivarkar:** Investigation, Methodology, Data curation, Writing-Review & Editing, Validation **Nikola Palevich:** Data analysis, Review & Editing the final draft **Prem Prashant Chaudhary:** Data curation, Validation **Prashant K. Dhakephalkar:** Conceptualization, Funding acquisition, Resources, Writing-Review & Editing **Sumit Singh Dagar:** Conceptualization, Supervision, Project administration, Funding acquisition, Resources, Validation, Writing-Review & Editing the final draft

## References

[1] A. Arevalo-Gallegos, Z. Ahmad, M. Asgher, R. Parra-Saldivar, H.M.N. Iqbal, Lignocellulose: A sustainable material to produce value-added products with a zero waste approach—A review, Int. J. Biol. Macromol. 99 (2017) 308–318. https://doi.org/10.1016/j.ijbiomac.2017.02.097.

[2] P. Susmel, B. Stefanon, C.R. Mills, M. Spanghero, Rumen degradability of organic matter, nitrogen and fibre fractions in forages, Anim. Prod. 51 (1990) 515–526. https://doi.org/10.1017/S0003356100012551.

[3] S.K. Sirohi, N. Singh, S.S. Dagar, A.K. Puniya, Molecular tools for deciphering the microbial community structure and diversity in rumen ecosystem, Appl. Microbiol. Biotechnol. 95 (2012) 1135–1154. https://doi.org/10.1007/s00253-012-4262-2.

[4] S. Koike, S. Yoshitani, Y. Kobayashi, K. Tanaka, Phylogenetic analysis of fiber-associated rumen bacterial community and PCR detection of uncultured bacteria, FEMS Microbiol. Lett. 229 (2003) 23–30. https://doi.org/10.1016/S0378-1097(03)00760-2.

[5] T.A. McAllister, H.D. Bae, G.A. Jones, K.J. Cheng, Microbial attachment and feed digestion in the rumen., J. Anim. Sci. 72 (1994) 3004–3018. https://doi.org/10.2527/1994.72113004x.

[6] S. Koike, Y. Kobayashi, Fibrolytic rumen bacteria: Their ecology and functions, Asian-Australasian J. Anim. Sci. 22 (2009) 131–138. https://doi.org/10.5713/ajas.2009.r.01.

[7] S. Fukuda, Y. Suzuki, M. Murai, N. Asanuma, T. Hino, Isolation of a novel strain of Butyrivibrio fibrisolvens that isomerizes linoleic acid to conjugated linoleic acid without hydrogenation, and its utilization as a probiotic for animals, J. Appl. Microbiol. 100 (2006) 787–794. https://doi.org/10.1111/j.1365-2672.2006.02864.x.

[8] W.J. Kelly, S.C. Leahy, E. Altermann, C.J. Yeoman, J.C. Dunne, Z. Kong, D.M. Pacheco, D. Li, S.J. Noel, C.D. Moon, A.L. Cookson, G.T. Attwood, The glycobiome of the rumen bacterium butyrivibrio proteoclasticus B316T highlights adaptation to a polysaccharide-rich environment, PLoS One. 5 (2010) e11942. https://doi.org/10.1371/journal.pone.0011942.

[9] N. Palevich, W.J. Kelly, S.C. Leahy, S. Denman, E. Altermann, J. Rakonjac, G.T. Attwood, Comparative genomics of rumen Butyrivibrio spp. uncovers a continuum of polysaccharide-degrading capabilities, Appl. Environ. Microbiol. 86 (2020) 1993– 2012. https://doi.org/10.1128/AEM.01993-19.

[10] M.P. Bryant, N. Small, The anaerobic monotrichous butyric acid-producing curved rod-shaped bacteria of the rumen., J. Bacteriol. 72 (1956) 16–21. https://doi.org/10.1128/jb.72.1.16-21.1956.

[11] R.B. Hespell, M.A. Cotta, Degradation and utilization by Butyrivibrio fibrisolvens H17c of xylans with different chemical and physical properties, Appl. Environ. Microbiol. 61 (1995) 3042–3050. https://doi.org/10.1128/aem.61.8.3042-3050.1995.

[12] A.G. Williams, S.E. Withers, Induction of xylan-degrading enzymes in Butyrivibrio fibrisolvens, Curr. Microbiol. 25 (1992) 297–303. https://doi.org/10.1007/BF01575865.

[13] N. Palevich, W.J. Kelly, S.C. Leahy, E. Altermann, J. Rakonjac, G.T. Attwood, The complete genome sequence of the rumen bacterium Butyrivibrio hungatei MB2003, Stand. Genomic Sci. 12 (2017) 72. https://doi.org/10.1186/s40793-017-0285-8.

[14] M.L. Kalmokoff, T.D. Cyr, M.A. Hefford, M.F. Whitford, R.M. Teather, Butyrivibriocin AR10, a new cyclic bacteriocin produced by the ruminal anaerobe Butyrivibrio fibrisolvens AR10: Characterization of the gene and peptide, Can. J. Microbiol. 49 (2003) 763–773. https://doi.org/10.1139/w03-101.

[15] H.P.S. Makkar, C.S. McSweeney, Methods in gut microbial ecology for ruminants, 2005. https://doi.org/10.1007/1-4020-3791-0.

[16] T.L. Miller, M.J. Wolin, A Serum Bottle Modification of the Hungate Technique for Cultivating Obligate Anaerobes, Appl. Microbiol. 27 (1974) 985–987. https://doi.org/10.1128/aem.27.5.985-987.1974.

[17] A.S. Dighe, Y.S. Shouche, D.R. Ranade, Selenomonas lipolytica sp. nov., an obligately anaerobic bacterium possessing lipolytic activity, Int. J. Syst. Bacteriol. 48 (1998) 783–791. https://doi.org/10.1099/00207713-48-3-783.

[18] L.B. Kamalaskar, P.K. Dhakephalkar, K.K. Meher, D.R. Ranade, High biohydrogen yielding Clostridium sp. DMHC-10 isolated from sludge of distillery waste treatment plant, Int. J. Hydrogen Energy. 35 (2010) 10639–10644. https://doi.org/10.1016/j.ijhydene.2010.05.020.

[19] K.G. Singh, K.L. Lapsiya, R.R. Gophane, D.R. Ranade, Optimization for butanol production using Response Surface Methodology by Clostridium beijerenckii strain CHTa isolated from distillery waste manure., J. Biochem. Technol. 7 (2016) 1063– 1068.

[20] M.T. Suzuki, S.J. Giovannoni, Bias caused by template annealing in the amplification of mixtures of 16S rRNA genes by PCR, Appl. Environ. Microbiol. 62 (1996) 625– 630. https://doi.org/10.1128/aem.62.2.625-630.1996.

[21] B.Q. Minh, H.A. Schmidt, O. Chernomor, D. Schrempf, M.D. Woodhams, A. Von Haeseler, R. Lanfear, E. Teeling, IQ-TREE 2: New Models and Efficient Methods for Phylogenetic Inference in the Genomic Era, Mol. Biol. Evol. 37 (2020) 1530–1534. https://doi.org/10.1093/molbev/msaa015.

[22] A. Bankevich, S. Nurk, D. Antipov, A.A. Gurevich, M. Dvorkin, A.S. Kulikov, V.M. Lesin, S.I. Nikolenko, S. Pham, A.D. Prjibelski, A. V. Pyshkin, A. V. Sirotkin, N. Vyahhi, G. Tesler, M.A. Alekseyev, P.A. Pevzner, SPAdes: A new genome assembly algorithm and its applications to single-cell sequencing, J. Comput. Biol. 19 (2012) 455–477. https://doi.org/10.1089/cmb.2012.0021.

[23] A.R. Wattam, T. Brettin, J.J. Davis, S. Gerdes, R. Kenyon, D. Machi, C. Mao, R. Olson, R. Overbeek, G.D. Pusch, M.P. Shukla, R. Stevens, V. Vonstein, A. Warren, F. Xia, H. Yoo, Assembly, annotation, and comparative genomics in PATRIC, the all bacterial bioinformatics resource center, in: Methods Mol. Biol., Humana Press Inc., 2018: pp. 79–101. https://doi.org/10.1007/978-1-4939-7463-4_4.

[24] I. Lee, M. Chalita, S.M. Ha, S.I. Na, S.H. Yoon, J. Chun, ContEst16S: An algorithm that identifies contaminated prokaryotic genomes using 16S RNA gene sequences, Int. J. Syst. Evol. Microbiol. 67 (2017) 2053–2057. https://doi.org/10.1099/ijsem.0.001872.

[25] C. Kerepesi, D. Bánky, V. Grolmusz, AmphoraNet: The webserver implementation of the AMPHORA2 metagenomic workflow suite, Gene. 533 (2014) 538–540. https://doi.org/10.1016/j.gene.2013.10.015.

[26] I.M.A. Chen, K. Chu, K. Palaniappan, A. Ratner, J. Huang, M. Huntemann, P. Hajek, S. Ritter, N. Varghese, R. Seshadri, S. Roux, T. Woyke, E.A. Eloe-Fadrosh, N.N. Ivanova, N.C. Kyrpides, The IMG/M data management and analysis system v.6.0: New tools and advanced capabilities, Nucleic Acids Res. 49 (2021) D751–D763. https://doi.org/10.1093/nar/gkaa939.

[27] N.F. Alikhan, N.K. Petty, N.L. Ben Zakour, S.A. Beatson, BLAST Ring Image Generator (BRIG): Simple prokaryote genome comparisons, BMC Genomics. 12 (2011) 1–10. https://doi.org/10.1186/1471-2164-12-402.

[28] T. Tatusova, M. Dicuccio, A. Badretdin, V. Chetvernin, E.P. Nawrocki, L. Zaslavsky, A. Lomsadze, K.D. Pruitt, M. Borodovsky, J. Ostell, NCBI prokaryotic genome annotation pipeline, Nucleic Acids Res. 44 (2016) 6614–6624. https://doi.org/10.1093/nar/gkw569.

[29] R.K. Aziz, D. Bartels, A. Best, M. DeJongh, T. Disz, R.A. Edwards, K. Formsma, S. Gerdes, E.M. Glass, M. Kubal, F. Meyer, G.J. Olsen, R. Olson, A.L. Osterman, R.A. Overbeek, L.K. McNeil, D. Paarmann, T. Paczian, B. Parrello, G.D. Pusch, C. Reich, R. Stevens, O. Vassieva, V. Vonstein, A. Wilke, O. Zagnitko, The RAST Server: Rapid annotations using subsystems technology, BMC Genomics. 9 (2008) 75. https://doi.org/10.1186/1471-2164-9-75.

[30] A. Carattoli, E. Zankari, A. Garciá-Fernández, M.V. Larsen, O. Lund, L. Villa, F.M. Aarestrup, H. Hasman, In Silico detection and typing of plasmids using plasmidfinder and plasmid multilocus sequence typing, Antimicrob. Agents Chemother. 58 (2014) 3895–3903. https://doi.org/10.1128/AAC.02412-14.

[31] Y. Zhou, Y. Liang, K.H. Lynch, J.J. Dennis, D.S. Wishart, PHAST: A Fast Phage Search Tool, Nucleic Acids Res. 39 (2011) W347–W352. https://doi.org/10.1093/nar/gkr485.

[32] S. Wu, Z. Zhu, L. Fu, B. Niu, W. Li, WebMGA: A customizable web server for fast metagenomic sequence analysis, BMC Genomics. 12 (2011) 444. https://doi.org/10.1186/1471-2164-12-444.

[33] S. El-Gebali, J. Mistry, A. Bateman, S.R. Eddy, A. Luciani, S.C. Potter, M. Qureshi, L.J. Richardson, G.A. Salazar, A. Smart, E.L.L. Sonnhammer, L. Hirsh, L. Paladin, D. Piovesan, S.C.E. Tosatto, R.D. Finn, The Pfam protein families database in 2019, Nucleic Acids Res. 47 (2019) D427–D432. https://doi.org/10.1093/nar/gky995.

[34] M. Barakat, P. Ortet, D.E. Whitworth, P2RP: A web-based framework for the identification and analysis of regulatory proteins in prokaryotic genomes, BMC Genomics. 14 (2013) 269. https://doi.org/10.1186/1471-2164-14-269.

[35] M.H. Saier, V.S. Reddy, B. V. Tsu, M.S. Ahmed, C. Li, G. Moreno-Hagelsieb, The Transporter Classification Database (TCDB): Recent advances, Nucleic Acids Res. 44 (2016) D372–D379. https://doi.org/10.1093/nar/gkv1103.

[36] B.P. Alcock, A.R. Raphenya, T.T.Y. Lau, K.K. Tsang, M. Bouchard, A. Edalatmand, W. Huynh, A.L. V. Nguyen, A.A. Cheng, S. Liu, S.Y. Min, A. Miroshnichenko, H.K. Tran, R.E. Werfalli, J.A. Nasir, M. Oloni, D.J. Speicher, A. Florescu, B. Singh, M. Faltyn, A. Hernandez-Koutoucheva, A.N. Sharma, E. Bordeleau, A.C. Pawlowski, H.L. Zubyk, D. Dooley, E. Griffiths, F. Maguire, G.L. Winsor, R.G. Beiko, F.S.L. Brinkman, W.W.L. Hsiao, G. V. Domselaar, A.G. McArthur, CARD 2020: Antibiotic resistome surveillance with the comprehensive antibiotic resistance database, Nucleic Acids Res. 48 (2020) D517–D525. https://doi.org/10.1093/nar/gkz935.

[37] A. de Jong, S.A.F.T. van Hijum, J.J.E. Bijlsma, J. Kok, O.P. Kuipers, BAGEL: A web-based bacteriocin genome mining tool, Nucleic Acids Res. 34 (2006) W273–W279. https://doi.org/10.1093/nar/gkl237.

[38] M.H. Medema, K. Blin, P. Cimermancic, V. De Jager, P. Zakrzewski, M.A. Fischbach, T. Weber, E. Takano, R. Breitling, AntiSMASH: Rapid identification, annotation and analysis of secondary metabolite biosynthesis gene clusters in bacterial and fungal genome sequences, Nucleic Acids Res. 39 (2011) W339–W346. https://doi.org/10.1093/nar/gkr466.

[39] Y. Yin, X. Mao, J. Yang, X. Chen, F. Mao, Y. Xu, DbCAN: A web resource for automated carbohydrate-active enzyme annotation, Nucleic Acids Res. 40 (2012) W445–W451. https://doi.org/10.1093/nar/gks479.

[40] V. Lombard, H. Golaconda Ramulu, E. Drula, P.M. Coutinho, B. Henrissat, The carbohydrate-active enzymes database (CAZy) in 2013, Nucleic Acids Res. 42 (2014) D490–D495. https://doi.org/10.1093/nar/gkt1178.

[41] M. Kanehisa, Y. Sato, KEGG Mapper for inferring cellular functions from protein sequences, Protein Sci. 29 (2020) 28–35. https://doi.org/10.1002/pro.3711.

[42] C. Bertelli, M.R. Laird, K.P. Williams, B.Y. Lau, G. Hoad, G.L. Winsor, F.S.L. Brinkman, IslandViewer 4: Expanded prediction of genomic islands for larger-scale datasets,Nucleic Acids Res. 45 (2017) W30–W35. https://doi.org/10.1093/nar/gkx343.

[43] D.M. Emms, S. Kelly, OrthoFinder: solving fundamental biases in whole genome comparisons dramatically improves orthogroup inference accuracy, Genome Biol. 16 (2015) 1–14. https://doi.org/10.1186/s13059-015-0721-2.

[44] M. Kearse, R. Moir, A. Wilson, S. Stones-Havas, M. Cheung, S. Sturrock, S. Buxton, A. Cooper, S. Markowitz, C. Duran, T. Thierer, B. Ashton, P. Meintjes, A. Drummond, Geneious Basic: An integrated and extendable desktop software platform for the organization and analysis of sequence data, Bioinformatics. 28 (2012) 1647– 1649. https://doi.org/10.1093/bioinformatics/bts199.

[45] M. Kim, H.S. Oh, S.C. Park, J. Chun, Towards a taxonomic coherence between average nucleotide identity and 16S rRNA gene sequence similarity for species demarcation of prokaryotes, Int. J. Syst. Evol. Microbiol. 64 (2014) 346–351. https://doi.org/10.1099/ijs.0.059774-0.

[46] R.E. Hungate, R.W. Dougherty, M.P. Bryant, R.M. Cello, Microbiological and physiological changes associated with acute indigestion in sheep, Cornell Vet. 42 (1952) 423–449. https://www.cabdirect.org/cabdirect/abstract/19531402099 (accessed August 25, 2020).

[47] F. Diez-Gonzalez, D.R. Bond, E. Jennings, J.B. Russell, Alternative schemes of butyrate production in Butyrivibrio fibrisolvens and their relationship to acetate utilization, lactate production, and phylogeny, Arch. Microbiol. 171 (1999) 324–330. https://doi.org/10.1007/s002030050717.

[48] A. Willems, M.D. Collins, Butyrivibrio, in: Bergey’s Man. Syst. Archaea Bact., Wiley, 2015: pp. 1–20. https://doi.org/10.1002/9781118960608.gbm00640.

[49] J.A. Patterson, S.C. Ricke, Effect of ethanol and methanol on growth of ruminal bacteria Selenomonas ruminantium and Butyrivibrio fibrisolvens, J. Environ. Sci. Heal. - Part B Pestic. Food Contam. Agric. Wastes. 50 (2015) 62–67. https://doi.org/10.1080/03601234.2015.965639.

[50] K.J. Shingfield, P. Kairenius, A. Ärölä, D. Paillard, S. Muetzel, S. Ahvenjärvi, A. Vanhatalo, P. Huhtanen, V. Toivonen, J.M. Griinari, R.J. Wallace, Dietary fish oil supplements modify ruminal biohydrogenation, alter the flow of fatty acids at the omasum, and induce changes in the ruminal butyrivibrio population in lactating cows, J. Nutr. 142 (2012) 1437–1448. https://doi.org/10.3945/jn.112.158576.

[51] J. Jeyanathan, M. Escobar, R.J. Wallace, V. Fievez, B. Vlaeminck, Biohydrogenation of 22:6n-3 by Butyrivibrio proteoclasticus P18, BMC Microbiol. 16 (2016) 104. https://doi.org/10.1186/s12866-016-0720-9.

[52] Y. Kobayashi, R.J. Forster, M. Alice Hefford, R.M. Teather, M. Wakita, K. Ohmiya, S. Hoshino, Analysis of the sequence of a new cryptic plasmid, pRJF2, from a rumen bacterium of the genus Butyrivibrio: Comparison with other Butyrivibrio plasmids and application in the development of a cloning vector, FEMS Microbiol. Lett. 130 (1995) 137–143. https://doi.org/10.1016/0378-1097(95)00193-9.

[53] J.P. Meier-Kolthoff, M. Göker, TYGS is an automated high-throughput platform for state-of-the-art genome-based taxonomy, Nat. Commun. 10 (2019) 1–10. https://doi.org/10.1038/s41467-019-10210-3.

[54] J.R. Hernáez, M.E.C. Cucchi, S. Cravero, M.C. Martinez, S. Gonzalez, A. Puebla, J. Dopazo, M. Farber, N. Paniego, M. Rivarola, The first complete genomic structure of Butyrivibrio fibrisolvens and its chromid, Microb. Genomics. 4 (2018) e000216. https://doi.org/10.1099/mgen.0.000216.

[55] N. Palevich, W.J. Kelly, S. Ganesh, J. Rakonjac, G.T. Attwood, Butyrivibrio hungatei MB2003 competes effectively for soluble sugars released by Butyrivibrio proteoclasticus B316 T during growth on xylan or pectin, Appl. Environ. Microbiol. 85 (2019). https://doi.org/10.1128/AEM.02056-18.

[56] B. Xin, F. Tao, Y. Wang, C. Gao, C. Ma, P. Xu, Genome sequence of Clostridium butyricum strain DSM 10702, a promising producer of biofuels and biochemicals, Genome Announc. 1 (2013). https://doi.org/10.1128/genomeA.00563-13.

[57] R. Seshadri, S.C. Leahy, G.T. Attwood, K.H. Teh, S.C. Lambie, A.L. Cookson, E.A. Eloe-Fadrosh, G.A. Pavlopoulos, M. Hadjithomas, N.J. Varghese, D. Paez-Espino, R. Perry, G. Henderson, C.J. Creevey, N. Terrapon, P. Lapebie, E. Drula, V. Lombard, E. Rubin, N.C. Kyrpides, B. Henrissat, T. Woyke, N.N. Ivanova, W.J. Kelly, N. Palevic, P.H. Janssen, R.S. Ronimus, S. Noel, P. Soni, K. Reilly, T. Atherly, C. Ziemer, A.D. Wright, S. Ishaq, M. Cotta, S. Thompson, K. Crosley, N. McKain, J.J. Wallace, H.J. Flint, J.C. Martin, R.J. Forster, R.J. Gruninger, T. McAllister, R. Gilbert, D.J. Ouwerkerk, A.J. Klieve, R. Al Jassim, S. Denman, C. McSweeney, C. Rosewarne, S. Koike, Y. Kobayashi, M. Mitsumori, T. Shinkai, S. Cravero, M. Cerón Cucchi, Cultivation and sequencing of rumen microbiome members from the Hungate1000 Collection, Nat. Biotechnol. 36 (2018) 359–367. https://doi.org/10.1038/nbt.4110.

[58] G. Henderson, F. Cox, S. Ganesh, A. Jonker, W. Young, P.H. Janssen, L. Abecia, E. Angarita, P. Aravena, G.N. Arenas, C. Ariza, G.T. Attwood, J.M. Avila, J. Avila-Stagno, A. Bannink, R. Barahona, M. Batistotti, M.F. Bertelsen, A. Brown-Kav, A.M. Carvajal, L. Cersosimo, A.V. Chaves, J. Church, N. Clipson, M.A. Cobos-Peralta, A.L. Cookson, S. Cravero, O.C. Carballo, K. Crosley, G. Cruz, M.C. Cucchi, R. De La Barra, A.B. De Menezes, E. Detmann, K. Dieho, J. Dijkstra, W.L.S. Dos Reis, M.E.R. Dugan, S.H. Ebrahimi, E. Eythórsdóttir, F.N. Fon, M. Fraga, F. Franco, C. Friedeman, N. Fukuma, D. Gagić, I. Gangnat, D.J. Grilli, L.L. Guan, V.H. Miri, E. Hernandez-Sanabria, A.X.I. Gomez, O.A. Isah, S. Ishaq, E. Jami, J. Jelincic, J. Kantanen, W.J. Kelly, S.H. Kim, A. Klieve, Y. Kobayashi, S. Koike, J. Kopecny, T.N. Kristensen, S.J. Krizsan, H. LaChance, M. Lachman, W.R. Lamberson, S. Lambie, J. Lassen, S.C. Leahy, S.S. Lee, F. Leiber, E. Lewis, B. Lin, R. Lira, P. Lund, E. Macipe, L.L. Mamuad, H.C. Mantovani, G.A. Marcoppido, C. Márquez, C. Martin, G. Martinez, M.E. Martinez, O.L. Mayorga, T.A. McAllister, C. McSweeney, L. Mestre, E. Minnee, M. Mitsumori, I. Mizrahi, I. Molina, A. Muenger, C. Munoz, B. Murovec, J. Newbold, V. Nsereko, M. O’Donovan, S. Okunade, B. O’Neill, S. Ospina, D. Ouwerkerk, D. Parra, L.G.R. Pereira, C. Pinares-Patino, P.B. Pope, M. Poulsen, M. Rodehutscord, T. Rodriguez, K. Saito, F. Sales, C. Sauer, K. Shingfield, N. Shoji, J. Simunek, Z. Stojanović-Radić, B. Stres, X. Sun, J. Swartz, Z.L. Tan, I. Tapio, T.M. Taxis, N. Tomkins, E. Ungerfeld, R. Valizadeh, P. Van Adrichem, J. Van Hamme, W. Van Hoven, G. Waghorn, R.J. Wallace, M. Wang, S.M. Waters, K. Keogh, M. Witzig, A.D.G. Wright, H. Yamano, T. Yan, D.R. Yanez-Ruiz, C.J. Yeoman, R. Zambrano, J. Zeitz, M. Zhou, H.W. Zhou, C.X. Zou, P. Zunino, Rumen microbial community composition varies with diet and host, but a core microbiome is found across a wide geographical range, Sci. Rep. 5 (2015) 1–15. https://doi.org/10.1038/srep14567.

[59] N. Palevich, P.H. Maclean, W.J. Kelly, S.C. Leahy, J. Rakonjac, G.T. Attwood, Complete genome sequence of the polysaccharide-degrading rumen bacterium Pseudobutyrivibrio xylanivorans MA3014 reveals an incomplete glycolytic pathway, Genome Biol. Evol. 12 (2020) 1566–1572. https://doi.org/10.1093/GBE/EVAA165.

[60] A.B. Boraston, D.N. Bolam, H.J. Gilbert, G.J. Davies, Carbohydrate-binding modules: Fine-tuning polysaccharide recognition, Biochem. J. 382 (2004) 769–781. https://doi.org/10.1042/BJ20040892.

[61] Y.S. Liu, Y. Zeng, Y. Luo, Q. Xu, M.E. Himmel, S.J. Smith, S.Y. Ding, Does the cellulose-binding module move on the cellulose surface?, Cellulose. 16 (2009) 587– 597. https://doi.org/10.1007/s10570-009-9306-0.

[62] H.J. Strobel, Pentose transport by the ruminal bacterium Butyrivibrio fibrisolvens, FEMS Microbiol. Lett. 122 (1994) 217–122. https://doi.org/10.1016/0378-1097(94)00324-6.

[63] M. Marounek, O. Petr, Fermentation of glucose and xylose in ruminal strains of Butyrivibrio fibrisolvens, Lett. Appl. Microbiol. 21 (1995) 272–276. https://doi.org/10.1111/j.1472-765X.1995.tb01058.x.

[64] T.J. Hackmann, J.L. Firkins, Electron transport phosphorylation in rumen butyrivibrios: Unprecedented ATP yield for glucose fermentation to butyrate, Front. Microbiol. 6 (2015) 622. https://doi.org/10.3389/fmicb.2015.00622.

[65] Y. Wang, Y. Tashiro, K. Sonomoto, Fermentative production of lactic acid from renewable materials: Recent achievements, prospects, and limits, J. Biosci. Bioeng. 119 (2015) 10–18. https://doi.org/10.1016/j.jbiosc.2014.06.003.

[66] M.F. Reid, C.A. Fewson, Molecular characterization of microbial alcohol dehydrogenases, Crit. Rev. Microbiol. 20 (1994) 13–56. https://doi.org/10.3109/10408419409113545.

[67] A. Joshi, G. Vasudevan, A. Engineer, S. Pore, S.S. Hivarkar, V.B. Lanjekar, P.K. Dhakephalkar, S.S. Dagar, Genomic Analysis of Actinomyces sp. Strain CtC72, a Novel Fibrolytic Anaerobic Bacterium Isolated from Cattle Rumen, Microbiol. Biotechnol. Lett. 46 (2018) 59–67. https://doi.org/10.4014/mbl.1712.12005.

[68] X. Guo, C. Cao, Y. Wang, C. Li, M. Wu, Y. Chen, C. Zhang, H. Pei, D. Xiao, Effect of the inactivation of lactate dehydrogenase, ethanol dehydrogenase, and phosphotransacetylase on 2,3-butanediol production in Klebsiella pneumoniae strain, Biotechnol. Biofuels. 7 (2014) 1–11. https://doi.org/10.1186/1754-6834-7-44.

[69] D.K. Kim, C. Rathnasingh, H. Song, H.J. Lee, D. Seung, Y.K. Chang, Metabolic engineering of a novel Klebsiella oxytoca strain for enhanced 2,3-butanediol production, J. Biosci. Bioeng. 116 (2013) 186–192. https://doi.org/10.1016/j.jbiosc.2013.02.021.

[70] J. Kopečný, M. Zorec, J. Mrázek, Y. Kobayashi, R. Marinšek-Logar, Butyrivibrio hungatei sp. nov. and Pseudobutyrivibrio xylanivorans sp. nov., butyrate-producing bacteria from the rumen, Int. J. Syst. Evol. Microbiol. 53 (2003) 201–209. https://doi.org/10.1099/ijs.0.02345-0.

[71] L. Molina G L. Giraldo V D. Polanco EL. Gutiérrez B, Cellulolytic and Butyrivibrio fibrisolvens bacteria population density, after supplementing fodder diets (Pennisetum clandestinum), Rev. MVZ Córdoba. (2015) 4947–4961. https://doi.org/10.21897/rmvz.10.

[72] W. Shao, Q. Wang, P.F. Rupani, S. Krishnan, F. Ahmad, S. Rezania, M.A. Rashid, C. Sha, M.F. Md Din, Biohydrogen production via thermophilic fermentation: A prospective application of Thermotoga species, Energy. 197 (2020) 117199. https://doi.org/10.1016/j.energy.2020.117199.

[73] M. Calusinska, T. Happe, B. Joris, A. Wilmotte, The surprising diversity of clostridial hydrogenases: A comparative genomic perspective, Microbiology. 156 (2010) 1575– 1588. https://doi.org/10.1099/mic.0.032771-0.

[74] P.J. Weimer, Manipulating Ruminal Fermentation: A Microbial Ecological Perspective 1, J. Anim. Sci. 76 (1998) 3114–3122. https://doi.org/10.2527/1998.76123114x.

[75] S.S. Dagar, S. Kumar, P. Mudgil, A.K. Puniya, Comparative evaluation of lignocellulolytic activities of filamentous cultures of monocentric and polycentric anaerobic fungi, Anaerobe. 50 (2018) 76–79. https://doi.org/10.1016/j.anaerobe.2018.02.004.

[76] J.J. Bond, J.C. Dunne, F.Y.S. Kwan, D. Li, K. Zhang, S.C. Leahy, W.J. Kelly, G.T. Attwood, T.W. Jordan, Carbohydrate transporting membrane proteins of the rumen bacterium, Butyrivibrio proteoclasticus, J. Proteomics. 75 (2012) 3138–3144. https://doi.org/10.1016/j.jprot.2011.12.013.

[77] B. Wang, M. Dukarevich, E.I. Sun, M.R. Yen, M.H. Saier, Membrane porters of ATP-binding cassette transport systems are polyphyletic, J. Membr. Biol. 231 (2009) 1–10. https://doi.org/10.1007/s00232-009-9200-6.

[78] J.S. Chen, V. Reddy, J.H. Chen, M.A. Shlykov, W.H. Zheng, J. Cho, M.R. Yen, M.H. Saier, Phylogenetic characterization of transport protein superfamilies: Superiority of SuperfamilyTree programs over those based on multiple alignments, J. Mol. Microbiol. Biotechnol. 21 (2012) 83–96. https://doi.org/10.1159/000334611.

[79] Y. Gbelska, J.J. Krijger, K.D. Breunig, Evolution of gene families: The multidrug resistance transporter genes in five related yeast species, FEMS Yeast Res. 6 (2006) 345–355. https://doi.org/10.1111/j.1567-1364.2006.00058.x.

[80] D.L. Jack, N.M. Yang, M.H. Saier, The drug/metabolite transporter superfamily, Eur. J. Biochem. 268 (2001) 3620–3639. https://doi.org/10.1046/j.1432-1327.2001.02265.x.

[81] K. Wallden, A. Rivera-Calzada, G. Waksman, Type IV secretion systems: Versatility and diversity in function, Cell. Microbiol. 12 (2010) 1203–1212. https://doi.org/10.1111/j.1462-5822.2010.01499.x.

[82] M.L. Kalmokoff, F. Bartlett, R.M. Teather, Are ruminal bacteria armed with bacteriocins?, J. Dairy Sci. 79 (1996) 2297–2306. https://doi.org/10.3168/jds.S0022-0302(96)76608-0.

[83] M.L. Kalmokoff, R.M. Teather, Isolation and characterization of a bacteriocin (butyrivibriocin AR10) from the ruminal anaerobe Butyrivibrio fibrisolvens AR10: Evidence in support of the widespread occurrence of bacteriocin-like activity among ruminal isolates of B. fibrisolvens, Appl. Environ. Microbiol. 63 (1997) 394–402. https://doi.org/10.1128/aem.63.2.394-402.1997.

[84] T.R. Klaenhammer, Genetics of bacteriocins produced by lactic acid bacteria, FEMS Microbiol. Rev. 12 (1993) 39–85. https://doi.org/10.1111/j.1574-6976.1993.tb00012.x.

[85] A. Lauková, M. Mareková, Antimicrobial spectrum of bacteriocin-like substances produced by rumen staphylococci, Folia Microbiol. (Praha). 38 (1993) 74–76. https://doi.org/10.1007/BF02814554.

[86] M.F. Whitford, M.A. McPherson, R.J. Forster, R.M. Teather, Identificafion of bacteriocin-like inhibitors from rumen Streptococcus spp. and isolation and characterization of bovicin 255, Appl. Environ. Microbiol. 67 (2001) 569–574. https://doi.org/10.1128/AEM.67.2.569-574.2001.

[87] E.L. Ongey, L. Santolin, S. Waldburger, L. Adrian, S.L. Riedel, P. Neubauer, Bioprocess Development for Lantibiotic Ruminococcin-A Production in Escherichia coli and Kinetic Insights Into LanM Enzymes Catalysis, Front. Microbiol. 10 (2019) 2133. https://doi.org/10.3389/fmicb.2019.02133.

[88] Y.N.V. Sabino, M.F. Santana, L.B. Oyama, F.G. Santos, A.J.S. Moreira, S.A. Huws, H.C. Mantovani, Characterization of antibiotic resistance genes in the species of the rumen microbiota, Nat. Commun. 10 (2019) 1–11. https://doi.org/10.1038/s41467-019-13118-0.

[89] S.M. Friedman, T. Lu, K. Drlica, Mutation in the DNA gyrase A gene of Escherichia coli that expands the quinolone resistance-determining region, Antimicrob. Agents Chemother. 45 (2001) 2378–2380. https://doi.org/10.1128/AAC.45.8.2378-2380.2001.

[90] A.L. Cooper, A.J. Low, A.G. Koziol, M.C. Thomas, D. Leclair, S. Tamber, A. Wong, B.W. Blais, C.D. Carrillo, Systematic Evaluation of Whole Genome Sequence-Based Predictions of Salmonella Serotype and Antimicrobial Resistance, Front. Microbiol. 11 (2020) 549. https://doi.org/10.3389/fmicb.2020.00549.

[91] R.B. Hespell, K. Kato, J.W. Costerton, Characterization of the cell wall of Butyrivibrio species, Can. J. Microbiol. 39 (1993) 912–921. https://doi.org/10.1139/m93-139.

[92] K.P. Scott, C.M. Melville, T.M. Barbosa, H.J. Flint, Occurrence of the new tetracycline resistance gene tet(W) in bacteria from the human gut, Antimicrob. Agents Chemother. 44 (2000) 775–777. https://doi.org/10.1128/AAC.44.3.775-777.2000.

[93] P. Spigaglia, F. Barbanti, P. Mastrantonio, Horizontal transfer of erythromycin resistance from Clostridium difficile to Butyrivibrio fibrisolvens, Antimicrob. Agents Chemother. 49 (2005) 5142–5145. https://doi.org/10.1128/AAC.49.12.5142-5145.2005.

[94] G.A. Preidis, C. Hill, R.L. Guerrant, B.S. Ramakrishna, G.W. Tannock, J. Versalovic, Probiotics, enteric and diarrheal diseases, and global health, Gastroenterology. 140 (2011) 8–14. https://doi.org/10.1053/j.gastro.2010.11.010.

[95] J. Franco-Lopez, M. Duplessis, A. Bui, C. Reymond, W. Poisson, L. Blais, J. Chong, R. Gervais, D.E. Rico, R.I. Cue, C.L. Girard, J. Ronholm, Correlations between the Composition of the Bovine Microbiota and Vitamin B 12 Abundance, MSystems. 5 (2020). https://doi.org/10.1128/msystems.00107-20.

[96] C. Harfoot, G. Hazlewood, Lipid metabolism in the rumen. In “The rumen microbial ecosystem”.(Ed. PN Hobson), (1997) 382–426.

[97] B. Yang, H. Chen, F. Tian, J. Zhao, Z. Gu, H. Zhang, Y.Q. Chen, W. Chen, Complete genome sequence of Lactobacillus plantarum ZS2058, a probiotic strain with high conjugated linoleic acid production ability, J. Biotechnol. 214 (2015) 212–213. https://doi.org/10.1016/j.jbiotec.2015.09.036.

[98] J.L.C.M. Van De Vossenberg, K.N. Joblin, Biohydrogenation of C18 unsaturated fatty acids to stearic acid by a strain of Butyrivibrio hungatei from the bovine rumen, Lett. Appl. Microbiol. 37 (2003) 424–428. https://doi.org/10.1046/j.1472-765X.2003.01421.x.

[99] B. Drogue, H. Doré, S. Borland, F. Wisniewski-Dyé, C. Prigent-Combaret, Which specificity in cooperation between phytostimulating rhizobacteria and plants?, Res. Microbiol. 163 (2012) 500–510. https://doi.org/10.1016/j.resmic.2012.08.006.

[100] M.M. Galicia-Jiménez, R. Rojas-Herrera, C. Sandoval-Castro, S.E. Murialdo, H. Magaña-Sevilla, Chemotactic responses of the rumen bacterial community towards the daidzein flavonoid, Livest. Sci. 167 (2014) 121–125. https://doi.org/10.1016/j.livsci.2014.05.022.

[101] S.E. Murialdo, G.H. Sendra, L.I. Passoni, R. Arizaga, J.F. Gonzalez, H. Rabal, M. Trivi, Analysis of bacterial chemotactic response using dynamic laser speckle, J. Biomed. Opt. 14 (2009) 064015–064016. https://doi.org/10.1117/1.3262608.

[102] V. Sourjik, Receptor clustering and signal processing in E. coli chemotaxis, Trends Microbiol. 12 (2004) 569–576. https://doi.org/10.1016/j.tim.2004.10.003.

[103] H.L. Diaz, S.K.R. Karnati, M.A. Lyons, B.A. Dehority, J.L. Firkins, Chemotaxis toward carbohydrates and peptides by mixed ruminal protozoa when fed, fasted, or incubated with polyunsaturated fatty acids, J. Dairy Sci. 97 (2014) 2231–2243. https://doi.org/10.3168/jds.2013-7428.

[104] C.G. Orpin, L. Bountiff, Zoospore chemotaxis in the rumen phycomycete Neocallimastix frontalis, J. Gen. Microbiol. 104 (1978) 113–122. https://doi.org/10.1099/00221287-104-1-113.

[105] D.A. Wubah, D.S.H. Kim, Chemoattraction of anaerobic ruminal fungi zoospores to selected phenolic acids, Microbiol. Res. 151 (1996) 257–262. https://doi.org/10.1016/S0944-5013(96)80022-X.

[106] Palevich, Nikola, Comparative genomics of Butyrivibrio and Pseudobutyrivibrio from the rumen, 2016. https://mro.massey.ac.nz/handle/10179/9992 (accessed September 10, 2020).

[107] R.G. Martin, J.L. Rosner, The AraC transcriptional activators, Curr. Opin. Microbiol. 4 (2001) 132–137. https://doi.org/10.1016/S1369-5274(00)00178-8.

[108] K. Wu, H. Xu, Y. Zheng, L. Wang, X. Zhang, Y. Yin, CpsR, a GntR family regulator, transcriptionally regulates capsular polysaccharide biosynthesis and governs bacterial virulence in Streptococcus pneumoniae, Sci. Rep. 6 (2016) 29255. https://doi.org/10.1038/srep29255.

[109] S.P. Wilkinson, A. Grove, Ligand-responsive transcriptional regulation by members of the MarR family of winged helix proteins, Curr. Issues Mol. Biol. 8 (2006) 51–62. https://doi.org/10.21775/cimb.008.051.

[110] S.S. Gupta, B.N. Borin, T.L. Cover, A.M. Krezel, Structural analysis of the DNA-binding domain of the Helicobacter pylori response regulator ArsR, J. Biol. Chem. 284 (2009) 6536–6545. https://doi.org/10.1074/jbc.M804592200.

[111] P. Oliveira, P. Lindblad, LexA, a transcription regulator binding in the promoter region of the bidirectional hydrogenase in the cyanobacterium Synechocystis sp. PCC 6803, FEMS Microbiol. Lett. 251 (2005) 59–66. https://doi.org/10.1016/j.femsle.2005.07.024.

[112] R.D. Rolfe, D.J. Hentges, J.T. Barrett, B.J. Campbell, Oxygen tolerance of human intestinal anaerobes, Am. J. Clin. Nutr. 30 (1977) 1762–1769. https://doi.org/10.1093/ajcn/30.11.1762.

[113] E.R. Rocha, T. Selby, J.P. Coleman, C. Jeffrey Smith, Oxidative stress response in an anaerobe, Bacteroides fragilis: A role for catalase in protection against hydrogen peroxide, J. Bacteriol. 178 (1996) 6895–6903. https://doi.org/10.1128/jb.178.23.6895-6903.1996.

[114] J.P. Schumann, D.T. Jones, D.R. Woods, Induction of proteins during phage reactivation induced by UV irradiation, oxygen and peroxide in Bacteroides fragilis, FEMS Microbiol. Lett. 23 (1984) 131–135. https://doi.org/10.1111/j.1574-6968.1984.tb01048.x.

[115] M. Juhas, J.R. Van Der Meer, M. Gaillard, R.M. Harding, D.W. Hood, D.W. Crook, Genomic islands: Tools of bacterial horizontal gene transfer and evolution, FEMS Microbiol. Rev. 33 (2009) 376–393. https://doi.org/10.1111/j.1574-6976.2008.00136.x.

[116] A.B. Kav, G. Sasson, E. Jami, A. Doron-Faigenboim, I. Benhar, I. Mizrahi, Insights into the bovine rumen plasmidome, Proc. Natl. Acad. Sci. U. S. A. 109 (2012) 5452– 5457. https://doi.org/10.1073/pnas.1116410109.

