## Supplementary file 1 for "Genomic architecture of three newly isolated unclassified *Butyrivibrio* species elucidate their potential role in the rumen ecosystem"

Report Date: 03/31/2020
Organism: Butyrivibrio sp. sp3-iso-ab22 CB0-8

### Summary

An assembled genome for Butyrivibrio sp. sp3-iso-ab22 CB0-8 was submitted to the comprehensive genome analysis service at PATRIC[1]. Based on the annotation statistics and a comparison to other genomes in PATRIC within this same species, this genome appears to be of Good quality. Details of the analysis, including genes of interest (Specialty Genes), a functional categorization (Subsystems), and a phylogenetic tree (Phylogenetic Analysis) are provided below.

### Genome Assembly

An assembled genome was submitted to the Comprehensive Genome Analysis service. This assembled genome had 34 contigs, with the total length of 3,699,710 bp and an average G+C content of 43.70% (Table 1).

Table 1. Assembly Details

| | |
|---|---|
| Contigs | 34 |
| GC Content | 43.70 |
| Plasmids | 0 |
| Contig L50 | 3 |
| Genome Length | 3,699,710 bp |
| Contig N50 | 332,484 |
| Chromosomes | 0 |

### Genome Annotation

The Butyrivibrio sp. sp3-iso-ab22 CB0-8 genome was annotated using RAST tool kit (RASTtk)[2] and assigned a unique genome identifier of 865693.3. This genome is in the superkingdom and was annotated using genetic code 11. The taxonomy of this genome is:

> cellular organisms > Bacteria > Terrabacteria group > Firmicutes > Clostridia > Clostridiales > Lachnospiraceae > Butyrivibrio > unclassified Butyrivibrio > Butyrivibrio sp. sp3-iso-ab22

This genome has 3,474 protein coding sequences (CDS), 46 transfer RNA (tRNA) genes, and 4 ribosomal RNA (rRNA) genes. The annotated features are summarized in Table 2.

Table 2. Annotated Genome Features

| | |
|---|---|
| CDS | 3,474 |
| tRNA | 46 |
| rRNA | 4 |
| Partial CDS | 0 |
| Miscellaneous RNA | 0 |
| Repeat Regions | 0 |
| Job ID | annotation_9206 |
| Job Started | March 31st 2020, 6:57:52am |
| Job Completed | March 31st 2020, 7:08:26am |
| Total Time | 10 minutes and 34 seconds |

The annotation included 1,415 hypothetical proteins and 2,059 proteins with functional assignments (Table 3). The proteins with functional assignments included 704 proteins with Enzyme Commission (EC) numbers[3], 577 with Gene Ontology (GO) assignments[4], and 511 proteins that were mapped to KEGG pathways[5]. PATRIC annotation includes two types of protein families[6], and this genome has 2,534 proteins that belong to the genus-specific protein families (PLFams) for , and 2,667 proteins that belong to the cross-genus protein families (PGFams).

Table 3. Protein Features

| | |
|---|---|
| Hypothetical proteins | 1,415 |
| Proteins with functional assignments | 2,059 |
| Proteins with EC number assignments | 704 |
| Proteins with GO assignments | 577 |
| Proteins with Pathway assignments | 511 |
| Proteins with PATRIC genus-specific family (PLfam) assignments | 2,534 |
| Proteins with PATRIC cross-genus family (PGfam) assignments | 2,667 |

A circular graphical display of the distribution of the genome annotations is provided (Figure 1). This includes, from outer to inner rings, the contigs, CDS on the forward strand, CDS on the reverse strand, RNA genes, CDS with homology to known antimicrobial resistance genes, CDS with homology to know virulence factors, GC content and GC skew. The colors of the CDS on the forward and reverse strand indicate the subsystem that these genes belong to (see Subsystems below).

Figure 1

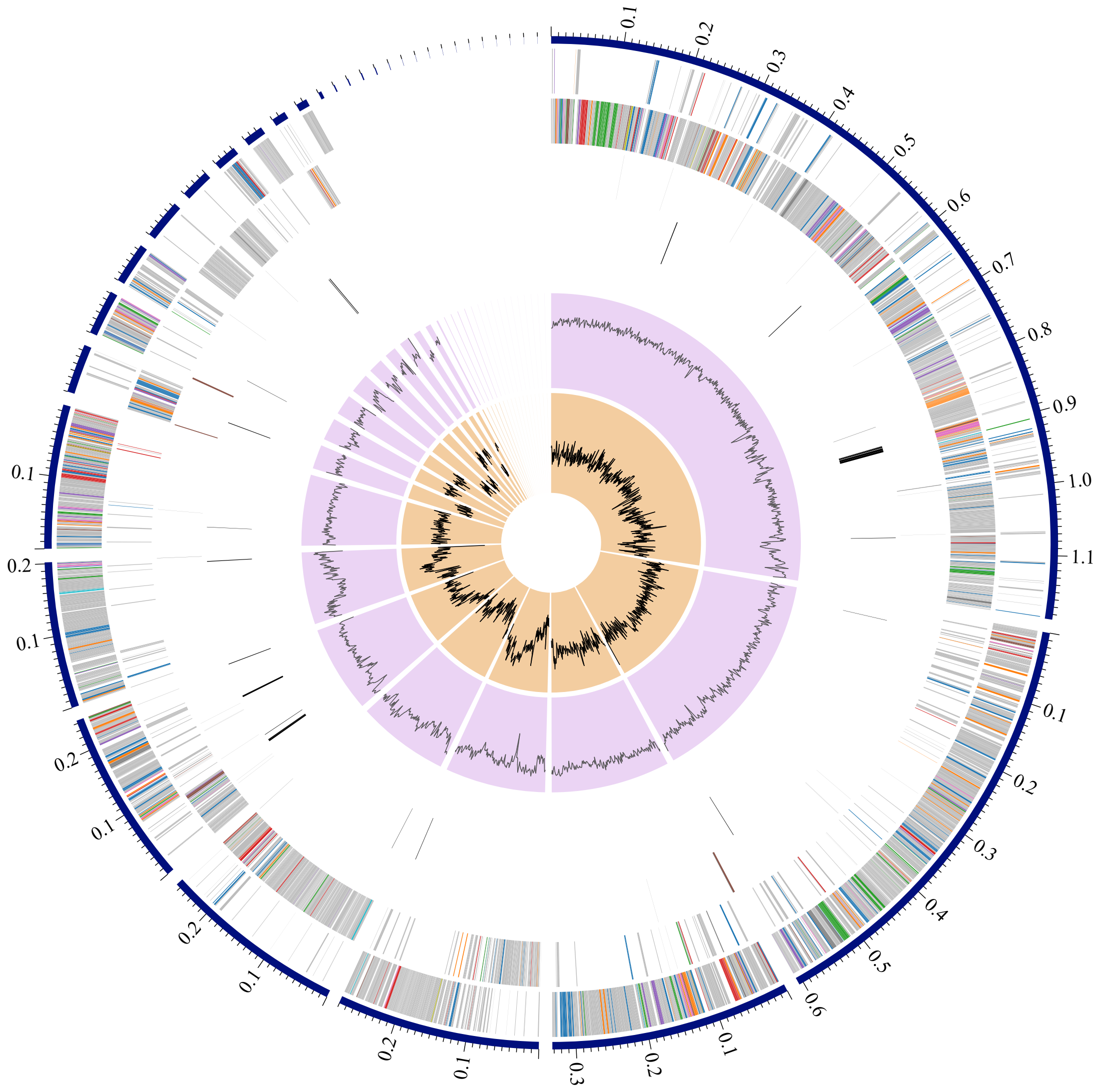

### Subsystem Analysis

A subsystem is a set of proteins that together implement a specific biological process or structural complex[7] and PATRIC annotation includes an analysis of the subsystems unique to each genome. An overview of the subsystems for this genome is provided in Figure 2.

Figure 2

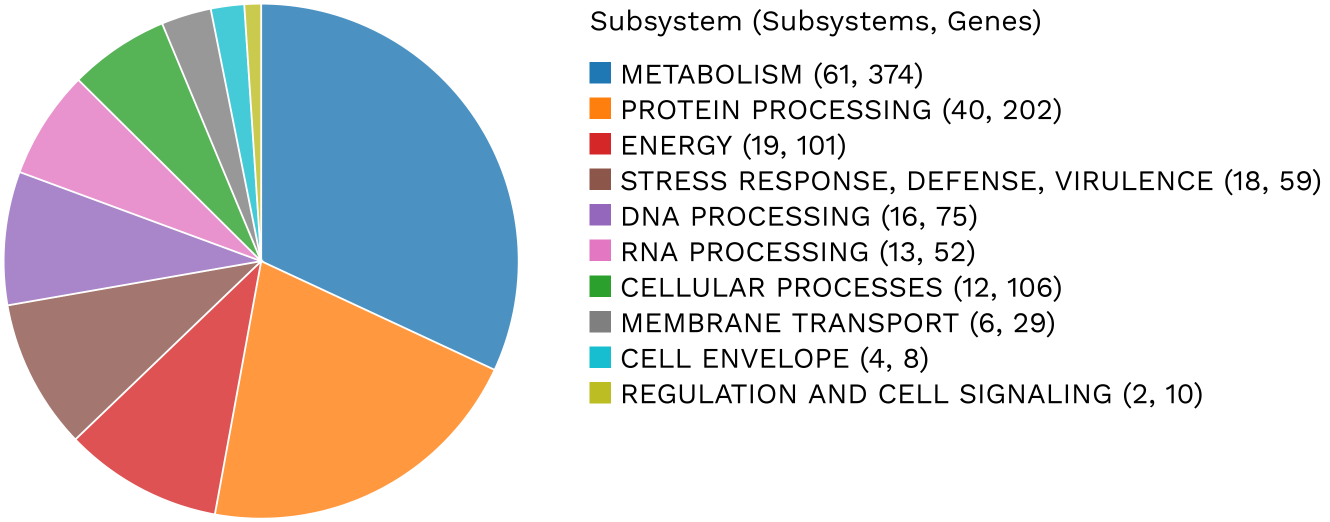

### Specialty Genes

Many of the genes annotated in have homology to known transporters[8], virulence factors[9][10], drug targets[11][12], and antibiotic resistance genes[13]. The number of genes and the specific source database where homology was found is provided (Table 4).

Table 4. Specialty Genes

| | Source | Genes |
|---|---|---|
| Antibiotic Resistance | PATRIC | 24 |
| Transporter | TCDB | 1 |

### Antimicrobial Resistance Genes

The Genome Annotation Service in PATRIC uses k-mer-based AMR genes detection method, which utilizes PATRIC's curated collection of representative AMR gene sequence variants[1] and assigns to each AMR gene functional annotation, broad mechanism of antibiotic resistance, drug class and, in some cases, specific antibiotic it confers resistance to. Please note, that the presence of AMR-related genes (even full length) in a given genome does not directly imply antibiotic resistant phenotype. It is important to consider specific AMR mechanisms and especially the absence/presence of SNP mutations conveying resistance. A summary of the AMR genes annotated in this genome and corresponding AMR mechanism is provided in Table 5.

Table 5. Antimicrobial Resistance Genes

| AMR Mechanism | Genes |
|---|---|
| Antibiotic inactivation enzyme | NimB |
| Antibiotic target in susceptible species | Alr, Ddl, dxr, EF-G, EF-Tu, folA, Dfr, gyrA, gyrB, Iso-tRNA, kasA, MurA, rho, rpoB, rpoC, S10p, S12p |
| Antibiotic target replacement protein | FabK |
| Gene conferring resistance via absence | gidB |
| Protein altering cell wall charge conferring antibiotic resistance | GdpD, PgsA |

### Phylogenetic Analysis

The National Center for Biotechnology Information (NCBI) staff manually select and categorize reference and representative genomes, which they consider to be of high quality and importance to the research community. PATRIC provides the reference and representative genomes, and includes them in the phylogenetic analysis that is part of the Comprehensive Genome Analysis report. The closest reference and representative genomes to were identified by Mash/MinHash[15]. PATRIC global protein families (PGFams)[6] were selected from these genomes to determine the phylogenetic placement of this genome. The protein sequences from these families were aligned with MUSCLE[17], and the nucleotides for each of those sequences were mapped to the protein alignment. The joint set of amino acid and nucleotide alignments were concatenated into a data matrix, and RaxML[18] was used to analyze this matrix, with fast bootstrapping[19] was used to generate the support values in the tree (Figure 3).

Figure 3

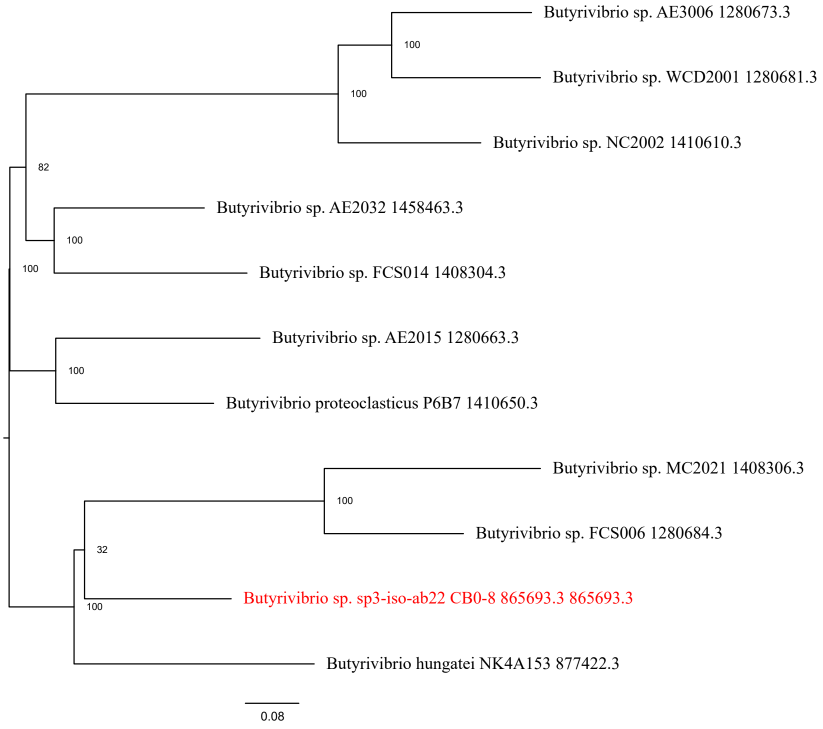


### References


1. Wattam AR, Davis JJ, Assaf R, Boisvert S, Brettin T, Bun C, Conrad N, Dietrich EM, Disz T, Gabbard JL, et al. 2017. Improvements to PATRIC, the all-bacterial Bioinformatics Database and Analysis Resource Center. Nucleic Acids Res 45:D535-D542.

2. Brettin T, Davis JJ, Disz T, Edwards RA, Gerdes S, Olsen GJ, Olson R, Overbeek R, Parrello B, Pusch GD, et al. 2015. RASTtk: a modular and extensible implementation of the RAST algorithm for building custom annotation pipelines and annotating batches of genomes. Sci Rep 5:8365.

3. Schomburg I, Chang A, Ebeling C, Gremse M, Heldt C, Huhn G, Schomburg D. 2004. BRENDA, the enzyme database: updates and major new developments. Nucleic Acids Res 32:D431-D433.

4. Ashburner M, Ball CA, Blake JA, Botstein D, Butler H, Cherry JM, Davis AP, Dolinski K, Dwight SS, Eppig JT. 2000. Gene Ontology: tool for the unification of biology. Nature genetics 25:25.

5. Kanehisa M, Sato Y, Kawashima M, Furumichi M, Tanabe M. 2016. KEGG as a reference resource for gene and protein annotation. Nucleic Acids Res 44:D457-462.

6. Davis JJ, Gerdes S, Olsen GJ, Olson R, Pusch GD, Shukla M, Vonstein V, Wattam AR, Yoo H. 2016. PATtyFams: Protein Families for the Microbial Genomes in the PATRIC Database. Front Microbiol 7:118.

7. Overbeek R, Begley T, Butler RM, Choudhuri JV, Chuang H-Y, Cohoon M, de Crécy-Lagard V, Diaz N, Disz T, Edwards R. 2005. The subsystems approach to genome annotation and its use in the project to annotate 1000 genomes. Nucleic Acids Res 33:5691-5702.

8. Saier Jr MH, Reddy VS, Tsu BV, Ahmed MS, Li C, Moreno-Hagelsieb G. 2015. The transporter classification database (TCDB): recent advances. Nucleic Acids Res 44:D372-D379.

9. Mao C, Abraham D, Wattam AR, Wilson MJ, Shukla M, Yoo HS, Sobral BW. 2015. Curation, integration and visualization of bacterial virulence factors in PATRIC. Bioinformatics 31:252-258.

10. Chen L, Zheng D, Liu B, Yang J, Jin Q. 2016. VFDB 2016: hierarchical and refined dataset for big data analysis-10 years on. Nucleic Acids Res 44:D694-D697.

11. Zhu F, Han B, Kumar P, Liu X, Ma X, Wei X, Huang L, Guo Y, Han L, Zheng C. 2009. Update of TTD: therapeutic target database. Nucleic Acids Res 38:D787-D791.

12. Law V, Knox C, Djoumbou Y, Jewison T, Guo AC, Liu Y, Maciejewski A, Arndt D, Wilson M, Neveu V, et al. 2014. DrugBank 4.0: shedding new light on drug metabolism. Nucleic Acids Res 42:D1091-1097.

13. McArthur AG, Waglechner N, Nizam F, Yan A, Azad MA, Baylay AJ, Bhullar K, Canova MJ, De Pascale G, Ejim L. 2013. The comprehensive antibiotic resistance database. Antimicrobial agents and chemotherapy 57:3348-3357.

14. Wattam AR, Davis JJ, Assaf R, Boisvert S, Brettin T, Bun C, Conrad N, Dietrich EM, Disz T, Gabbard JL, et al. 2017. Improvements to PATRIC, the all-bacterial Bioinformatics Database and Analysis Resource Center. Nucleic Acids Res 45:D535-D542.

15. Ondov BD, Treangen TJ, Melsted P, Mallonee AB, Bergman NH, Koren S, Phillippy AM. 2016. Mash: fast genome and metagenome distance estimation using MinHash. Genome biology 17:132.

16. Davis JJ, Gerdes S, Olsen GJ, Olson R, Pusch GD, Shukla M, Vonstein V, Wattam AR, Yoo H. 2016. PATtyFams: Protein Families for the Microbial Genomes in the PATRIC Database. Front Microbiol 7:118.

17. Edgar RC. 2004. MUSCLE: multiple sequence alignment with high accuracy and high throughput. Nucleic Acids Res 32:1792-1797.

18. Stamatakis A. 2014. RAxML version 8: a tool for phylogenetic analysis and post-analysis of large phylogenies. Bioinformatics 30:1312-1313.

19. Stamatakis A, Hoover P, Rougemont J. 2008. A rapid bootstrap algorithm for the RAxML web servers. Systematic biology 57:758-771.

Report Date: 03/31/2020
Organism: Butyrivibrio sp. sp3-iso-ab22 CB500-5

### Summary

An assembled genome for Butyrivibrio sp. sp3-iso-ab22 CB500-5 was submitted to the comprehensive genome analysis service at PATRIC[1]. Based on the annotation statistics and a comparison to other genomes in PATRIC within this same species, this genome appears to be of Good quality. Details of the analysis, including genes of interest (Specialty Genes), a functional categorization (Subsystems), and a phylogenetic tree (Phylogenetic Analysis) are provided below.

### Genome Assembly

An assembled genome was submitted to the Comprehensive Genome Analysis service. This assembled genome had 39 contigs, with the total length of 3,334,106 bp and an average G+C content of 42.37% (Table 1).

Table 1. Assembly Details

| | |
|---|---|
| Contigs | 39 |
| GC Content | 42.37 |
| Plasmids | 0 |
| Contig L50 | 4 |
| Genome Length | 3,334,106 bp |
| Contig N50 | 299,830 |
| Chromosomes | 0 |

### Genome Annotation

The Butyrivibrio sp. sp3-iso-ab22 CB500-5 genome was annotated using RAST tool kit (RASTtk)[2] and assigned a unique genome identifier of 865693.4. This genome is in the superkingdom and was annotated using genetic code 11. The taxonomy of this genome is:

> cellular organisms > Bacteria > Terrabacteria group > Firmicutes > Clostridia > Clostridiales > Lachnospiraceae > Butyrivibrio > unclassified Butyrivibrio > Butyrivibrio sp. sp3-iso-ab22

This genome has 3,093 protein coding sequences (CDS), 45 transfer RNA (tRNA) genes, and 4 ribosomal RNA (rRNA) genes. The annotated features are summarized in Table 2.

Table 2. Annotated Genome Features

| | |
|---|---|
| CDS | 3,093 |
| tRNA | 45 |
| rRNA | 4 |
| Partial CDS | 0 |
| Miscellaneous RNA | 0 |
| Repeat Regions | 0 |
| Job ID | annotation_9838 |
| Job Started | March 31st 2020, 6:59:08am |
| Job Completed | March 31st 2020, 7:09:52am |
| Total Time | 10 minutes and 44 seconds |

The annotation included 1,206 hypothetical proteins and 1,887 proteins with functional assignments (Table 3). The proteins with functional assignments included 657 proteins with Enzyme Commission (EC) numbers[3], 553 with Gene Ontology (GO) assignments[4], and 502 proteins that were mapped to KEGG pathways[5]. PATRIC annotation includes two types of protein families[6], and this genome has 2,357 proteins that belong to the genus-specific protein families (PLFams) for , and 2,406 proteins that belong to the cross-genus protein families (PGFams).

Table 3. Protein Features

| | |
|---|---|
| Hypothetical proteins | 1,206 |
| Proteins with functional assignments | 1,887 |
| Proteins with EC number assignments | 657 |
| Proteins with GO assignments | 553 |
| Proteins with Pathway assignments | 502 |
| Proteins with PATRIC genus-specific family (PLfam) assignments | 2,357 |
| Proteins with PATRIC cross-genus family (PGfam) assignments | 2,406 |

A circular graphical display of the distribution of the genome annotations is provided (Figure 1). This includes, from outer to inner rings, the contigs, CDS on the forward strand, CDS on the reverse strand, RNA genes, CDS with homology to known antimicrobial resistance genes, CDS with homology to know virulence factors, GC content and GC skew. The colors of the CDS on the forward and reverse strand indicate the subsystem that these genes belong to (see Subsystems below).

Figure 1

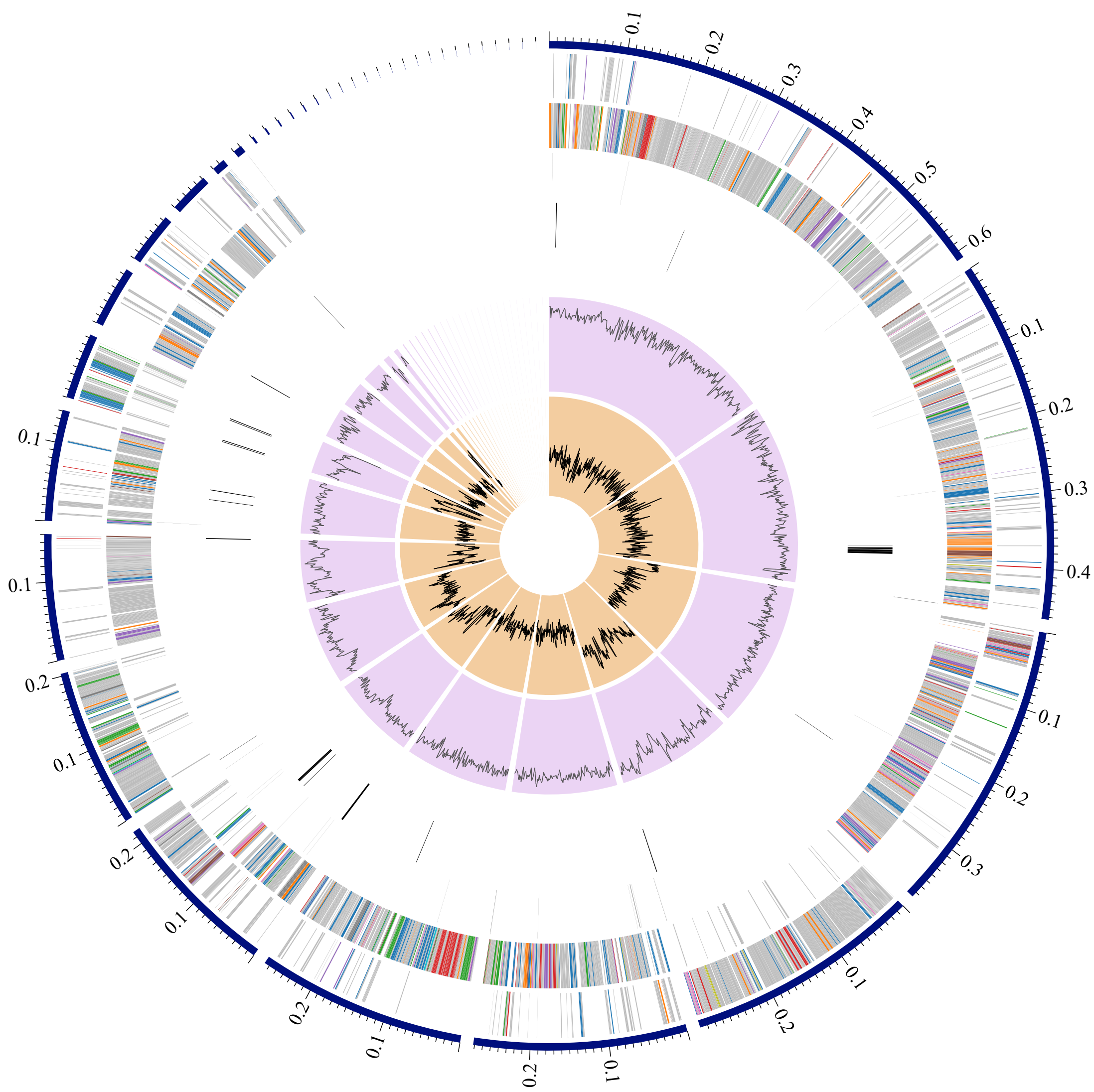

### Subsystem Analysis

A subsystem is a set of proteins that together implement a specific biological process or structural complex[7] and PATRIC annotation includes an analysis of the subsystems unique to each genome. An overview of the subsystems for this genome is provided in Figure 2.

Figure 2

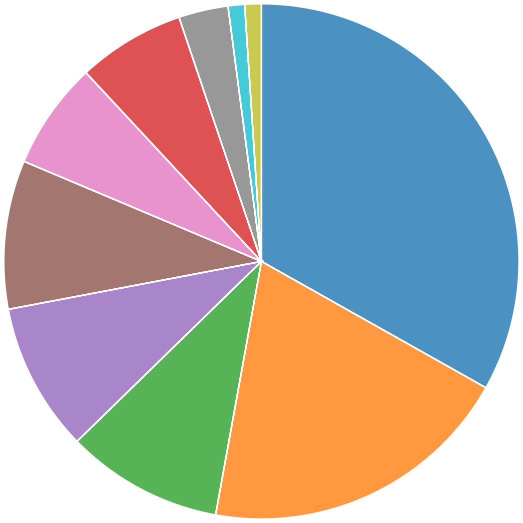

Subsystem (Subsystems, Genes)

- METABOLISM (64, 413)
- PROTEIN PROCESSING (38, 194)
- ENERGY (19, 115)
- DNA PROCESSING (18, 81)
- STRESS RESPONSE, DEFENSE, VIRULENCE (18, 57)
- RNA PROCESSING (13, 51)
- CELLULAR PROCESSES (13, 106)
- MEMBRANE TRANSPORT (6, 25)
- REGULATION AND CELL SIGNALING (2, 7)
- CELL ENVELOPE (2, 8)

### Specialty Genes

Many of the genes annotated in have homology to known transporters[8], virulence factors[9][10], drug targets[11][12], and antibiotic resistance genes[13]. The number of genes and the specific source database where homology was found is provided (Table 4).

Table 4. Specialty Genes

| | Source | Genes |
|---|---|---|
| Antibiotic Resistance | PATRIC | 24 |
| Transporter | TCDB | 1 |

### Antimicrobial Resistance Genes

The Genome Annotation Service in PATRIC uses k-mer-based AMR genes detection method, which utilizes PATRIC's curated collection of representative AMR gene sequence variants[1] and assigns to each AMR gene functional annotation, broad mechanism of antibiotic resistance, drug class and, in some cases, specific antibiotic it confers resistance to. Please note, that the presence of AMR-related genes (even full length) in a given genome does not directly imply antibiotic resistant phenotype. It is important to consider specific AMR mechanisms and especially the absence/presence of SNP mutations conveying resistance. A summary of the AMR genes annotated in this genome and corresponding AMR mechanism is provided in Table 5.

Table 5. Antimicrobial Resistance Genes

| AMR Mechanism | Genes |
|---|---|
| Antibiotic inactivation enzyme | NimB |
| Antibiotic target in susceptible species | Alr, Ddl, dxr, EF-G, EF-Tu, folA, Dfr, gyrA, gyrB, Iso-tRNA, kasA, MurA, rho, rpoB, rpoC, S10p, S12p |
| Antibiotic target replacement protein | FabK |
| Gene conferring resistance via absence | gidB |
| Protein altering cell wall charge conferring antibiotic resistance | GdpD, PgsA |

### Phylogenetic Analysis

The National Center for Biotechnology Information (NCBI) staff manually select and categorize reference and representative genomes, which they consider to be of high quality and importance to the research community. PATRIC provides the reference and representative genomes, and includes them in the phylogenetic analysis that is part of the Comprehensive Genome Analysis report. The closest reference and representative genomes to were identified by Mash/MinHash[15]. PATRIC global protein families (PGFams)[6] were selected from these genomes to determine the phylogenetic placement of this genome. The protein sequences from these families were aligned with MUSCLE[17], and the nucleotides for each of those sequences were mapped to the protein alignment. The joint set of amino acid and nucleotide alignments were concatenated into a data matrix, and RaxML[18] was used to analyze this matrix, with fast bootstrapping[19] was used to generate the support values in the tree (Figure 3).

Figure 3

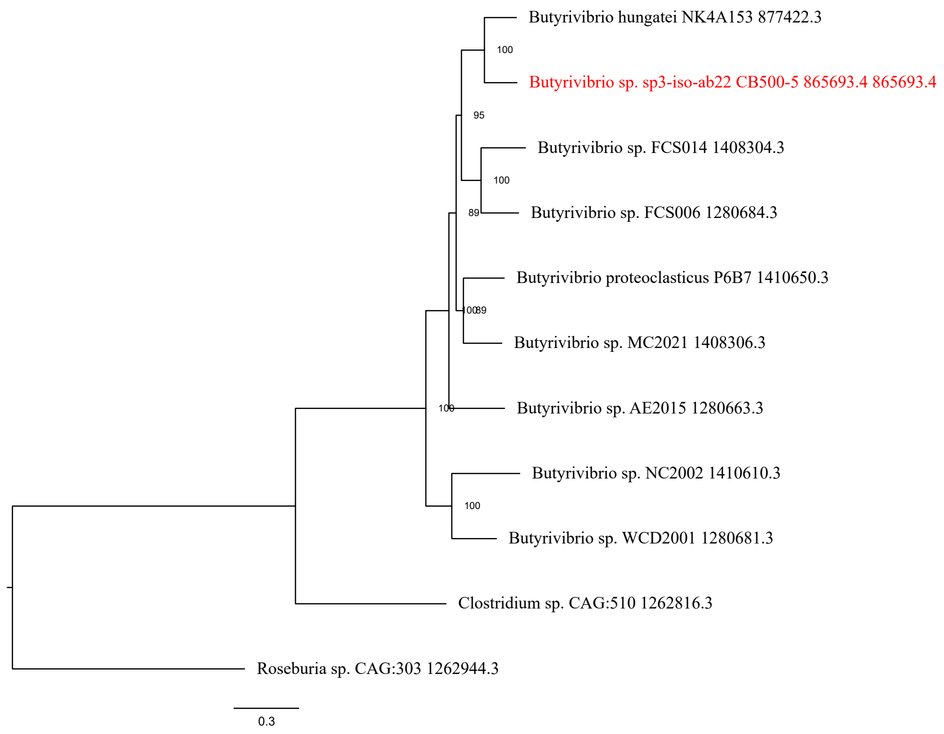


### References


1. Wattam AR, Davis JJ, Assaf R, Boisvert S, Brettin T, Bun C, Conrad N, Dietrich EM, Disz T, Gabbard JL, et al. 2017. Improvements to PATRIC, the all-bacterial Bioinformatics Database and Analysis Resource Center. Nucleic Acids Res 45:D535-D542.

2. Brettin T, Davis JJ, Disz T, Edwards RA, Gerdes S, Olsen GJ, Olson R, Overbeek R, Parrello B, Pusch GD, et al. 2015. RASTtk: a modular and extensible implementation of the RAST algorithm for building custom annotation pipelines and annotating batches of genomes. Sci Rep 5:8365.

3. Schomburg I, Chang A, Ebeling C, Gremse M, Heldt C, Huhn G, Schomburg D. 2004. BRENDA, the enzyme database: updates and major new developments. Nucleic Acids Res 32:D431-D433.

4. Ashburner M, Ball CA, Blake JA, Botstein D, Butler H, Cherry JM, Davis AP, Dolinski K, Dwight SS, Eppig JT. 2000. Gene Ontology: tool for the unification of biology. Nature genetics 25:25.

5. Kanehisa M, Sato Y, Kawashima M, Furumichi M, Tanabe M. 2016. KEGG as a reference resource for gene and protein annotation. Nucleic Acids Res 44:D457-462.

6. Davis JJ, Gerdes S, Olsen GJ, Olson R, Pusch GD, Shukla M, Vonstein V, Wattam AR, Yoo H. 2016. PATtyFams: Protein Families for the Microbial Genomes in the PATRIC Database. Front Microbiol 7:118.

7. Overbeek R, Begley T, Butler RM, Choudhuri JV, Chuang H-Y, Cohoon M, de Crécy-Lagard V, Diaz N, Disz T, Edwards R. 2005. The subsystems approach to genome annotation and its use in the project to annotate 1000 genomes. Nucleic Acids Res 33:5691-5702.

8. Saier Jr MH, Reddy VS, Tsu BV, Ahmed MS, Li C, Moreno-Hagelsieb G. 2015. The transporter classification database (TCDB): recent advances. Nucleic Acids Res 44:D372-D379.

9. Mao C, Abraham D, Wattam AR, Wilson MJ, Shukla M, Yoo HS, Sobral BW. 2015. Curation, integration and visualization of bacterial virulence factors in PATRIC. Bioinformatics 31:252-258.

10. Chen L, Zheng D, Liu B, Yang J, Jin Q. 2016. VFDB 2016: hierarchical and refined dataset for big data analysis-10 years on. Nucleic Acids Res 44:D694-D697.

11. Zhu F, Han B, Kumar P, Liu X, Ma X, Wei X, Huang L, Guo Y, Han L, Zheng C. 2009. Update of TTD: therapeutic target database. Nucleic Acids Res 38:D787-D791.

12. Law V, Knox C, Djoumbou Y, Jewison T, Guo AC, Liu Y, Maciejewski A, Arndt D, Wilson M, Neveu V, et al. 2014. DrugBank 4.0: shedding new light on drug metabolism. Nucleic Acids Res 42:D1091-1097.

13. McArthur AG, Waglechner N, Nizam F, Yan A, Azad MA, Baylay AJ, Bhullar K, Canova MJ, De Pascale G, Ejim L. 2013. The comprehensive antibiotic resistance database. Antimicrobial agents and chemotherapy 57:3348-3357.

14. Wattam AR, Davis JJ, Assaf R, Boisvert S, Brettin T, Bun C, Conrad N, Dietrich EM, Disz T, Gabbard JL, et al. 2017. Improvements to PATRIC, the all-bacterial Bioinformatics Database and Analysis Resource Center. Nucleic Acids Res 45:D535-D542.

15. Ondov BD, Treangen TJ, Melsted P, Mallonee AB, Bergman NH, Koren S, Phillippy AM. 2016. Mash: fast genome and metagenome distance estimation using MinHash. Genome biology 17:132.

16. Davis JJ, Gerdes S, Olsen GJ, Olson R, Pusch GD, Shukla M, Vonstein V, Wattam AR, Yoo H. 2016. PATtyFams: Protein Families for the Microbial Genomes in the PATRIC Database. Front Microbiol 7:118.

17. Edgar RC. 2004. MUSCLE: multiple sequence alignment with high accuracy and high throughput. Nucleic Acids Res 32:1792-1797.

18. Stamatakis A. 2014. RAxML version 8: a tool for phylogenetic analysis and post-analysis of large phylogenies. Bioinformatics 30:1312-1313.

19. Stamatakis A, Hoover P, Rougemont J. 2008. A rapid bootstrap algorithm for the RAxML web servers. Systematic biology 57:758-771.

Report Date: 03/31/2020
Organism: Butyrivibrio sp. X503

### Summary

An assembled genome for Butyrivibrio sp. X503 was submitted to the comprehensive genome analysis service at PATRIC[1]. Based on the annotation statistics and a comparison to other genomes in PATRIC within this same species, this genome appears to be of Good quality. Details of the analysis, including genes of interest (Specialty Genes), a functional categorization (Subsystems), and a phylogenetic tree (Phylogenetic Analysis) are provided below.

### Genome Assembly

An assembled genome was submitted to the Comprehensive Genome Analysis service. This assembled genome had 34 contigs, with the total length of 3,287,024 bp and an average G+C content of 42.17% (Table 1).

Table 1. Assembly Details

| | |
|---|---|
| Contigs | 34 |
| GC Content | 42.17 |
| Plasmids | 0 |
| Contig L50 | 3 |
| Genome Length | 3,287,024 bp |
| Contig N50 | 380,899 |
| Chromosomes | 0 |

### Genome Annotation

The Butyrivibrio sp. X503 genome was annotated using RAST tool kit (RASTtk)[2] and assigned a unique genome identifier of 28121.22. This genome is in the superkingdom and was annotated using genetic code 11. The taxonomy of this genome is:

> cellular organisms > Bacteria > Terrabacteria group > Firmicutes > Clostridia > Clostridiales > Lachnospiraceae > Butyrivibrio > unclassified Butyrivibrio > Butyrivibrio sp.

This genome has 3,080 protein coding sequences (CDS), 54 transfer RNA (tRNA) genes, and 6 ribosomal RNA (rRNA) genes. The annotated features are summarized in Table 2.

Table 2. Annotated Genome Features

| | |
|---|---|
| CDS | 3,080 |
| tRNA | 54 |
| rRNA | 6 |
| Partial CDS | 0 |
| Miscellaneous RNA | 0 |
| Repeat Regions | 0 |
| Job ID | annotation_13476 |
| Job Started | March 31st 2020, 7:03:03am |
| Job Completed | March 31st 2020, 7:13:28am |
| Total Time | 10 minutes and 25 seconds |

The annotation included 1,260 hypothetical proteins and 1,820 proteins with functional assignments (Table 3). The proteins with functional assignments included 639 proteins with Enzyme Commission (EC) numbers[3], 536 with Gene Ontology (GO) assignments[4], and 487 proteins that were mapped to KEGG pathways[5]. PATRIC annotation includes two types of protein families[6], and this genome has 2,271 proteins that belong to the genus-specific protein families (PLFams) for , and 2,340 proteins that belong to the cross-genus protein families (PGFams).

Table 3. Protein Features

| | |
|---|---|
| Hypothetical proteins | 1,260 |
| Proteins with functional assignments | 1,820 |
| Proteins with EC number assignments | 639 |
| Proteins with GO assignments | 536 |
| Proteins with Pathway assignments | 487 |
| Proteins with PATRIC genus-specific family (PLfam) assignments | 2,271 |
| Proteins with PATRIC cross-genus family (PGfam) assignments | 2,340 |

A circular graphical display of the distribution of the genome annotations is provided (Figure 1). This includes, from outer to inner rings, the contigs, CDS on the forward strand, CDS on the reverse strand, RNA genes, CDS with homology to known antimicrobial resistance genes, CDS with homology to know virulence factors, GC content and GC skew. The colors of the CDS on the forward and reverse strand indicate the subsystem that these genes belong to (see Subsystems below).

Figure 1

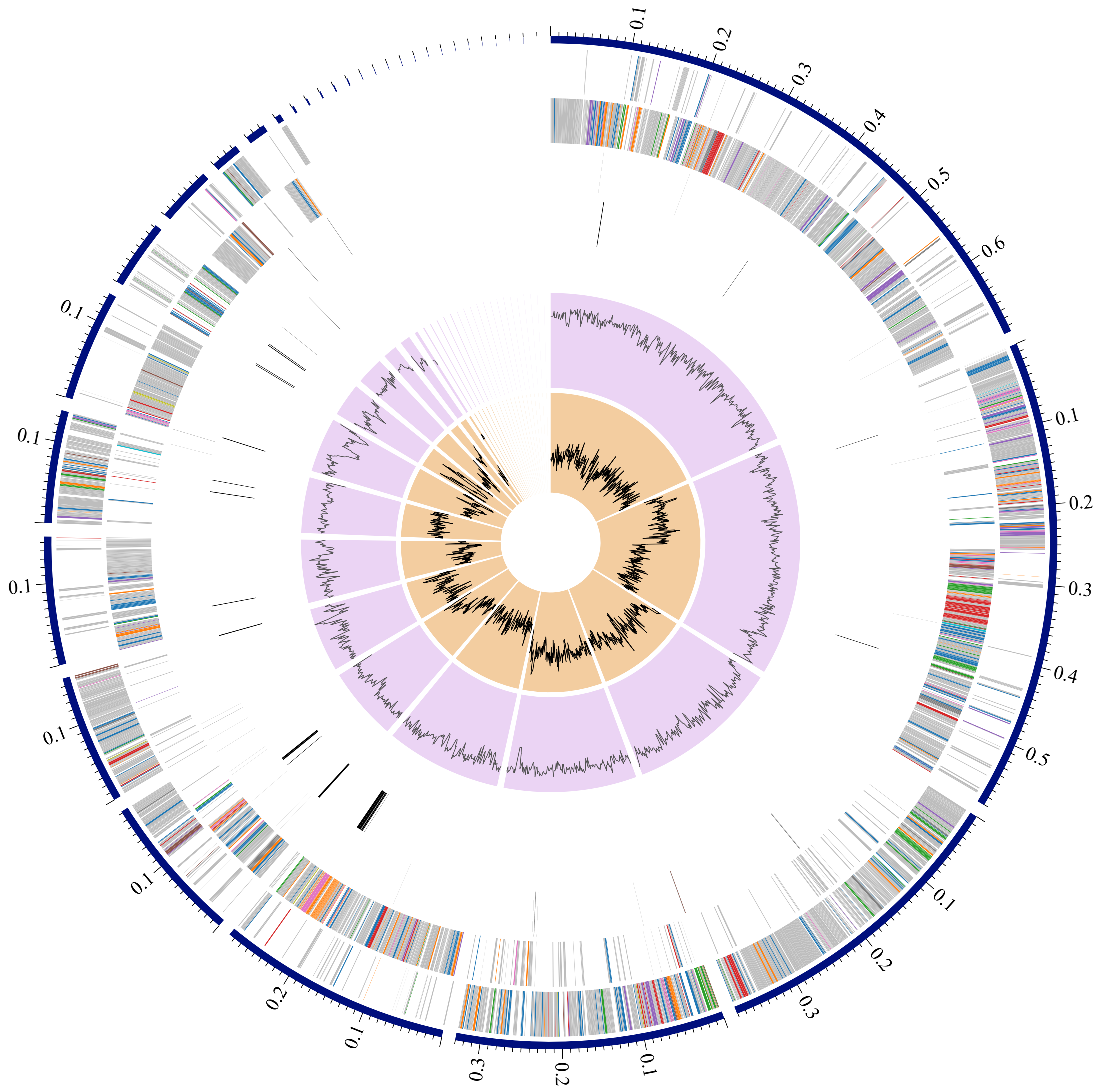

### Subsystem Analysis

A subsystem is a set of proteins that together implement a specific biological process or structural complex[7] and PATRIC annotation includes an analysis of the subsystems unique to each genome. An overview of the subsystems for this genome is provided in Figure 2.

Figure 2

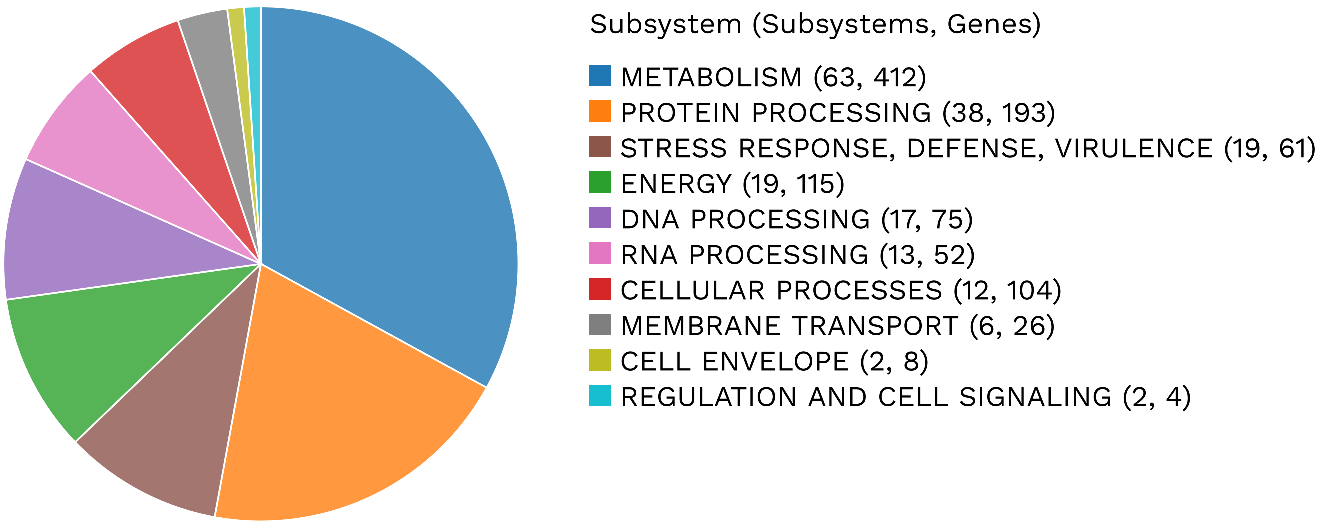

### Specialty Genes

Many of the genes annotated in have homology to known transporters[8], virulence factors[9][10], drug targets[11][12], and antibiotic resistance genes[13]. The number of genes and the specific source database where homology was found is provided (Table 4).

Table 4. Specialty Genes

| | Source | Genes |
|---|---|---|
| Antibiotic Resistance | PATRIC | 25 |
| Transporter | TCDB | 1 |

### Antimicrobial Resistance Genes

The Genome Annotation Service in PATRIC uses k-mer-based AMR genes detection method, which utilizes PATRIC's curated collection of representative AMR gene sequence variants[1] and assigns to each AMR gene functional annotation, broad mechanism of antibiotic resistance, drug class and, in some cases, specific antibiotic it confers resistance to. Please note, that the presence of AMR-related genes (even full length) in a given genome does not directly imply antibiotic resistant phenotype. It is important to consider specific AMR mechanisms and especially the absence/presence of SNP mutations conveying resistance. A summary of the AMR genes annotated in this genome and corresponding AMR mechanism is provided in Table 5.

Table 5. Antimicrobial Resistance Genes

| AMR Mechanism | Genes |
|---|---|
| Antibiotic inactivation enzyme | NimB |
| Antibiotic target in susceptible species | Alr, Ddl, dxr, EF-G, EF-Tu, folA, Dfr, gyrA, gyrB, Iso-tRNA, kasA, MurA, rho, rpoB, rpoC, S10p, S12p |
| Antibiotic target replacement protein | FabK |
| Gene conferring resistance via absence | gidB |
| Protein altering cell wall charge conferring antibiotic resistance | GdpD, PgsA |

### Phylogenetic Analysis

The National Center for Biotechnology Information (NCBI) staff manually select and categorize reference and representative genomes, which they consider to be of high quality and importance to the research community. PATRIC provides the reference and representative genomes, and includes them in the phylogenetic analysis that is part of the Comprehensive Genome Analysis report. The closest reference and representative genomes to were identified by Mash/MinHash[15]. PATRIC global protein families (PGFams)[6] were selected from these genomes to determine the phylogenetic placement of this genome. The protein sequences from these families were aligned with MUSCLE[17], and the nucleotides for each of those sequences were mapped to the protein alignment. The joint set of amino acid and nucleotide alignments were concatenated into a data matrix, and RaxML[18] was used to analyze this matrix, with fast bootstrapping[19] was used to generate the support values in the tree (Figure 3).

Figure 3

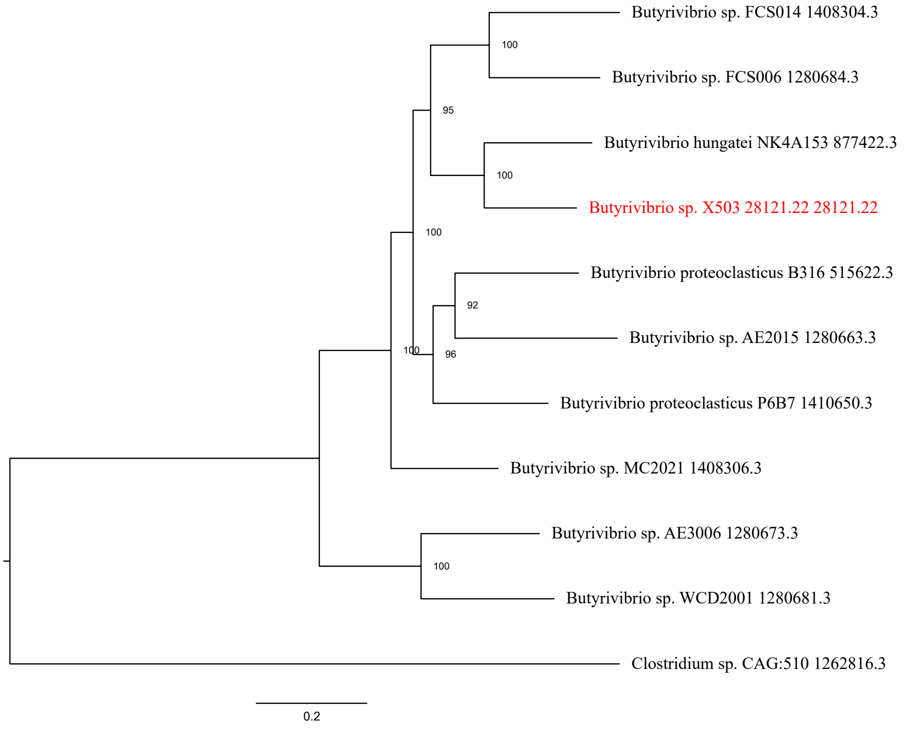


### References


1. Wattam AR, Davis JJ, Assaf R, Boisvert S, Brettin T, Bun C, Conrad N, Dietrich EM, Disz T, Gabbard JL, et al. 2017. Improvements to PATRIC, the all-bacterial Bioinformatics Database and Analysis Resource Center. Nucleic Acids Res 45:D535-D542.

2. Brettin T, Davis JJ, Disz T, Edwards RA, Gerdes S, Olsen GJ, Olson R, Overbeek R, Parrello B, Pusch GD, et al. 2015. RASTtk: a modular and extensible implementation of the RAST algorithm for building custom annotation pipelines and annotating batches of genomes. Sci Rep 5:8365.

3. Schomburg I, Chang A, Ebeling C, Gremse M, Heldt C, Huhn G, Schomburg D. 2004. BRENDA, the enzyme database: updates and major new developments. Nucleic Acids Res 32:D431-D433.

4. Ashburner M, Ball CA, Blake JA, Botstein D, Butler H, Cherry JM, Davis AP, Dolinski K, Dwight SS, Eppig JT. 2000. Gene Ontology: tool for the unification of biology. Nature genetics 25:25.

5. Kanehisa M, Sato Y, Kawashima M, Furumichi M, Tanabe M. 2016. KEGG as a reference resource for gene and protein annotation. Nucleic Acids Res 44:D457-462.

6. Davis JJ, Gerdes S, Olsen GJ, Olson R, Pusch GD, Shukla M, Vonstein V, Wattam AR, Yoo H. 2016. PATtyFams: Protein Families for the Microbial Genomes in the PATRIC Database. Front Microbiol 7:118.

7. Overbeek R, Begley T, Butler RM, Choudhuri JV, Chuang H-Y, Cohoon M, de Crécy-Lagard V, Diaz N, Disz T, Edwards R. 2005. The subsystems approach to genome annotation and its use in the project to annotate 1000 genomes. Nucleic Acids Res 33:5691-5702.

8. Saier Jr MH, Reddy VS, Tsu BV, Ahmed MS, Li C, Moreno-Hagelsieb G. 2015. The transporter classification database (TCDB): recent advances. Nucleic Acids Res 44:D372-D379.

9. Mao C, Abraham D, Wattam AR, Wilson MJ, Shukla M, Yoo HS, Sobral BW. 2015. Curation, integration and visualization of bacterial virulence factors in PATRIC. Bioinformatics 31:252-258.

10. Chen L, Zheng D, Liu B, Yang J, Jin Q. 2016. VFDB 2016: hierarchical and refined dataset for big data analysis-10 years on. Nucleic Acids Res 44:D694-D697.

11. Zhu F, Han B, Kumar P, Liu X, Ma X, Wei X, Huang L, Guo Y, Han L, Zheng C. 2009. Update of TTD: therapeutic target database. Nucleic Acids Res 38:D787-D791.
12. Law V, Knox C, Djoumbou Y, Jewison T, Guo AC, Liu Y, Maciejewski A, Arndt D, Wilson M, Neveu V, et al. 2014. DrugBank 4.0: shedding new light on drug metabolism. Nucleic Acids Res 42:D1091-1097.
13. McArthur AG, Waglechner N, Nizam F, Yan A, Azad MA, Baylay AJ, Bhullar K, Canova MJ, De Pascale G, Ejim L. 2013. The comprehensive antibiotic resistance database. Antimicrobial agents and chemotherapy 57:3348-3357.
14. Wattam AR, Davis JJ, Assaf R, Boisvert S, Brettin T, Bun C, Conrad N, Dietrich EM, Disz T, Gabbard JL, et al. 2017. Improvements to PATRIC, the all-bacterial Bioinformatics Database and Analysis Resource Center. Nucleic Acids Res 45:D535-D542.
15. Ondov BD, Treangen TJ, Melsted P, Mallonee AB, Bergman NH, Koren S, Phillippy AM. 2016. Mash: fast genome and metagenome distance estimation using MinHash. Genome biology 17:132.
16. Davis JJ, Gerdes S, Olsen GJ, Olson R, Pusch GD, Shukla M, Vonstein V, Wattam AR, Yoo H. 2016. PATtyFams: Protein Families for the Microbial Genomes in the PATRIC Database. Front Microbiol 7:118.
17. Edgar RC. 2004. MUSCLE: multiple sequence alignment with high accuracy and high throughput. Nucleic Acids Res 32:1792-1797.
18. Stamatakis A. 2014. RAxML version 8: a tool for phylogenetic analysis and post-analysis of large phylogenies. Bioinformatics 30:1312-1313.
19. Stamatakis A, Hoover P, Rougemont J. 2008. A rapid bootstrap algorithm for the RAxML web servers. Systematic biology 57:758-771.
