## Supplementary figures and images for "Genomic architecture of three newly isolated unclassified *Butyrivibrio* species elucidate their potential role in the rumen ecosystem"

### Supplementary file 2a

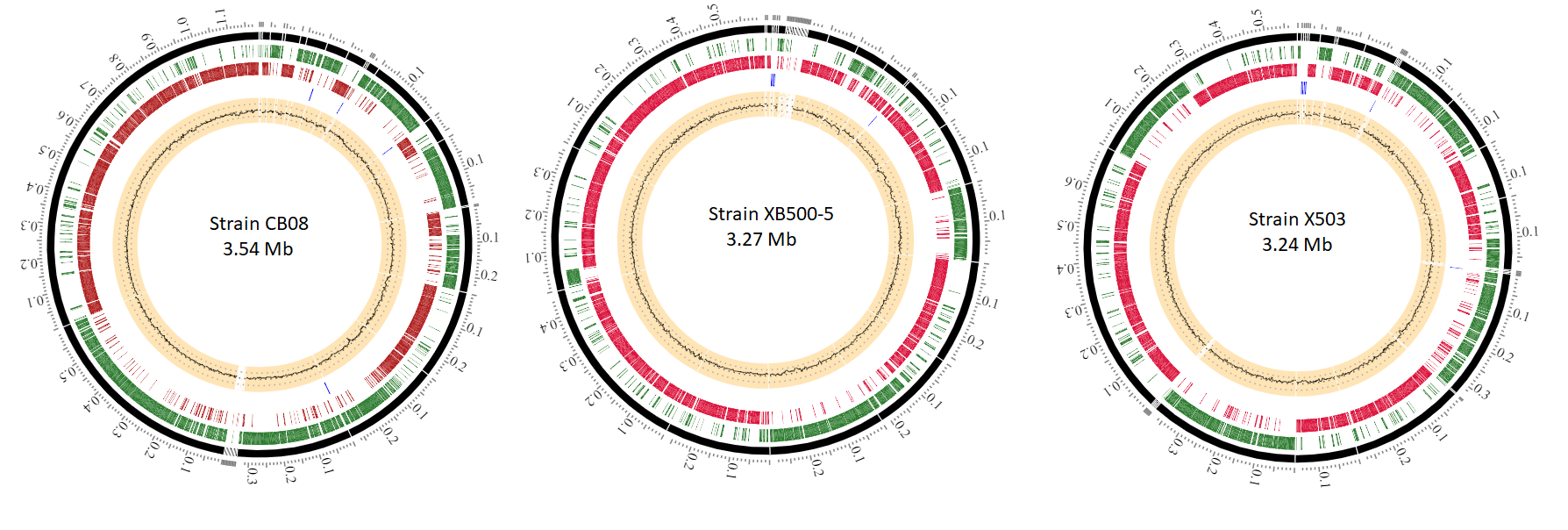

### Supplementary file 2b

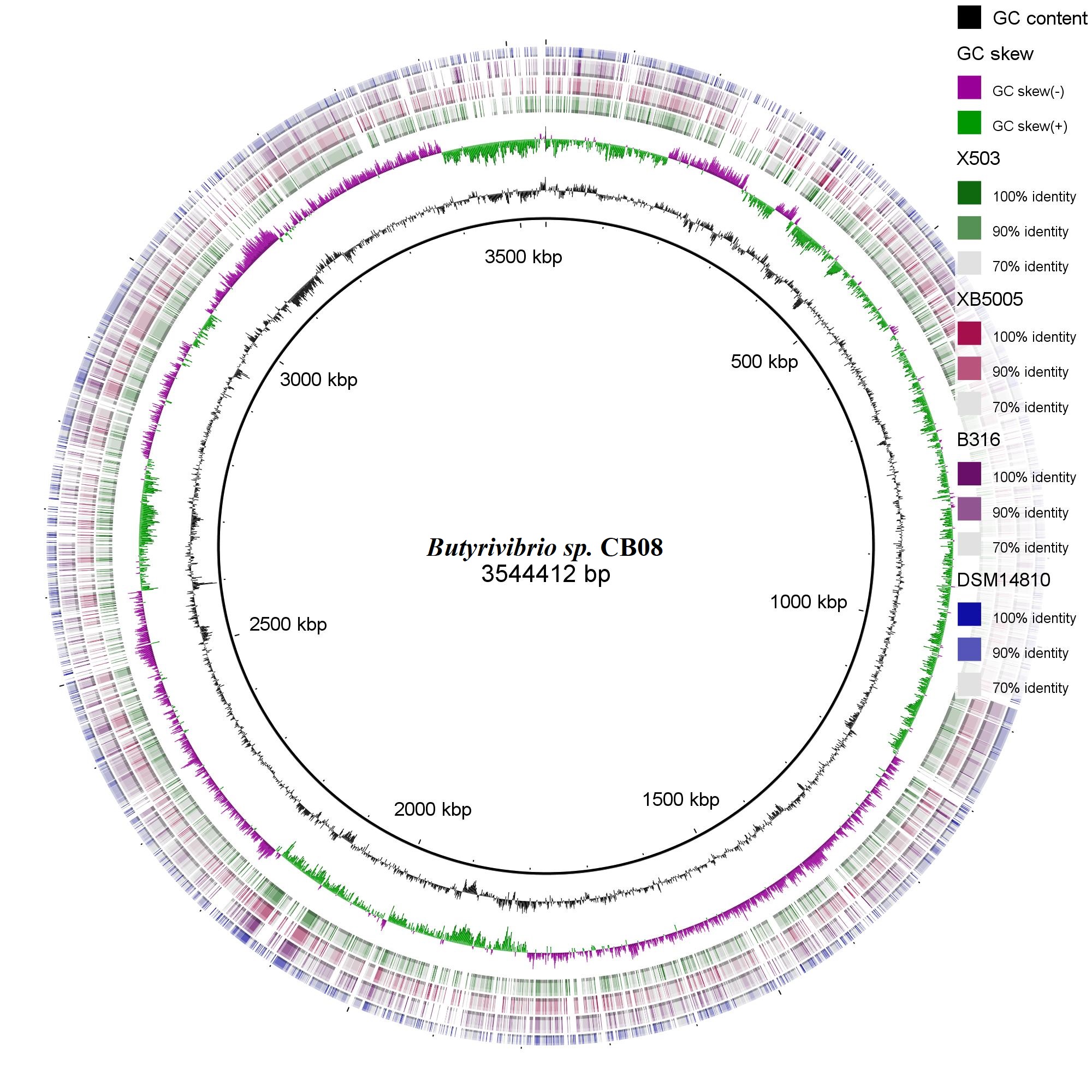

### Supplementary file 6

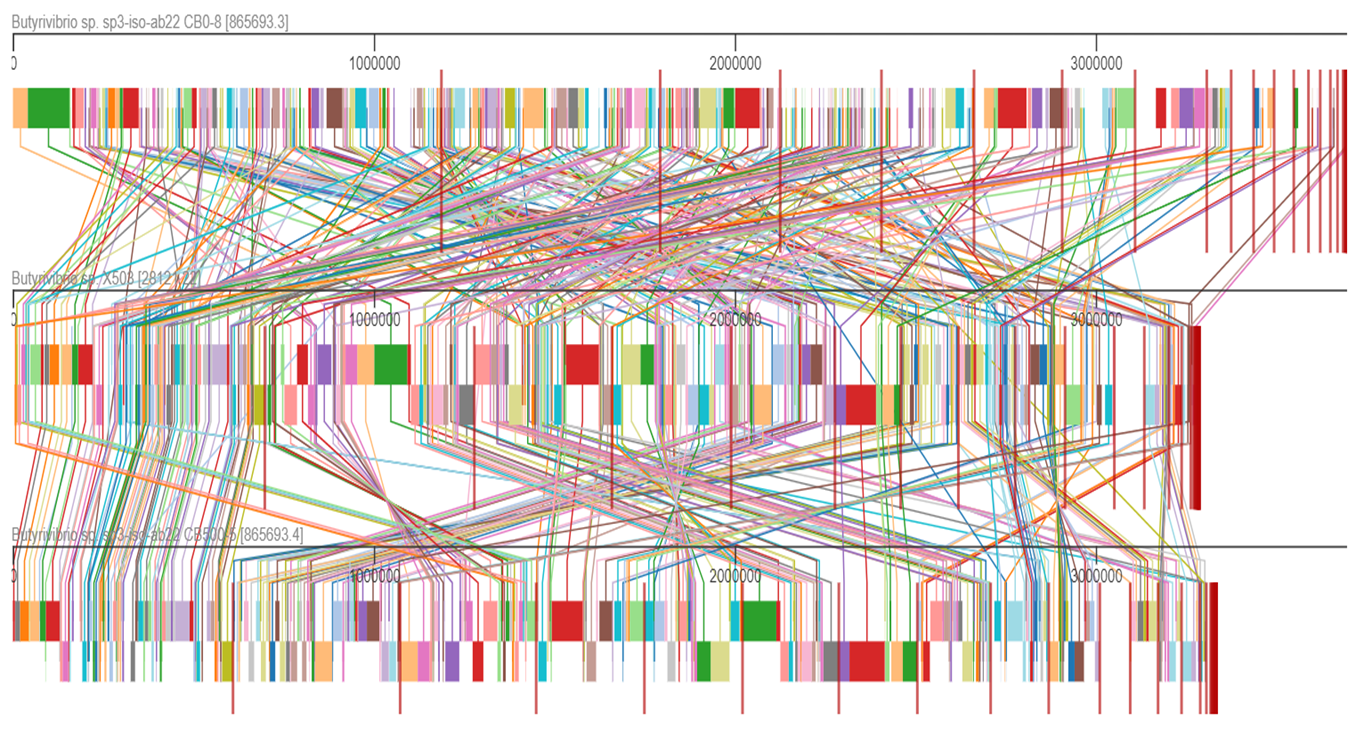

### Supplementary file 7

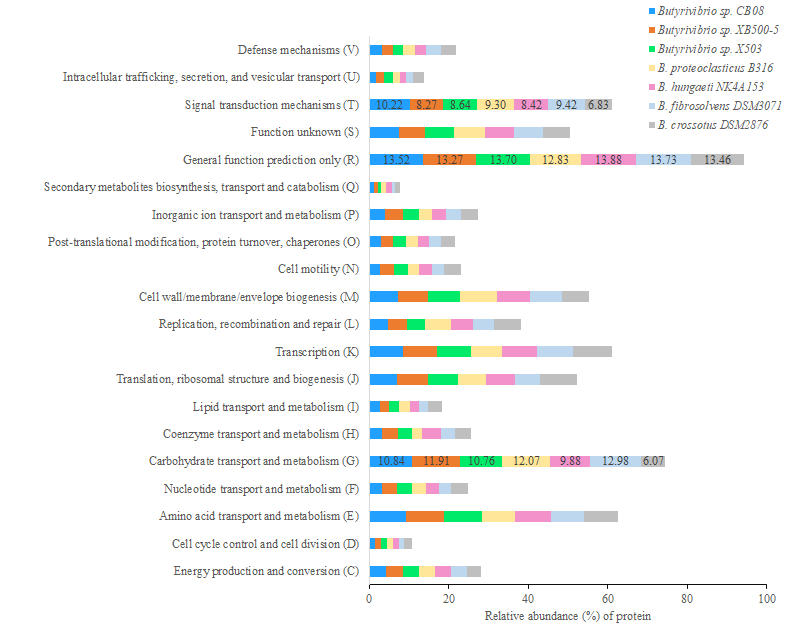
