## Supplementary file 4 for "Genomic architecture of three newly isolated unclassified *Butyrivibrio* species elucidate their potential role in the rumen ecosystem"

**Table S4a.** ANI value for pair wise genome comparisons between *Butyrivibrio* spp. (strains CB08, XB500-5 and X503) and type strains *Butyrivibrio* *hungatei* DSM 14810^T^, *Butyrivibrio* *proteoclasticus* B316^T^. The ANI values were inferred using three different ANI calculators. All values are in %; SD = standard deviation.

| **Genome 1** | **Genome 2** | **ANI calculators by** | | | **Average ANI** | **SD** |
| --- | --- | --- | --- | --- | --- | --- |
|  |  | **JGI-IMG** | **Kostas** | **ChunLab** |  |  |
| *Butyrivibrio* sp. CB08 | *Butyrivibrio* *proteoclasticus* B316^T^ | 75.93 | 80 | 74.71 | **76.88** | 2.77 |
| *Butyrivibrio* sp. XB500-5 | *Butyrivibrio* *hungatei* DSM 14810^T^ | 75.93 | 80 | 74.79 | **76.91** | 2.74 |
| *Butyrivibrio* sp. X503 | *Butyrivibrio* *hungatei* DSM 14810^T^ | 75.84 | 80 | 74.64 | **76.83** | 2.81 |
| *Butyrivibrio* sp. XB500-5 | *Butyrivibrio* sp. X503 | 90.52 | 91 | 90.34 | **90.62** | 0.34 |
| *Butyrivibrio* sp. CB08 | *Butyrivibrio* sp. XB500-5 | 75.28 | 80 | 74.04 | **76.44** | 3.14 |
| *Butyrivibrio* sp. CB08 | *Butyrivibrio* sp. X503 | 75.49 | 80 | 74.01 | **76.50** | 3.12 |

**Table S4b.** Comparative account of dDDH, 16S rDNA sequence similarity and GC difference between *Butyrivibrio* spp. (strains CB08, XB500-5 and X503) and type strains *Butyrivibrio* *hungatei* DSM 14810^T^, *Butyrivibrio* *proteoclasticus* B316^T^. All values are in %. Value of dDDH >60%, 16S rDNA sequence similarity >98% and GC difference <1% are shaded in grey.

|  | ***Butyrivibrio* sp. CB08** | | | ***Butyrivibrio* sp. XB500-5** | | | ***Butyrivibrio* sp. X503** | | |
| --- | --- | --- | --- | --- | --- | --- | --- | --- | --- |
|  | dDDH | 16S rDNA | GC diff | dDDH | 16S rDNA | GC diff | dDDH | 16S rDNA | GC diff |
| ***Butyrivibrio* sp. CB08** | 100 | 100 | 100 | 16 | 97.3 | 1.3 | 16 | 97.6 | 1.5 |
| ***Butyrivibrio* sp. XB500-5** | 16 | 97.3 | 1.3 | 100 | 100.0 | 100.0 | 66 | 98.9 | 0.2 |
| ***Butyrivibrio* sp. X503** | 16 | 97.6 | 1.5 | 66 | 98.9 | 0.2 | 100 | 100.0 | 100.0 |
| ***B. proteoclasticus* B316^T^** | 15.4 | 96.3 | 3.7 | 14.8 | 96.9 | 2.4 | 14.8 | 97.0 | 2.2 |
| ***B. hungatei* DSM 14810^T^** | 17.1 | 96.4 | 3.8 | 16.8 | 97.7 | 2.5 | 16.8 | 97.7 | 2.3 |
