## Supplementary file 5 for "Genomic architecture of three newly isolated unclassified *Butyrivibrio* species elucidate their potential role in the rumen ecosystem"

TYPE (STRAIN) GENOME SERVER

### JOB SUMMARY

| | |
|---|---|
| PRINT DATE: | 2020-04-09 07:20:49 +0200 |
| JOB ID: | ce213f76-fce9-4c50-a681-d9763c10a3d6 |
| RESULT PAGE: | https://tygs.dsmz.de/user_results/show?guid=ce213f76-fce9-4c50-a681-d9763c10a3d6 |

#### Table 1: Phylogenies

**Publication-ready versions** of both the genome-scale GBDP tree and the 16S rRNA gene sequence tree can be customized and exported either in SVG (vector graphic) or PNG format from within the phylogeny viewers in your TYGS result page. For publications the **SVG format is recommended** because it is lossless, always keeps its high resolution and can also be easily converted to other popular formats such as PDF or EPS. Please follow the link provided above!

#### Table 2: Identification

The below list contains the result of the TYGS species identification routine.

Explanation of remarks that might occur in the below table:

**remark [R1]:** The TYGS type strain database is automatically updated on an almost daily basis. However, if a particular type strain genome is not available in the TYGS database, this can have several reasons which are detailed in the FAQ. You can request an extended 16S rRNA gene analysis via the 16S tree viewer found in your result page to detect **not yet genome-sequenced** type strains relevant for your study.

**remark [R2]:** > 70% dDDH value (formula $d_4$) and (almost) minimal dDDH values for gene-content formulae $d_0$ and $d_6$ indicate a potentially unreliable identification result and should thus be checked via the 16S rRNA gene sequence similarity. Such strong deviations can, in principle, be caused by sequence contamination.

**remark [R3]:** G+C content difference of > 1 % indicates a potentially unreliable identification result because within species G+C content varies no more than 1 %, if computed from genome sequences (PMID: 24505073).

| Strain | Conclusion | Identification result | Remark |
|---|---|---|---|
| **'Butyrivibrio sp. X503' (RAIQ01000001-RAIQ01000019)** | potential new species | | see [R1] |
| **'Butyrivibrio sp. CB08' (RAIR01000001-RAIR01000029)** | potential new species | | see [R1] |
| **'Butyrivibrio sp. XB500-5' (RAIS01000001-RAIS01000016)** | potential new species | | see [R1] |

#### Table 3: Pairwise comparisons of user genomes vs. type-strain genomes

The following table contains the pairwise dDDH values between your user genomes and the selected type-strain genomes. The dDDH values are provided along with their confidence intervals (C.I.) for the three different GBDP formulas:

- formula $d_0$ (a.k.a. GGDC formula 1): length of all HSPs divided by total genome length
- formula $d_4$ (a.k.a. GGDC formula 2): sum of all identities found in HSPs divided by overall HSP length
- formula $d_6$ (a.k.a. GGDC formula 3): sum of all identities found in HSPs divided by total genome length

**Note:** Formula $d_4$ is independent of genome length and is thus robust against the use of incomplete draft genomes. For other reasons for preferring formula $d_4$, see the FAQ.

| Query | Subject | $d_0$ | C.I. $d_0$ | $d_4$ | C.I. $d_4$ | $d_6$ | C.I. $d_6$ | Diff. G+C Percent |
|---|---|---|---|---|---|---|---|---|
| **'Butyrivibrio sp. X503' (RAIQ01000001-RAIQ01000019)** | **'Butyrivibrio sp. XB500-5' (RAIS01000001-RAIS01000016)** | 73.1 | [69.1 - 76.7] | 39.9 | [37.4 - 42.4] | 66.0 | [62.6 - 69.2] | 0.21 |
| **'Butyrivibrio sp. CB08' (RAIR01000001-RAIR01000029)** | *Enterocloster clostridioformis* ATCC 25537 | 12.5 | [9.8 - 15.8] | 30.7 | [28.3 - 33.2] | 12.9 | [10.6 - 15.7] | 5.26 |
| **'Butyrivibrio sp. CB08' (RAIR01000001-RAIR01000029)** | *Enterocloster bolteae* ATCC BAA-613 | 12.5 | [9.9 - 15.8] | 30.2 | [27.8 - 32.7] | 12.9 | [10.6 - 15.7] | 5.36 |
| **'Butyrivibrio sp. CB08' (RAIR01000001-RAIR01000029)** | *Anaerocolumna jejuensis* DSM 15929 | 12.5 | [9.8 - 15.8] | 28.8 | [26.4 - 31.3] | 12.9 | [10.6 - 15.7] | 3.08 |
| **'Butyrivibrio sp. CB08' (RAIR01000001-RAIR01000029)** | *Eubacterium cellulosolvens* 6 | 12.6 | [9.9 - 15.9] | 27.1 | [24.8 - 29.6] | 13.0 | [10.7 - 15.8] | 4.48 |
| **'Butyrivibrio sp. X503' (RAIQ01000001-RAIQ01000019)** | *Anaerocolumna jejuensis* DSM 15929 | 12.5 | [9.9 - 15.8] | 26.7 | [24.3 - 29.2] | 12.9 | [10.6 - 15.7] | 1.6 |
| **'Butyrivibrio sp. CB08' (RAIR01000001-RAIR01000029)** | *Blautia hansenii* DSM 20583 | 12.6 | [9.9 - 15.8] | 26.3 | [24.0 - 28.8] | 13.0 | [10.6 - 15.7] | 4.71 |
| **'Butyrivibrio sp. X503' (RAIQ01000001-RAIQ01000019)** | *Anaerostipes rhamnosivorans* 1y-2 | 12.6 | [9.9 - 15.8] | 26.2 | [23.9 - 28.7] | 13.0 | [10.6 - 15.7] | 2.32 |
| **'Butyrivibrio sp. XB500-5' (RAIS01000001-RAIS01000016)** | *Anaerocolumna jejuensis* DSM 15929 | 12.5 | [9.9 - 15.8] | 25.9 | [23.5 - 28.4] | 12.9 | [10.6 - 15.7] | 1.81 |
| **'Butyrivibrio sp. X503' (RAIQ01000001-RAIQ01000019)** | *Lacrimispora celerecrescens* 18A | 12.6 | [9.9 - 15.9] | 25.4 | [23.1 - 27.9] | 13.0 | [10.7 - 15.7] | 1.68 |
| **'Butyrivibrio sp. CB08' (RAIR01000001-RAIR01000029)** | *Kineothrix alysoides* KNHs209 | 12.6 | [9.9 - 15.8] | 25.1 | [22.8 - 27.6] | 13.0 | [10.7 - 15.7] | 0.95 |
| **'Butyrivibrio sp. CB08' (RAIR01000001-RAIR01000029)** | *Anaerostipes rhamnosivorans* 1y-2 | 12.6 | [9.9 - 15.8] | 25.1 | [22.8 - 27.6] | 13.0 | [10.6 - 15.7] | 0.83 |
| **'Butyrivibrio sp. X503' (RAIQ01000001-RAIQ01000019)** | *Lachnobacterium bovis* DSM 14045 | 12.5 | [9.9 - 15.8] | 24.5 | [22.2 - 27.0] | 12.9 | [10.6 - 15.7] | 10.86 |
| **'Butyrivibrio sp. XB500-5' (RAIS01000001-RAIS01000016)** | *Anaerostipes rhamnosivorans* 1y-2 | 12.6 | [9.9 - 15.8] | 24.0 | [21.7 - 26.5] | 13.0 | [10.6 - 15.7] | 2.1 |
| **'Butyrivibrio sp. XB500-5' (RAIS01000001-RAIS01000016)** | *Lacrimispora celerecrescens* 18A | 12.6 | [9.9 - 15.9] | 24.0 | [21.7 - 26.5] | 13.0 | [10.7 - 15.7] | 1.46 |
| **'Butyrivibrio sp. X503' (RAIQ01000001-RAIQ01000019)** | *Blautia hansenii* DSM 20583 | 12.6 | [9.9 - 15.8] | 24.0 | [21.7 - 26.5] | 13.0 | [10.6 - 15.7] | 3.22 |

| Query | Subject | $d_0$ | C.I. $d_0$ | $d_4$ | C.I. $d_4$ | $d_6$ | C.I. $d_6$ | Diff. G+C Percent |
|---|---|---|---|---|---|---|---|---|
| **'Butyrivibrio sp. XB500-5' (RAIS01000001-RAIS01000016)** | *Coprococcus eutactus* ATCC 27759 | 12.7 | [10.0 - 16.0] | 23.9 | [21.6 - 26.3] | 13.1 | [10.8 - 15.9] | 0.66 |
| **'Butyrivibrio sp. XB500-5' (RAIS01000001-RAIS01000016)** | *Blautia hansenii* DSM 20583 | 12.6 | [9.9 - 15.8] | 23.6 | [21.3 - 26.0] | 13.0 | [10.6 - 15.7] | 3.44 |
| **'Butyrivibrio sp. X503' (RAIQ01000001-RAIQ01000019)** | *Eubacterium cellulosolvens* 6 | 12.7 | [10.0 - 15.9] | 23.6 | [21.3 - 26.1] | 13.1 | [10.7 - 15.8] | 5.97 |
| **'Butyrivibrio sp. XB500-5' (RAIS01000001-RAIS01000016)** | *Eubacterium cellulosolvens* 6 | 12.7 | [10.0 - 16.0] | 23.5 | [21.2 - 26.0] | 13.1 | [10.8 - 15.8] | 5.75 |
| **'Butyrivibrio sp. CB08' (RAIR01000001-RAIR01000029)** | *Lacrimispora celerecrescens* 18A | 12.6 | [9.9 - 15.9] | 23.4 | [21.1 - 25.9] | 13.0 | [10.7 - 15.8] | 0.19 |
| **'Butyrivibrio sp. X503' (RAIQ01000001-RAIQ01000019)** | *Kineothrix alysoides* KNHs209 | 12.6 | [9.9 - 15.9] | 22.9 | [20.6 - 25.3] | 13.0 | [10.7 - 15.7] | 0.53 |
| **'Butyrivibrio sp. CB08' (RAIR01000001-RAIR01000029)** | *Kineothrix alysoides* DSM 100556 | 12.5 | [9.9 - 15.8] | 22.7 | [20.4 - 25.1] | 12.9 | [10.6 - 15.7] | 1.0 |
| **'Butyrivibrio sp. CB08' (RAIR01000001-RAIR01000029)** | *Eubacterium ruminantium* ATCC 17233 | 12.7 | [10.0 - 16.0] | 22.6 | [20.3 - 25.0] | 13.1 | [10.7 - 15.8] | 6.46 |
| **'Butyrivibrio sp. XB500-5' (RAIS01000001-RAIS01000016)** | *Dorea phocaeensis* Marseille-P4003 | 12.6 | [9.9 - 15.8] | 22.6 | [20.3 - 25.0] | 13.0 | [10.6 - 15.7] | 0.74 |
| **'Butyrivibrio sp. X503' (RAIQ01000001-RAIQ01000019)** | *Dorea phocaeensis* Marseille-P4003 | 12.6 | [9.9 - 15.8] | 22.6 | [20.3 - 25.1] | 13.0 | [10.6 - 15.7] | 0.96 |
| **'Butyrivibrio sp. CB08' (RAIR01000001-RAIR01000029)** | *Merdimonas faecis* BR31 | 12.5 | [9.9 - 15.8] | 22.5 | [20.2 - 25.0] | 12.9 | [10.6 - 15.7] | 3.26 |
| **'Butyrivibrio sp. X503' (RAIQ01000001-RAIQ01000019)** | *Coprococcus eutactus* ATCC 27759 | 12.8 | [10.1 - 16.0] | 22.5 | [20.2 - 24.9] | 13.1 | [10.8 - 15.9] | 0.88 |
| **'Butyrivibrio sp. CB08' (RAIR01000001-RAIR01000029)** | *Eubacterium ventriosum* ATCC 27560 | 12.6 | [9.9 - 15.9] | 22.5 | [20.2 - 25.0] | 13.0 | [10.7 - 15.7] | 8.78 |
| **'Butyrivibrio sp. X503' (RAIQ01000001-RAIQ01000019)** | *Roseburia intestinalis* L1-82 | 12.6 | [9.9 - 15.8] | 22.4 | [20.1 - 24.8] | 13.0 | [10.6 - 15.7] | 0.42 |
| **'Butyrivibrio sp. CB08' (RAIR01000001-RAIR01000029)** | *Lachnobacterium bovis* DSM 14045 | 12.5 | [9.9 - 15.8] | 22.1 | [19.8 - 24.5] | 12.9 | [10.6 - 15.7] | 12.35 |
| **'Butyrivibrio sp. CB08' (RAIR01000001-RAIR01000029)** | *Butyrivibrio proteoclasticus* B316 | 15.2 | [12.3 - 18.7] | 21.7 | [19.5 - 24.2] | 15.4 | [12.9 - 18.2] | 3.69 |
| **'Butyrivibrio sp. X503' (RAIQ01000001-RAIQ01000019)** | *Roseburia faecis* M72 | 12.6 | [9.9 - 15.9] | 21.6 | [19.3 - 24.0] | 13.0 | [10.7 - 15.7] | 0.75 |
| **'Butyrivibrio sp. XB500-5' (RAIS01000001-RAIS01000016)** | *Kineothrix alysoides* KNHs209 | 12.6 | [9.9 - 15.9] | 21.6 | [19.4 - 24.0] | 13.0 | [10.7 - 15.8] | 0.32 |
| **'Butyrivibrio sp. X503' (RAIQ01000001-RAIQ01000019)** | *Butyrivibrio proteoclasticus* B316 | 14.6 | [11.7 - 18.0] | 21.5 | [19.2 - 23.9] | 14.8 | [12.3 - 17.6] | 2.2 |
| **'Butyrivibrio sp. XB500-5' (RAIS01000001-RAIS01000016)** | *Butyrivibrio proteoclasticus* B316 | 14.6 | [11.8 - 18.0] | 21.4 | [19.2 - 23.9] | 14.8 | [12.4 - 17.7] | 2.42 |
| **'Butyrivibrio sp. CB08' (RAIR01000001-RAIR01000029)** | *Butyrivibrio fibrisolvens* DSM 3071 | 13.2 | [10.5 - 16.5] | 21.4 | [19.1 - 23.8] | 13.6 | [11.2 - 16.4] | 4.04 |

| Query | Subject | $d_0$ | C.I. $d_0$ | $d_4$ | C.I. $d_4$ | $d_6$ | C.I. $d_6$ | Diff. G+C Percent |
|---|---|---|---|---|---|---|---|---|
| **'Butyrivibrio sp. XB500-5' (RAIS01000001-RAIS01000016)** | *Eubacterium oxidoreducens* DSM 3217 | 12.5 | [9.9 - 15.8] | 21.4 | [19.1 - 23.8] | 13.0 | [10.6 - 15.7] | 2.57 |
| **'Butyrivibrio sp. X503' (RAIQ01000001-RAIQ01000019)** | *Enterocloster bolteae* ATCC BAA-613 | 12.5 | [9.9 - 15.8] | 21.4 | [19.2 - 23.9] | 12.9 | [10.6 - 15.7] | 6.85 |
| **'Butyrivibrio sp. XB500-5' (RAIS01000001-RAIS01000016)** | *Mesorhizobium loti* DSM 2626 | 12.5 | [9.8 - 15.8] | 21.3 | [19.1 - 23.7] | 12.9 | [10.6 - 15.6] | 19.93 |
| **'Butyrivibrio sp. X503' (RAIQ01000001-RAIQ01000019)** | *Eubacterium oxidoreducens* DSM 3217 | 12.6 | [9.9 - 15.8] | 21.2 | [19.0 - 23.6] | 13.0 | [10.7 - 15.7] | 2.35 |
| **'Butyrivibrio sp. CB08' (RAIR01000001-RAIR01000029)** | *Roseburia intestinalis* L1-82 | 12.6 | [9.9 - 15.8] | 21.2 | [18.9 - 23.6] | 13.0 | [10.7 - 15.7] | 1.06 |
| **'Butyrivibrio sp. XB500-5' (RAIS01000001-RAIS01000016)** | *Butyrivibrio fibrisolvens* DSM 3071 | 13.1 | [10.4 - 16.4] | 21.2 | [19.0 - 23.7] | 13.4 | [11.1 - 16.2] | 2.77 |
| **'Butyrivibrio sp. X503' (RAIQ01000001-RAIQ01000019)** | *Enterocloster clostridioformis* ATCC 25537 | 12.5 | [9.9 - 15.8] | 21.1 | [18.8 - 23.5] | 12.9 | [10.6 - 15.7] | 6.74 |
| **'Butyrivibrio sp. CB08' (RAIR01000001-RAIR01000029)** | *Coprococcus eutactus* ATCC 27759 | 12.8 | [10.1 - 16.0] | 21.1 | [18.9 - 23.5] | 13.1 | [10.8 - 15.9] | 0.61 |
| **'Butyrivibrio sp. CB08' (RAIR01000001-RAIR01000029)** | *Dorea phocaeensis* Marseille-P4003 | 12.6 | [9.9 - 15.9] | 20.9 | [18.6 - 23.3] | 13.0 | [10.7 - 15.7] | 0.53 |
| **'Butyrivibrio sp. XB500-5' (RAIS01000001-RAIS01000016)** | *Roseburia intestinalis* L1-82 | 12.6 | [9.9 - 15.8] | 20.8 | [18.6 - 23.3] | 13.0 | [10.7 - 15.7] | 0.21 |
| **'Butyrivibrio sp. X503' (RAIQ01000001-RAIQ01000019)** | *Butyrivibrio fibrisolvens* DSM 3071 | 13.1 | [10.3 - 16.3] | 20.8 | [18.6 - 23.3] | 13.4 | [11.0 - 16.2] | 2.55 |
| **'Butyrivibrio sp. XB500-5' (RAIS01000001-RAIS01000016)** | *Clostridium aminophilum* DSM 10710 | 12.6 | [9.9 - 15.8] | 20.8 | [18.6 - 23.2] | 13.0 | [10.6 - 15.7] | 8.32 |
| **'Butyrivibrio sp. XB500-5' (RAIS01000001-RAIS01000016)** | *Lachnobacterium bovis* DSM 14045 | 12.6 | [9.9 - 15.9] | 20.7 | [18.4 - 23.1] | 13.0 | [10.7 - 15.7] | 11.08 |
| **'Butyrivibrio sp. X503' (RAIQ01000001-RAIQ01000019)** | *Kineothrix alysoides* DSM 100556 | 12.6 | [9.9 - 15.8] | 20.6 | [18.4 - 23.0] | 13.0 | [10.6 - 15.7] | 0.48 |
| **'Butyrivibrio sp. XB500-5' (RAIS01000001-RAIS01000016)** | *Roseburia faecis* M72 | 12.6 | [9.9 - 15.9] | 20.5 | [18.3 - 22.9] | 13.0 | [10.7 - 15.8] | 0.53 |
| **'Butyrivibrio sp. XB500-5' (RAIS01000001-RAIS01000016)** | *Eubacterium ruminantium* ATCC 17233 | 12.8 | [10.1 - 16.1] | 20.4 | [18.2 - 22.8] | 13.2 | [10.8 - 15.9] | 5.19 |
| **'Butyrivibrio sp. CB08' (RAIR01000001-RAIR01000029)** | *Roseburia faecis* M72 | 12.6 | [10.0 - 15.9] | 20.2 | [18.0 - 22.6] | 13.0 | [10.7 - 15.8] | 0.74 |
| **'Butyrivibrio sp. CB08' (RAIR01000001-RAIR01000029)** | *Butyrivibrio hungatei* DSM 14810 | 17.3 | [14.3 - 20.9] | 20.1 | [17.9 - 22.5] | 17.1 | [14.5 - 20.0] | 3.8 |
| **'Butyrivibrio sp. X503' (RAIQ01000001-RAIQ01000019)** | **'Butyrivibrio sp. CB08' (RAIR01000001-RAIR01000029)** | 16.0 | [13.1 - 19.5] | 20.0 | [17.8 - 22.4] | 16.0 | [13.4 - 18.9] | 1.48 |
| **'Butyrivibrio sp. XB500-5' (RAIS01000001-RAIS01000016)** | *Merdimonas faecis* BR31 | 12.6 | [9.9 - 15.9] | 20.0 | [17.8 - 22.4] | 13.0 | [10.7 - 15.7] | 4.53 |
| **'Butyrivibrio sp. CB08' (RAIR01000001-RAIR01000029)** | **'Butyrivibrio sp. XB500-5' (RAIS01000001-RAIS01000016)** | 16.0 | [13.0 - 19.5] | 20.0 | [17.8 - 22.4] | 16.0 | [13.4 - 18.9] | 1.27 |

| Query | Subject | $d_0$ | C.I. $d_0$ | $d_4$ | C.I. $d_4$ | $d_6$ | C.I. $d_6$ | Diff. G+C Percent |
|---|---|---|---|---|---|---|---|---|
| **'Butyrivibrio sp. XB500-5' (RAIS01000001-RAIS01000016)** | *Enterocloster clostridioformis* ATCC 25537 | 12.5 | [9.9 - 15.8] | 20.0 | [17.8 - 22.4] | 12.9 | [10.6 - 15.7] | 6.53 |
| **'Butyrivibrio sp. XB500-5' (RAIS01000001-RAIS01000016)** | *Enterocloster bolteae* ATCC BAA-613 | 12.5 | [9.9 - 15.8] | 20.0 | [17.8 - 22.4] | 12.9 | [10.6 - 15.7] | 6.63 |
| **'Butyrivibrio sp. X503' (RAIQ01000001-RAIQ01000019)** | *Butyrivibrio hungatei* DSM 14810 | 17.0 | [14.0 - 20.6] | 20.0 | [17.8 - 22.4] | 16.8 | [14.2 - 19.8] | 2.31 |
| **'Butyrivibrio sp. XB500-5' (RAIS01000001-RAIS01000016)** | *Butyrivibrio hungatei* DSM 14810 | 17.0 | [14.0 - 20.5] | 19.9 | [17.7 - 22.3] | 16.8 | [14.2 - 19.7] | 2.52 |
| **'Butyrivibrio sp. X503' (RAIQ01000001-RAIQ01000019)** | *Eubacterium ruminantium* ATCC 17233 | 12.8 | [10.1 - 16.1] | 19.9 | [17.7 - 22.3] | 13.2 | [10.8 - 15.9] | 4.97 |
| **'Butyrivibrio sp. XB500-5' (RAIS01000001-RAIS01000016)** | *Eubacterium xylanophilum* ATCC 35991 | 12.6 | [10.0 - 15.9] | 19.9 | [17.7 - 22.3] | 13.0 | [10.7 - 15.8] | 2.58 |
| **'Butyrivibrio sp. CB08' (RAIR01000001-RAIR01000029)** | *Mesorhizobium loti* DSM 2626 | 12.5 | [9.8 - 15.8] | 19.9 | [17.7 - 22.4] | 12.9 | [10.6 - 15.6] | 18.66 |
| **'Butyrivibrio sp. X503' (RAIQ01000001-RAIQ01000019)** | *Eubacterium xylanophilum* ATCC 35991 | 12.7 | [10.0 - 15.9] | 19.8 | [17.6 - 22.2] | 13.0 | [10.7 - 15.8] | 2.37 |
| **'Butyrivibrio sp. X503' (RAIQ01000001-RAIQ01000019)** | *Eubacterium ventriosum* ATCC 27560 | 12.7 | [10.0 - 15.9] | 19.8 | [17.6 - 22.2] | 13.0 | [10.7 - 15.8] | 7.3 |
| **'Butyrivibrio sp. X503' (RAIQ01000001-RAIQ01000019)** | *Clostridium aminophilum* DSM 10710 | 12.6 | [9.9 - 15.8] | 19.7 | [17.5 - 22.1] | 13.0 | [10.6 - 15.7] | 8.53 |
| **'Butyrivibrio sp. X503' (RAIQ01000001-RAIQ01000019)** | *Mesorhizobium loti* DSM 2626 | 12.5 | [9.8 - 15.8] | 19.7 | [17.5 - 22.1] | 12.9 | [10.6 - 15.6] | 20.14 |
| **'Butyrivibrio sp. XB500-5' (RAIS01000001-RAIS01000016)** | *Lactobacillus rogosae* ATCC 27753 | 12.6 | [9.9 - 15.9] | 19.5 | [17.3 - 21.9] | 13.0 | [10.7 - 15.7] | 6.3 |
| **'Butyrivibrio sp. XB500-5' (RAIS01000001-RAIS01000016)** | *Kineothrix alysoides* DSM 100556 | 12.6 | [9.9 - 15.8] | 19.5 | [17.3 - 21.9] | 13.0 | [10.7 - 15.7] | 0.27 |
| **'Butyrivibrio sp. XB500-5' (RAIS01000001-RAIS01000016)** | *Eubacterium ventriosum* ATCC 27560 | 12.7 | [10.0 - 15.9] | 19.5 | [17.3 - 21.9] | 13.0 | [10.7 - 15.8] | 7.51 |
| **'Butyrivibrio sp. X503' (RAIQ01000001-RAIQ01000019)** | *Merdimonas faecis* BR31 | 12.6 | [9.9 - 15.9] | 19.4 | [17.3 - 21.8] | 13.0 | [10.7 - 15.7] | 4.74 |
| **'Butyrivibrio sp. CB08' (RAIR01000001-RAIR01000029)** | *Blautia luti* DSM 14534 | 12.6 | [10.0 - 15.9] | 19.4 | [17.2 - 21.8] | 13.0 | [10.7 - 15.8] | 0.85 |
| **'Butyrivibrio sp. X503' (RAIQ01000001-RAIQ01000019)** | *Blautia luti* DSM 14534 | 12.6 | [9.9 - 15.9] | 19.2 | [17.1 - 21.6] | 13.0 | [10.7 - 15.8] | 0.64 |
| **'Butyrivibrio sp. X503' (RAIQ01000001-RAIQ01000019)** | *Clostridium nexile* DSM 1787 | 12.6 | [9.9 - 15.9] | 19.2 | [17.0 - 21.6] | 13.0 | [10.7 - 15.7] | 2.12 |
| **'Butyrivibrio sp. X503' (RAIQ01000001-RAIQ01000019)** | *Lactobacillus rogosae* ATCC 27753 | 12.6 | [9.9 - 15.9] | 19.1 | [17.0 - 21.5] | 13.0 | [10.7 - 15.8] | 6.08 |
| **'Butyrivibrio sp. CB08' (RAIR01000001-RAIR01000029)** | *Eubacterium xylanophilum* ATCC 35991 | 12.7 | [10.0 - 16.0] | 19.0 | [16.8 - 21.4] | 13.1 | [10.7 - 15.8] | 3.86 |
| **'Butyrivibrio sp. XB500-5' (RAIS01000001-RAIS01000016)** | *Bacteroides galacturonicus* DSM 3978 | 12.6 | [10.0 - 15.9] | 18.9 | [16.8 - 21.3] | 13.0 | [10.7 - 15.8] | 6.18 |

| Query | Subject | $d_0$ | C.I. $d_0$ | $d_4$ | C.I. $d_4$ | $d_6$ | C.I. $d_6$ | Diff. G+C Percent |
|---|---|---|---|---|---|---|---|---|
| **'Butyrivibrio sp. CB08' (RAIR01000001-RAIR01000029)** | *Bacteroides galacturonicus* DSM 3978 | 12.7 | [10.0 - 15.9] | 18.9 | [16.7 - 21.3] | 13.0 | [10.7 - 15.8] | 7.45 |
| **'Butyrivibrio sp. XB500-5' (RAIS01000001-RAIS01000016)** | *Butyrivibrio crossotus* DSM 2876 | 12.6 | [10.0 - 15.9] | 18.9 | [16.7 - 21.3] | 13.0 | [10.7 - 15.8] | 4.69 |
| **'Butyrivibrio sp. XB500-5' (RAIS01000001-RAIS01000016)** | *Eisenbergiella tayi* DSM 26961 | 12.5 | [9.9 - 15.8] | 18.8 | [16.6 - 21.2] | 12.9 | [10.6 - 15.7] | 4.36 |
| **'Butyrivibrio sp. CB08' (RAIR01000001-RAIR01000029)** | *Clostridium aminophilum* DSM 10710 | 12.6 | [9.9 - 15.8] | 18.8 | [16.6 - 21.2] | 13.0 | [10.6 - 15.7] | 7.05 |
| **'Butyrivibrio sp. XB500-5' (RAIS01000001-RAIS01000016)** | *Clostridium nexile* DSM 1787 | 12.6 | [9.9 - 15.9] | 18.7 | [16.5 - 21.0] | 13.0 | [10.7 - 15.7] | 2.33 |
| **'Butyrivibrio sp. CB08' (RAIR01000001-RAIR01000029)** | *Eubacterium oxidoreducens* DSM 3217 | 12.6 | [9.9 - 15.9] | 18.7 | [16.5 - 21.1] | 13.0 | [10.7 - 15.7] | 3.84 |
| **'Butyrivibrio sp. X503' (RAIQ01000001-RAIQ01000019)** | *Bacteroides galacturonicus* DSM 3978 | 12.6 | [10.0 - 15.9] | 18.6 | [16.4 - 21.0] | 13.0 | [10.7 - 15.8] | 5.97 |
| **'Butyrivibrio sp. CB08' (RAIR01000001-RAIR01000029)** | *Lacrimispora aerotolerans* DSM 5434 | 12.5 | [9.9 - 15.8] | 18.6 | [16.5 - 21.0] | 12.9 | [10.6 - 15.7] | 1.33 |
| **'Butyrivibrio sp. CB08' (RAIR01000001-RAIR01000029)** | *Clostridium nexile* DSM 1787 | 12.6 | [9.9 - 15.8] | 18.5 | [16.4 - 20.9] | 13.0 | [10.7 - 15.7] | 3.6 |
| **'Butyrivibrio sp. X503' (RAIQ01000001-RAIQ01000019)** | *Anaerocolumna aminovalerica* DSM 1283 | 12.5 | [9.9 - 15.8] | 18.5 | [16.4 - 20.9] | 12.9 | [10.6 - 15.7] | 7.22 |
| **'Butyrivibrio sp. CB08' (RAIR01000001-RAIR01000029)** | *Lactobacillus rogosae* ATCC 27753 | 12.6 | [10.0 - 15.9] | 18.5 | [16.3 - 20.9] | 13.0 | [10.7 - 15.8] | 7.57 |
| **'Butyrivibrio sp. XB500-5' (RAIS01000001-RAIS01000016)** | *Blautia luti* DSM 14534 | 12.6 | [9.9 - 15.9] | 18.4 | [16.3 - 20.8] | 13.0 | [10.7 - 15.8] | 0.43 |
| **'Butyrivibrio sp. CB08' (RAIR01000001-RAIR01000029)** | *Eisenbergiella tayi* DSM 26961 | 12.5 | [9.9 - 15.8] | 18.0 | [15.9 - 20.4] | 12.9 | [10.6 - 15.7] | 3.09 |
| **'Butyrivibrio sp. CB08' (RAIR01000001-RAIR01000029)** | *Butyrivibrio crossotus* DSM 2876 | 12.7 | [10.0 - 16.0] | 18.0 | [15.9 - 20.4] | 13.1 | [10.7 - 15.8] | 5.96 |
| **'Butyrivibrio sp. X503' (RAIQ01000001-RAIQ01000019)** | *Butyrivibrio crossotus* DSM 2876 | 12.6 | [9.9 - 15.9] | 18.0 | [15.9 - 20.4] | 13.0 | [10.7 - 15.8] | 4.48 |
| **'Butyrivibrio sp. XB500-5' (RAIS01000001-RAIS01000016)** | *Anaerocolumna aminovalerica* DSM 1283 | 12.5 | [9.9 - 15.8] | 18.0 | [15.9 - 20.4] | 12.9 | [10.6 - 15.7] | 7.43 |
| **'Butyrivibrio sp. XB500-5' (RAIS01000001-RAIS01000016)** | *Lacrimispora aerotolerans* DSM 5434 | 12.6 | [9.9 - 15.8] | 17.8 | [15.7 - 20.2] | 13.0 | [10.6 - 15.7] | 0.05 |
| **'Butyrivibrio sp. CB08' (RAIR01000001-RAIR01000029)** | *Anaerocolumna aminovalerica* DSM 1283 | 12.5 | [9.9 - 15.8] | 17.7 | [15.5 - 20.0] | 12.9 | [10.6 - 15.7] | 8.7 |
| **'Butyrivibrio sp. X503' (RAIQ01000001-RAIQ01000019)** | *Eisenbergiella tayi* DSM 26961 | 12.6 | [9.9 - 15.8] | 17.3 | [15.2 - 19.6] | 13.0 | [10.6 - 15.7] | 4.57 |
| **'Butyrivibrio sp. X503' (RAIQ01000001-RAIQ01000019)** | *Lacrimispora aerotolerans* DSM 5434 | 12.6 | [9.9 - 15.8] | 17.2 | [15.1 - 19.5] | 13.0 | [10.6 - 15.7] | 0.16 |
| **'Butyrivibrio sp. X503' (RAIQ01000001-RAIQ01000019)** | *Chryseobacterium bernardetii* G229 | 12.5 | [9.9 - 15.8] | 15.6 | [13.6 - 17.9] | 12.9 | [10.6 - 15.7] | 5.89 |

| Query | Subject | $d_0$ | C.I. $d_0$ | $d_4$ | C.I. $d_4$ | $d_6$ | C.I. $d_6$ | Diff. G+C Percent |
|---|---|---|---|---|---|---|---|---|
| **'Butyrivibrio sp. XB500-5' (RAIS01000001-RAIS01000016)** | *Chryseobacterium bernardetii* G229 | 12.5 | [9.9 - 15.8] | 15.5 | [13.5 - 17.8] | 12.9 | [10.6 - 15.7] | 6.1 |
| **'Butyrivibrio sp. CB08' (RAIR01000001-RAIR01000029)** | *Chryseobacterium bernardetii* G229 | 12.5 | [9.9 - 15.8] | 15.4 | [13.4 - 17.7] | 12.9 | [10.6 - 15.7] | 7.37 |

#### Table 4: Strains in your dataset

Joint dataset of automatically determined closest type strains (if this mode was chosen), manually selected type strains (if selected accordingly) and the provided user strains, if provided (marked in yellow).

| Strain | Authority | Other deposits | Synonyms | Base pairs | Percent G+C | No. proteins | Goldstamp | Bioproject accession | Biosample accession | Assembly accession | IMG OID |
|---|---|---|---|---|---|---|---|---|---|---|---|
| *Eisenbergiella tayi* DSM 26961 | Amir et al. 2014 | LMG 27400; ATCC BAA-2558; B086562 | *Eisenbergiella tayi* | 7552 069 | 46.8 | 6418 | Gp0371963 | PRJNA224116 | SAMN05941924 | GCF_001881565 | |
| *Merdimonas faecis* BR31 | Seo et al. 2017 | KCTC 15482; JCM 30748 | *Merdimonas faecis* | 3318 223 | 47.0 | 3203 | Gp0314539 | PRJNA224116 | SAMN05727926 | GCF_001754075 | |
| *Lacrimispora celerecrescens* 18A | (Palop et al. 1989) Haas and Blanchard 2020 | CECT 954; DSM 5628; ATCC 49205 | *Clostridium celerecrescens*; *Lacrimispora celerecrescens* | 5272 838 | 43.9 | 4625 | Gp0032364 | PRJNA187124 | SAMN02745763 | GCA_002797975 | |
| *Enterocloster clostridioformis* ATCC 25537 | (Burri and Ankersmit 1906) Haas and Blanchard 2020 | BCRC 14545; CCRC 14545; CCUG 16791; DSM 933; NCTC 11224; JCM 1291; CIP 104318; NCIMB 11018; VPI 316 | *Clostridium clostridioforme*; *Enterocloster clostridioformis* | 5465 381 | 49.0 | 5166 | Gp0088045 | PRJEB17285 | SAMN05660211 | GCA_900113155 | |
| *Blautia luti* DSM 14534 | (Simmering et al. 2002) Liu et al. 2008 | CCUG 45635; BlnIX | *Blautia luti*; *Ruminococcus luti* | 3704 127 | 42.9 | 3244 | | PRJNA590133 | SAMN13319600 | GCA_009707925 | |
| *Eubacterium xylanophilum* ATCC 35991 | van Gylswyk and van der Toorn 1985 | X6C58 | *Eubacterium xylanophilum* | 2574 620 | 39.8 | 2188 | Gp0032280 | PRJNA204102 | SAMN02584969 | GCA_000518685 | 2545555857 |
| *Dorea phocaeensis* Marseille-P4003 | Takakura et al. 2019 | | *Dorea phocaeensis* | 2462 468 | 43.2 | 2298 | | PRJEB22820 | SAMEA104415757 | GCA_900240315 | |
| *Anaerostipes rhamnosivorans* 1y-2 | Bui et al. 2014 | KCTC 15316; DSM 26241 | *Anaerostipes rhamnosivorans* | 3588 860 | 44.5 | 3556 | Gp0441668 | PRJNA540423 | SAMN11535143 | GCA_005280655 | |

| Strain | Authority | Other deposits | Synonyms | Base pairs | Percent G+C | No. proteins | Goldstamp | Bioproject accession | Biosample accession | Assembly accession | IMG OID |
|---|---|---|---|---|---|---|---|---|---|---|---|
| *Clostridium aminophilum* DSM 10710 | Paster et al. 1993 | ATCC 49906; F | *Clostridium aminophilum* | 3116 031 | 50.7 | 2570 | Gp0013480 | PRJNA234874 | SAMN02745353 | GCA_000711825 | 2565956525 |
| *Butyrivibrio fibrisolvens* DSM 3071 | Bryant and Small 1956 | ATCC 19171 | *Butyrivibrio fibrisolvens* | 4837 257 | 39.7 | 4062 | Gp0013621 | PRJNA245644 | SAMN02745229 | GCA_900129945 | 2585428068 |
| *Anaerocolumna jejuensis* DSM 15929 | (Jeong et al. 2004) Ueki et al. 2016 | KCTC 5026; HY-35-12; IMSNU 40003 | *Anaerocolumna jejuensis*; *Clostridium jejuense* | 6620 253 | 40.6 | 5676 | Gp0013560 | PRJNA245653 | SAMN02745136 | GCA_900142215 | 2585428173 |
| *Chryseobacterium bernardetii* G229 | Holmes et al. 2013 emend. Kim et al. 2016 | CCUG 60564; NCTC 13530; CDC G229; CL318/82 | *Chryseobacterium bernardetii* | 5318 634 | 36.3 | 4844 | Gp0389805 | PRJNA224116 | SAMN10343177 | GCA_003815975 | |
| *Butyrivibrio hungatei* DSM 14810 | Kopecný et al. 2003 | ATCC BAA-456; JK 615 | *Butyrivibrio hungatei* | 3391 547 | 39.9 | 3030 | Gp0013622 | PRJNA245645 | SAMN02745247 | GCA_900143205 | 2582580726 |
| *Roseburia faecis* M72 | Duncan et al. 2006 | DSM 16840; M72/1; NCIMB 14031 | *Roseburia faecis* | 3317 798 | 43.0 | 3168 | Gp0142292 | PRJEB9321 | SAMEA1710514 | GCA_001406815 | |
| *Eubacterium ruminantium* ATCC 17233 | Bryant 1959 | DSM 20704; GA 195 | *Eubacterium ruminantium* | 2840 588 | 37.2 | 2488 | Gp0088735 | PRJNA245567 | SAMN02745110 | GCA_900167085 | 2585428142 |
| *Lactobacillus rogosae* ATCC 27753 | Holdeman and Moore 1974 | VPI C37-38 | *Lactobacillus rogosae* | 2817 405 | 36.1 | 2497 | Gp0099458 | PRJNA257876 | SAMN02982992 | GCA_900112995 | 2597490368 |
| *Mesorhizobium loti* DSM 2626 | (Jarvis et al. 1982) Jarvis et al. 1997 emend. Hameed et al. 2015 | LMG 6125; CCUG 27878; ATCC 700743; JCM 21464; IFO 14779; NBRC 14779; HAMBI 1129; NZP 2213 | *Mesorhizobium loti*; *Rhizobium loti* | 7451 510 | 62.4 | 7264 | Gp0251913 | PRJNA442649 | SAMN08775555 | GCA_003148495 | 2756170246 |
| *Bacteroides galacturonicus* DSM 3978 | Jensen and Canale-Parola 1987 | ATCC 43244; N6 | *Bacteroides galacturonicus* | 2924 074 | 36.2 | 2658 | Gp0251964 | PRJNA439857 | SAMN08769267 | GCA_003096855 | 2757320519 |

| Strain | Authority | Other deposits | Synonyms | Base pairs | Percent G+C | No. proteins | Goldstamp | Bioproject accession | Biosample accession | Assembly accession | IMG OID |
|---|---|---|---|---|---|---|---|---|---|---|---|
| *Anaerocolumna aminovalerica* DSM 1283 | (Hardman and Stadtman 1960) Ueki et al. 2016 | ATCC 13725; JCM 11016; CIP 104304 | *Anaerocolumna aminovalerica*; *Clostridium aminovalericum* | 4624 034 | 35.0 | 4045 | Gp0116495 | PRJNA303727 | SAMN04489757 | GCA_900115365 | 2634166351 |
| *Butyrivibrio proteoclasticus* B316 | (Attwood et al. 1996) Moon et al. 2008 | DSM 14932; ATCC 51982 | *Butyrivibrio proteoclasticus*; *Clostridium proteoclasticum* | 4404 877 | 40.0 | 3811 | Gp0003213 | PRJNA29153 | SAMN02602973 | GCA_000145035 | 648028012 |
| *Eubacterium ventriosum* ATCC 27560 | (Tissier 1908) Prévot 1938 | DSM 3988 | *Eubacterium ventriosum* | 2869 695 | 34.9 | 2802 | Gp0000913 | PRJNA18159 | SAMN00627091 | GCA_000153885 | 640963022 |
| *Butyrivibrio crossotus* DSM 2876 | Moore et al. 1976 | ATCC 29175; VPI T9-40A | *Butyrivibrio crossotus* | 2480 245 | 37.7 | 2529 | Gp0003344 | PRJNA28999 | SAMN00008799 | GCA_000156015 | 645951834 |
| *Roseburia intestinalis* L1-82 | Duncan et al. 2002 emend. Duncan et al. 2006 | DSM 14610; NCIMB 13810 | *Roseburia intestinalis* | 4379 102 | 42.6 | 4707 | Gp0003375 | PRJNA30005 | SAMN00008856 | GCA_000156535 | 2562617159 |
| *Eubacterium oxidoreducens* DSM 3217 | Krumholz and Bryant 1986 | ATCC 43585; G41 | *Eubacterium oxidoreducens* | 2910 102 | 39.9 | 2629 | Gp0087953 | PRJEB16259 | SAMN02910417 | GCA_900104415 | |
| *Lachnobacterium bovis* DSM 14045 | Whitford et al. 2001 | ATCC BAA-151; LRC 5382; YZ 87 | *Lachnobacterium bovis* | 2709 759 | 31.3 | 2399 | Gp0095053 | PRJEB16629 | SAMN02910414 | GCA_900107245 | |
| *Enterocloster bolteae* ATCC BAA-613 | (Song et al. 2003) Haas and Blanchard 2020 | 16351; CCUG 46953; DSM 15670; WAL 16351 | *Clostridium bolteae*; *Enterocloster bolteae* | 1311 3976 | 49.1 | 7284 | Gp0000896 | PRJNA18165 | SAMN00627070 | GCA_000154365 | 641380428 |
| *Lacrimispora aerotolerans* DSM 5434 | (van Gylswyk and van der Toorn 1987) Haas and Blanchard 2020 | ATCC 43524; strain X8A62 | *Clostridium aerotolerans*; *Lacrimispora aerotolerans* | 4732 373 | 42.4 | 4271 | Gp0046989 | PRJNA223500 | SAMN02743875 | GCA_000687555 | 2558860131 |
| *Clostridium nexile* DSM 1787 | Holdeman and Moore 1974 | ATCC 27757 | *Clostridium nexile* | 3861 016 | 40.1 | 4239 | Gp0003367 | PRJNA28659 | SAMN00000730 | GCA_000156035 | 642979369 |
| *Kineothrix alysoides* KNHs209 | Haas and Blanchard 2017 | DSM 100556; ATCC TSD-26; KNHs209 | *Kineothrix alysoides* | 4676 087 | 42.7 | 4033 | Gp0040515 | PRJNA224116 | SAMN02910261 | GCF_000732725 | |

| Strain | Authority | Other deposits | Synonyms | Base pairs | Percent G+C | No. proteins | Goldstamp | Bioproject accession | Biosample accession | Assembly accession | IMG OID |
|---|---|---|---|---|---|---|---|---|---|---|---|
| *Kineothrix alysoides* DSM 100556 | Haas and Blanchard 2017 | DSM 100556; ATCC TSD-26; KNHs209 | *Kineothrix alysoides* | 4611 545 | 42.7 | 4122 | Gp0290642 | PRJNA500716 | SAMN10362787 | GCA_004345255 | 2788499853 |
| *Eubacterium cellulosolvens* 6 | (Bryant et al. 1958) Holdeman and Moore 1972 emend. van Gylswyk and van der Toorn 1986 | ATCC 43171; JCM 9499; van Gylswyk and Hoffman strain 6 | *Eubacterium cellulosolvens* | 3383 756 | 48.2 | 2749 | Gp0003902 | PRJNA45821 | SAMN02256545 | GCA_000183525 | 2509276007 |
| *Blautia hansenii* DSM 20583 | (Holdeman and Moore 1974) Liu et al. 2008 | ATCC 27752; JCM 14655; CIP 104219 | *Blautia hansenii*; *Ruminococcus hansenii*; *Streptococcus hansenii* | 3052 612 | 39.0 | 3171 | Gp0003378 | PRJNA30021 | SAMN00008797 | GCA_000156675 | 2562617096 |
| *Coprococcus eutactus* ATCC 27759 | Holdeman and Moore 1974 | | *Coprococcus eutactus* | 3101 325 | 43.1 | 2982 | Gp0000905 | PRJNA18187 | SAMN00627069 | GCA_000154425 | 641380422 |
| 'Butyrivibrio sp. X503' (RAIQ01000001-RAIQ01000019) | | | | 3239 716 | 42.2 | 2991 | | | | | |
| 'Butyrivibrio sp. CB08' (RAIR01000001-RAIR01000029) | | | | 3544 402 | 43.7 | 3280 | | | | | |
| 'Butyrivibrio sp. XB500-5' (RAIS01000001-RAIS01000016) | | | | 3273 773 | 42.4 | 2974 | | | | | |

#### Methods, Results and References

The genome sequence data were uploaded to the Type (Strain) Genome Server (TYGS), a free bioinformatics platform available under https://tygs.dsmz.de, for a whole genome-based taxonomic analysis [1]. The results were provided by the TYGS on 2020-04-09. In brief, the TYGS analysis was subdivided into the following steps:

##### Determination of closely related type strains

Determination of closest type strain genomes was done in two complementary ways: First, all user genomes were compared against all type strain genomes available in the TYGS database via the MASH algorithm, a fast approximation of intergenomic relatedness [2], and, the ten type strains with the smallest MASH distances chosen per user genome. Second, an additional set of ten closely related type strains was determined via the 16S rDNA gene sequences. These were extracted from the user genomes using RNAmmer [3] and each sequence was subsequently BLASTed [4] against the 16S rDNA gene sequence of each of the currently 11354 type strains available in the TYGS database. This was used as a proxy to find the best 50 matching type strains (according to the bitscore) for each user genome and to subsequently calculate precise distances using the Genome BLAST Distance Phylogeny approach (GBDP) under the algorithm 'coverage' and distance formula $d_5$ [5]. These distances were finally used to determine the 10 closest type strain genomes for each of the user genomes.

##### Pairwise comparison of genome sequences

All pairwise comparisons among the set of genomes were conducted using GBDP and accurate intergenomic distances inferred under the algorithm 'trimming' and distance formula $d_5$ [5]. 100 distance replicates were calculated each. Digital DDH values and confidence intervals were calculated using the recommended settings of the GGDC 2.1 [5].

##### Phylogenetic inference

The resulting intergenomic distances were used to infer a balanced minimum evolution tree with branch support via FASTME 2.1.4 including SPR postprocessing [6]. Branch support was inferred from 100 pseudo-bootstrap replicates each. The trees were rooted at the midpoint [7] and visualized with PhyD3 [8].

##### Type-based species and subspecies clustering

The type-based species clustering using a 70% dDDH radius around each of the 33 type strains was done as previously described [1]. The resulting groups are shown in Table 1 and 4. Subspecies clustering was done using a 79% dDDH threshold as previously introduced [9].

#### Results

##### Type-based species and subspecies clustering

The resulting species and subspecies clusters are listed in Table 4, whereas the taxonomic identification of the query strains is found in Table 1. Briefly, the clustering yielded 34 species clusters and the provided query strains were assigned to 3 of these. Moreover, user strains were located in 3 of 34 subspecies clusters.

##### Figure caption genome tree

**Figure 1.** Tree inferred with FastME 2.1.6.1 [6] from GBDP distances calculated from genome sequences. The branch lengths are scaled in terms of GBDP distance formula $d_5$. The numbers above branches are GBDP pseudo-bootstrap support values > 60 % from 100 replications, with an average branch support of 21.9 %. The tree was rooted at the midpoint [7].

##### Figure caption SSU tree

**Figure 2.** Tree inferred with FastME 2.1.6.1 [6] from GBDP distances calculated from 16S rDNA gene sequences. The branch lengths are scaled in terms of GBDP distance formula $d_5$. The numbers above branches are GBDP pseudo-bootstrap support values > 60 % from 100 replications, with an average branch support of 61.3 %. The tree was rooted at the midpoint [7].

#### References


[1] Meier-Kolthoff JP, Göker M. TYGS is an automated high-throughput platform for state-of-the-art genome-based taxonomy. Nat. Commun. 2019;10: 2182. DOI: 10.1038/s41467-019-10210-3

[2] Ondov BD, Treangen TJ, Melsted P, et al. Mash: Fast genome and metagenome distance estimation using MinHash. Genome Biol 2016;17: 1–14. DOI: 10.1186/s13059-016-0997-x

[3] Lagesen K, Hallin P. RNAmmer: consistent and rapid annotation of ribosomal RNA genes. Nucleic Acids Res. Oxford Univ Press; 2007;35: 3100–3108. DOI: 10.1093/nar/gkm160

[4] Camacho C, Coulouris G, Avagyan V, Ma N, Papadopoulos J, Bealer K, et al. BLAST+: architecture and applications. BMC Bioinformatics. 2009;10: 421. DOI: 10.1186/1471-2105-10-421

[5] Meier-Kolthoff JP, Auch AF, Klenk H-P, Göker M. Genome sequence-based species delimitation with confidence intervals and improved distance functions. BMC Bioinformatics. 2013;14: 60. DOI: 10.1186/1471-2105-14-60

[6] Lefort V, Desper R, Gascuel O. FastME 2.0: A comprehensive, accurate, and fast distance-based phylogeny inference program. Mol Biol Evol. 2015;32: 2798–2800. DOI: 10.1093/molbev/msv150

[7] Farris JS. Estimating phylogenetic trees from distance matrices. Am Nat. 1972;106: 645–667.

[8] Kreft L, Botzki A, Coppens F, Vandepoele K, Van Bel M. PhyD3: A phylogenetic tree viewer with extended phyloXML support for functional genomics data visualization. Bioinformatics. 2017;33: 2946–2947. DOI: 10.1093/bioinformatics/btx324

[9] Meier-Kolthoff JP, Hahnke RL, Petersen J, Scheuner C, Michael V, Fiebig A, et al. Complete genome sequence of DSM 30083$^T$, the type strain (U5/41$^T$) of *Escherichia coli*, and a proposal for delineating subspecies in microbial taxonomy. Stand Genomic Sci. 2014;9: 2. DOI: 10.1186/1944-3277-9-2
