## Supplementary file 8 for "Genomic architecture of three newly isolated unclassified *Butyrivibrio* species elucidate their potential role in the rumen ecosystem"

### dbCAN meta server: automated CAZyme annotation

(index.php)

Home (index.php) | Annotate (blast.php) | Download (download/) | Help (help.php) | About us (about.php)

[illegible]79 (https://doi.org/10.1093/nar/gk[illegible]79)

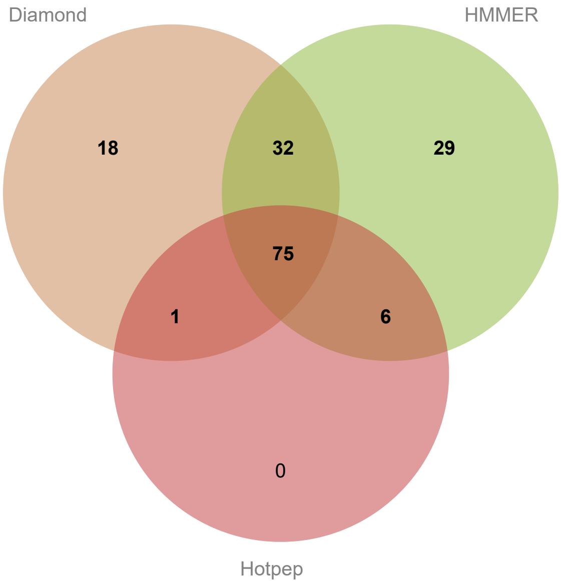


Result of job: 2020040800435

Overview | HMMER | DIAMOND | Hotpep | CGC-Finder

**Download SignalP output (data/blast/2020040800435/signalp.out)** **Download Prodigal predictions (data/blast/2020040800435/uniInput)** **Download this table** (keep those with # of Tools >=2 will give you best result; and use dbCAN domain assignment is recommended) **(data/blast/2020040800435/overview.txt) ? (help.php#over)**

**Show** All ▼ **entries** **Search:**

| Gene ID | # of Tools | HMMER | DIAMOND | Hotpep | Signal Peptide |
|---|---|---|---|---|---|
| scaffold10\|size62189_16 (domain.php?jobid=2020040800435&gene=scaffold10%7Csize62189_16) | 3 | GT5 (http://www.cazy.org/GT5.html)(3-476) | GT5 (http://www.cazy.org/GT5.html) | GT5 (http://www.cazy.org/GT5.html) | N |
| scaffold11\|size56834_16 (domain.php?jobid=2020040800435&gene=scaffold11%7Csize56834_16) | 3 | GH3 (http://www.cazy.org/GH3.html)(520-734) | GH3 (http://www.cazy.org/GH3.html) | GH3 (http://www.cazy.org/GH3.html) | N |
| scaffold12\|size54477_15 (domain.php?jobid=2020040800435&gene=scaffold12%7Csize54477_15) | 2 | GH73 (http://www.cazy.org/GH73.html)(15-150) | N | GH73 (http://www.cazy.org/GH73.html) | N |
| scaffold13\|size40669_29 (domain.php?jobid=2020040800435&gene=scaffold13%7Csize40669_29) | 2 | GH73 (http://www.cazy.org/GH73.html)(15-150) | N | GH73 (http://www.cazy.org/GH73.html) | N |
| scaffold14\|size32532_16 (domain.php?jobid=2020040800435&gene=scaffold14%7Csize32532_16) | 3 | GH43_12 (http://www.cazy.org/GH43_12.html)(129-405) | GH43_12 (http://www.cazy.org/GH43_12.html) | GH43 (http://www.cazy.org/GH43.html) | N |
| scaffold14\|size32532_32 (domain.php?jobid=2020040800435&gene=scaffold14%7Csize32532_32) | 1 | GH114 (http://www.cazy.org/GH114.html)(41-144) | N | N | Y (1-31) |
| scaffold16\|size19705_9 | 1 | N | GH28 (http://www.cazy.org/GH28.html) | N | N |
| scaffold1\|size1187143_114 (domain.php?jobid=2020040800435&gene=scaffold1%7Csize1187143_114) | 2 | GT2_Glycos_transf_2 (http://www.cazy.org/GT2_Glycos_transf_2.html)(55-228) | GT2 (http://www.cazy.org/GT2.html) | N | N |
| scaffold1\|size1187143_123 (domain.php?jobid=2020040800435&gene=scaffold1%7Csize1187143_123) | 2 | CBM48 (http://www.cazy.org/CBM48.html)(127-213)+GH13_9 (http://www.cazy.org/GH13_9.html)(279-572) | CBM48 (http://www.cazy.org/CBM48.html)+GH13_9 (http://www.cazy.org/GH13_9.html) | N | N |

| Gene ID | # of Tools | HMMER | DIAMOND | Hotpep | Signal Peptide |
|---|---|---|---|---|---|
| scaffold1\|size1187143_145 (domain.php?jobid=2020040800435&gene=scaffold1%7Csize1187143_145) | 3 | GH2 (http://www.cazy.org/GH2.html)(30-905) | GH2 (http://www.cazy.org/GH2.html) | GH2 (http://www.cazy.org/GH2.html) | N |
| scaffold1\|size1187143_183 (domain.php?jobid=2020040800435&gene=scaffold1%7Csize1187143_183) | 3 | GH43_10 (http://www.cazy.org/GH43_10.html)(12-283) | GH43_10 (http://www.cazy.org/GH43_10.html) | GH43 (http://www.cazy.org/GH43.html) | N |
| scaffold1\|size1187143_184 (domain.php?jobid=2020040800435&gene=scaffold1%7Csize1187143_184) | 2 | CE12 (http://www.cazy.org/CE12.html)(4-213) | N | CE12 (http://www.cazy.org/CE12.html) | N |
| scaffold1\|size1187143_220 (domain.php?jobid=2020040800435&gene=scaffold1%7Csize1187143_220) | 1 | CE13 (http://www.cazy.org/CE13.html)(64-256) | N | N | N |
| scaffold1\|size1187143_223 (domain.php?jobid=2020040800435&gene=scaffold1%7Csize1187143_223) | 3 | GH120 (http://www.cazy.org/GH120.html)(296-386) | GH120 (http://www.cazy.org/GH120.html) | GH120 (http://www.cazy.org/GH120.html) | N |
| scaffold1\|size1187143_263 (domain.php?jobid=2020040800435&gene=scaffold1%7Csize1187143_263) | 2 | GH3 (http://www.cazy.org/GH3.html)(173-380) | GH3 (http://www.cazy.org/GH3.html) | N | N |
| scaffold1\|size1187143_265 | 1 | N | GH13_11 (http://www.cazy.org/GH13_11.html) | N | N |
| scaffold1\|size1187143_268 (domain.php?jobid=2020040800435&gene=scaffold1%7Csize1187143_268) | 3 | GT2_Glycos_transf_2 (http://www.cazy.org/GT2_Glycos_transf_2.html)(5-111) | GT2 (http://www.cazy.org/GT2.html) | GT2 (http://www.cazy.org/GT2.html) | N |
| scaffold1\|size1187143_309 (domain.php?jobid=2020040800435&gene=scaffold1%7Csize1187143_309) | 3 | GH13_20 (http://www.cazy.org/GH13_20.html)(34-321) | GH13 (http://www.cazy.org/GH13.html) | GH13 (http://www.cazy.org/GH13.html) | N |
| scaffold1\|size1187143_348 (domain.php?jobid=2020040800435&gene=scaffold1%7Csize1187143_348) | 3 | GH161 (http://www.cazy.org/GH161.html)(7-1013) | GH161 (http://www.cazy.org/GH161.html) | GH161 (http://www.cazy.org/GH161.html) | N |
| scaffold1\|size1187143_349 (domain.php?jobid=2020040800435&gene=scaffold1%7Csize1187143_349) | 3 | GH3 (http://www.cazy.org/GH3.html)(558-766) | GH3 (http://www.cazy.org/GH3.html) | GH3 (http://www.cazy.org/GH3.html) | N |
| scaffold1\|size1187143_357 (domain.php?jobid=2020040800435&gene=scaffold1%7Csize1187143_357) | 3 | GT2_Glycos_transf_2 (http://www.cazy.org/GT2_Glycos_transf_2.html)(3-115) | GT2 (http://www.cazy.org/GT2.html) | GT2 (http://www.cazy.org/GT2.html) | N |
| scaffold1\|size1187143_382 (domain.php?jobid=2020040800435&gene=scaffold1%7Csize1187143_382) | 3 | GH31 (http://www.cazy.org/GH31.html)(157-582) | GH31 (http://www.cazy.org/GH31.html) | GH31 (http://www.cazy.org/GH31.html) | N |
| scaffold1\|size1187143_383 (domain.php?jobid=2020040800435&gene=scaffold1%7Csize1187143_383) | 3 | GH13_21 (http://www.cazy.org/GH13_21.html)(182-488) | GH13_21 (http://www.cazy.org/GH13_21.html) | GH13 (http://www.cazy.org/GH13.html) | N |
| scaffold1\|size1187143_43 (domain.php?jobid=2020040800435&gene=scaffold1%7Csize1187143_43) | 1 | CE4 (http://www.cazy.org/CE4.html)(19-147) | N | N | N |
| scaffold1\|size1187143_488 (domain.php?jobid=2020040800435&gene=scaffold1%7Csize1187143_488) | 3 | CE4 (http://www.cazy.org/CE4.html)(118-241) | CE4 (http://www.cazy.org/CE4.html) | CE4 (http://www.cazy.org/CE4.html) | N |
| scaffold1\|size1187143_491 (domain.php?jobid=2020040800435&gene=scaffold1%7Csize1187143_491) | 3 | GH9 (http://www.cazy.org/GH9.html)(89-542) | GH9 (http://www.cazy.org/GH9.html) | GH9 (http://www.cazy.org/GH9.html) | N |

| Gene ID | # of Tools | HMMER | DIAMOND | Hotpep | Signal Peptide |
|---|---|---|---|---|---|
| scaffold1\|size1187143_501 (domain.php?jobid=2020040800435&gene=scaffold1%7Csize1187143_501) | 1 | CBM2 (http://www.cazy.org/CBM2.html)(59-155) | N | N | N |
| scaffold1\|size1187143_512 (domain.php?jobid=2020040800435&gene=scaffold1%7Csize1187143_512) | 3 | GH13_31 (http://www.cazy.org/GH13_31.html)(32-370) | GH13_31 (http://www.cazy.org/GH13_31.html) | GH13 (http://www.cazy.org/GH13.html) | N |
| scaffold1\|size1187143_513 (domain.php?jobid=2020040800435&gene=scaffold1%7Csize1187143_513) | 2 | GT4 (http://www.cazy.org/GT4.html)(202-365) | GT4 (http://www.cazy.org/GT4.html) | N | N |
| scaffold1\|size1187143_542 (domain.php?jobid=2020040800435&gene=scaffold1%7Csize1187143_542) | 3 | GH2 (http://www.cazy.org/GH2.html)(16-461) | GH2 (http://www.cazy.org/GH2.html) | GH2 (http://www.cazy.org/GH2.html) | N |
| scaffold1\|size1187143_547 (domain.php?jobid=2020040800435&gene=scaffold1%7Csize1187143_547) | 3 | GH43_33 (http://www.cazy.org/GH43_33.html)(46-329) | GH43_33 (http://www.cazy.org/GH43_33.html) | GH117 (http://www.cazy.org/GH117.html) | Y (1-29) |
| scaffold1\|size1187143_567 (domain.php?jobid=2020040800435&gene=scaffold1%7Csize1187143_567) | 3 | GH3 (http://www.cazy.org/GH3.html)(29-249) | GH3 (http://www.cazy.org/GH3.html) | GH3 (http://www.cazy.org/GH3.html) | N |
| scaffold1\|size1187143_573 (domain.php?jobid=2020040800435&gene=scaffold1%7Csize1187143_573) | 3 | GH36 (http://www.cazy.org/GH36.html)(8-717) | GH36 (http://www.cazy.org/GH36.html) | GH36 (http://www.cazy.org/GH36.html) | N |
| scaffold1\|size1187143_582 (domain.php?jobid=2020040800435&gene=scaffold1%7Csize1187143_582) | 3 | GH13_36 (http://www.cazy.org/GH13_36.html)(75-417) | GH13_36 (http://www.cazy.org/GH13_36.html) | GH13 (http://www.cazy.org/GH13.html) | Y (1-32) |
| scaffold1\|size1187143_596 | 1 | N | GT13 (http://www.cazy.org/GT13.html) | N | N |
| scaffold1\|size1187143_611 (domain.php?jobid=2020040800435&gene=scaffold1%7Csize1187143_611) | 3 | GH13 (http://www.cazy.org/GH13.html)(27-364) | GH13 (http://www.cazy.org/GH13.html) | GH13 (http://www.cazy.org/GH13.html) | N |
| scaffold1\|size1187143_632 (domain.php?jobid=2020040800435&gene=scaffold1%7Csize1187143_632) | 1 | GT2_Glycos_transf_2 (http://www.cazy.org/GT2_Glycos_transf_2.html)(16-145) | N | N | N |
| scaffold1\|size1187143_634 (domain.php?jobid=2020040800435&gene=scaffold1%7Csize1187143_634) | 2 | GT2_Glycos_transf_2 (http://www.cazy.org/GT2_Glycos_transf_2.html)(5-127) | N | GT2 (http://www.cazy.org/GT2.html) | N |
| scaffold1\|size1187143_635 (domain.php?jobid=2020040800435&gene=scaffold1%7Csize1187143_635) | 1 | GT4 (http://www.cazy.org/GT4.html)(194-301) | N | N | N |
| scaffold1\|size1187143_636 (domain.php?jobid=2020040800435&gene=scaffold1%7Csize1187143_636) | 1 | GT4 (http://www.cazy.org/GT4.html)(199-342) | N | N | N |
| scaffold1\|size1187143_746 | 1 | N | CE12 (http://www.cazy.org/CE12.html) | N | N |
| scaffold1\|size1187143_777 (domain.php?jobid=2020040800435&gene=scaffold1%7Csize1187143_777) | 3 | CE9 (http://www.cazy.org/CE9.html)(4-373) | CE9 (http://www.cazy.org/CE9.html) | CE9 (http://www.cazy.org/CE9.html) | N |
| scaffold1\|size1187143_784 (domain.php?jobid=2020040800435&gene=scaffold1%7Csize1187143_784) | 3 | GH94 (http://www.cazy.org/GH94.html)(2-797) | GH94 (http://www.cazy.org/GH94.html) | GH94 (http://www.cazy.org/GH94.html) | N |

| Gene ID | # of Tools | HMMER | DIAMOND | Hotpep | Signal Peptide |
|---|---|---|---|---|---|
| scaffold1\|size1187143_821 (domain.php?jobid=2020040800435&gene=scaffold1%7Csize1187143_821) | 2 | N | GT2 (http://www.cazy.org/GT2.html) | GT2 (http://www.cazy.org/GT2.html) | N |
| scaffold1\|size1187143_827 | 1 | N | GT2 (http://www.cazy.org/GT2.html) | N | N |
| scaffold1\|size1187143_851 (domain.php?jobid=2020040800435&gene=scaffold1%7Csize1187143_851) | 3 | GH77 (http://www.cazy.org/GH77.html)(16-503) | GH77 (http://www.cazy.org/GH77.html) | GH77 (http://www.cazy.org/GH77.html) | N |
| scaffold1\|size1187143_877 (domain.php?jobid=2020040800435&gene=scaffold1%7Csize1187143_877) | 3 | GH94 (http://www.cazy.org/GH94.html)(2-808) | GH94 (http://www.cazy.org/GH94.html) | GH94 (http://www.cazy.org/GH94.html) | N |
| scaffold1\|size1187143_887 (domain.php?jobid=2020040800435&gene=scaffold1%7Csize1187143_887) | 3 | GT51 (http://www.cazy.org/GT51.html)(81-264) | GT51 (http://www.cazy.org/GT51.html) | GT51 (http://www.cazy.org/GT51.html) | N |
| scaffold1\|size1187143_907 (domain.php?jobid=2020040800435&gene=scaffold1%7Csize1187143_907) | 2 | GH78 (http://www.cazy.org/GH78.html)(439-965) | GH78 (http://www.cazy.org/GH78.html) | N | N |
| scaffold1\|size1187143_922 (domain.php?jobid=2020040800435&gene=scaffold1%7Csize1187143_922) | 3 | GH105 (http://www.cazy.org/GH105.html)(36-378) | GH105 (http://www.cazy.org/GH105.html) | GH105 (http://www.cazy.org/GH105.html) | N |
| scaffold1\|size1187143_95 (domain.php?jobid=2020040800435&gene=scaffold1%7Csize1187143_95) | 3 | CBM4 (http://www.cazy.org/CBM4.html)(70-196)+GH9 (http://www.cazy.org/GH9.html)(325-757) | GH9 (http://www.cazy.org/GH9.html) | GH9 (http://www.cazy.org/GH9.html)+CBM4 (http://www.cazy.org/CBM4.html) | Y (1-23) |
| scaffold1\|size1187143_983 (domain.php?jobid=2020040800435&gene=scaffold1%7Csize1187143_983) | 1 | CE4 (http://www.cazy.org/CE4.html)(173-358) | N | N | N |
| scaffold1\|size1187143_984 (domain.php?jobid=2020040800435&gene=scaffold1%7Csize1187143_984) | 1 | GT4 (http://www.cazy.org/GT4.html)(367-522) | N | N | N |
| scaffold1\|size1187143_985 (domain.php?jobid=2020040800435&gene=scaffold1%7Csize1187143_985) | 2 | GT2_Glycos_transf_2 (http://www.cazy.org/GT2_Glycos_transf_2.html)(6-139) | N | GT2 (http://www.cazy.org/GT2.html) | N |
| scaffold1\|size1187143_993 | 1 | N | GT2 (http://www.cazy.org/GT2.html) | N | N |
| scaffold2\|size606019_138 (domain.php?jobid=2020040800435&gene=scaffold2%7Csize606019_138) | 2 | GH13_18 (http://www.cazy.org/GH13_18.html)(240-452) | GH13_18 (http://www.cazy.org/GH13_18.html) | N | N |
| scaffold2\|size606019_18 (domain.php?jobid=2020040800435&gene=scaffold2%7Csize606019_18) | 3 | GT2_Glycos_transf_2 (http://www.cazy.org/GT2_Glycos_transf_2.html)(5-121) | GT2 (http://www.cazy.org/GT2.html) | GT2 (http://www.cazy.org/GT2.html) | N |
| scaffold2\|size606019_181 (domain.php?jobid=2020040800435&gene=scaffold2%7Csize606019_181) | 3 | GT35 (http://www.cazy.org/GT35.html)(94-768) | GT35 (http://www.cazy.org/GT35.html) | GT35 (http://www.cazy.org/GT35.html) | N |
| scaffold2\|size606019_209 (domain.php?jobid=2020040800435&gene=scaffold2%7Csize606019_209) | 3 | GH35 (http://www.cazy.org/GH35.html)(16-363) | GH35 (http://www.cazy.org/GH35.html) | GH35 (http://www.cazy.org/GH35.html) | N |
| scaffold2\|size606019_284 | 1 | N | GH6 (http://www.cazy.org/GH6.html) | N | N |
| scaffold2\|size606019_393 (domain.php?jobid=2020040800435&gene=scaffold2%7Csize606019_393) | 3 | GH94 (http://www.cazy.org/GH94.html)(116-886) | GH94 (http://www.cazy.org/GH94.html) | GH94 (http://www.cazy.org/GH94.html) | N |

| Gene ID | # of Tools | HMMER | DIAMOND | Hotpep | Signal Peptide |
|---|---|---|---|---|---|
| scaffold2\|size606019_56 | 1 | N | GH6 (http://www.cazy.org/GH6.html) | N | N |
| scaffold2\|size606019_57 (domain.php?jobid=2020040800435&gene=scaffold2%7Csize606019_57) | 2 | GH28 (http://www.cazy.org/GH28.html)(125-489) | GH28 (http://www.cazy.org/GH28.html) | N | N |
| scaffold2\|size606019_58 (domain.php?jobid=2020040800435&gene=scaffold2%7Csize606019_58) | 3 | CE12 (http://www.cazy.org/CE12.html)(173-368) | CE12 (http://www.cazy.org/CE12.html) | CE12 (http://www.cazy.org/CE12.html) | N |
| scaffold2\|size606019_59 (domain.php?jobid=2020040800435&gene=scaffold2%7Csize606019_59) | 3 | GH112 (http://www.cazy.org/GH112.html)(10-720) | GH112 (http://www.cazy.org/GH112.html) | GH112 (http://www.cazy.org/GH112.html) | N |
| scaffold2\|size606019_60 (domain.php?jobid=2020040800435&gene=scaffold2%7Csize606019_60) | 3 | GH105 (http://www.cazy.org/GH105.html)(27-336) | GH105 (http://www.cazy.org/GH105.html) | GH105 (http://www.cazy.org/GH105.html) | N |
| scaffold2\|size606019_8 (domain.php?jobid=2020040800435&gene=scaffold2%7Csize606019_8) | 3 | GT2_Glycos_transf_2 (http://www.cazy.org/GT2_Glycos_transf_2.html)(5-111) | GT2 (http://www.cazy.org/GT2.html) | GT2 (http://www.cazy.org/GT2.html) | N |
| scaffold3\|size332484_130 (domain.php?jobid=2020040800435&gene=scaffold3%7Csize332484_130) | 3 | GH10 (http://www.cazy.org/GH10.html)(65-345) | GH10 (http://www.cazy.org/GH10.html) | GH10 (http://www.cazy.org/GH10.html) | N |
| scaffold3\|size332484_136 (domain.php?jobid=2020040800435&gene=scaffold3%7Csize332484_136) | 3 | GH36 (http://www.cazy.org/GH36.html)(12-707) | GH36 (http://www.cazy.org/GH36.html) | GH36 (http://www.cazy.org/GH36.html) | N |
| scaffold3\|size332484_140 (domain.php?jobid=2020040800435&gene=scaffold3%7Csize332484_140) | 3 | GH36 (http://www.cazy.org/GH36.html)(64-622) | GH36 (http://www.cazy.org/GH36.html) | GH36 (http://www.cazy.org/GH36.html) | N |
| scaffold3\|size332484_177 (domain.php?jobid=2020040800435&gene=scaffold3%7Csize332484_177) | 3 | GT2_Glycos_transf_2 (http://www.cazy.org/GT2_Glycos_transf_2.html)(5-171) | GT2 (http://www.cazy.org/GT2.html) | GT2 (http://www.cazy.org/GT2.html) | N |
| scaffold3\|size332484_178 (domain.php?jobid=2020040800435&gene=scaffold3%7Csize332484_178) | 2 | GT2_Glycos_transf_2 (http://www.cazy.org/GT2_Glycos_transf_2.html)(3-143) | GT2 (http://www.cazy.org/GT2.html) | N | N |
| scaffold3\|size332484_180 | 1 | N | GT0 (http://www.cazy.org/GT0.html) | N | N |
| scaffold3\|size332484_181 | 1 | N | GT2 (http://www.cazy.org/GT2.html) | N | N |
| scaffold3\|size332484_182 (domain.php?jobid=2020040800435&gene=scaffold3%7Csize332484_182) | 2 | GT4 (http://www.cazy.org/GT4.html)(226-390) | GT4 (http://www.cazy.org/GT4.html) | N | N |
| scaffold3\|size332484_183 (domain.php?jobid=2020040800435&gene=scaffold3%7Csize332484_183) | 2 | GT4 (http://www.cazy.org/GT4.html)(179-329) | GT4 (http://www.cazy.org/GT4.html) | N | N |
| scaffold3\|size332484_185 (domain.php?jobid=2020040800435&gene=scaffold3%7Csize332484_185) | 3 | GT2_Glycos_transf_2 (http://www.cazy.org/GT2_Glycos_transf_2.html)(9-173) | GT2 (http://www.cazy.org/GT2.html) | CE4 (http://www.cazy.org/CE4.html)+GT2 (http://www.cazy.org/GT2.html) | N |
| scaffold3\|size332484_19 (domain.php?jobid=2020040800435&gene=scaffold3%7Csize332484_19) | 3 | PL33_1 (http://www.cazy.org/PL33_1.html)(403-566) | PL33_1 (http://www.cazy.org/PL33_1.html) | PL33 (http://www.cazy.org/PL33.html) | N |
| scaffold3\|size332484_191 (domain.php?jobid=2020040800435&gene=scaffold3%7Csize332484_191) | 2 | GT2_Glycos_transf_2 (http://www.cazy.org/GT2_Glycos_transf_2.html)(4-132) | GT2 (http://www.cazy.org/GT2.html) | N | N |
| scaffold3\|size332484_20 (domain.php?jobid=2020040800435&gene=scaffold3%7Csize332484_20) | 3 | GH88 (http://www.cazy.org/GH88.html)(49-369) | GH88 (http://www.cazy.org/GH88.html) | GH88 (http://www.cazy.org/GH88.html) | N |

| Gene ID | # of Tools | HMMER | DIAMOND | Hotpep | Signal Peptide |
|---|---|---|---|---|---|
| scaffold3\|size332484_200 (domain.php?jobid=2020040800435&gene=scaffold3%7Csize332484_200) | 2 | GT2_Glycos_transf_2 (http://www.cazy.org/GT2_Glycos_transf_2.html)(7-140) | GT2 (http://www.cazy.org/GT2.html) | N | N |
| scaffold3\|size332484_208 (domain.php?jobid=2020040800435&gene=scaffold3%7Csize332484_208) | 2 | GT4 (http://www.cazy.org/GT4.html)(192-337) | GT4 (http://www.cazy.org/GT4.html) | N | N |
| scaffold3\|size332484_21 (domain.php?jobid=2020040800435&gene=scaffold3%7Csize332484_21) | 3 | GH154 (http://www.cazy.org/GH154.html)(18-384) | GH154 (http://www.cazy.org/GH154.html) | GH154 (http://www.cazy.org/GH154.html) | N |
| scaffold3\|size332484_212 (domain.php?jobid=2020040800435&gene=scaffold3%7Csize332484_212) | 2 | GT4 (http://www.cazy.org/GT4.html)(223-342) | GT4 (http://www.cazy.org/GT4.html) | N | N |
| scaffold3\|size332484_213 (domain.php?jobid=2020040800435&gene=scaffold3%7Csize332484_213) | 2 | GT4 (http://www.cazy.org/GT4.html)(189-304) | GT4 (http://www.cazy.org/GT4.html) | N | N |
| scaffold3\|size332484_214 (domain.php?jobid=2020040800435&gene=scaffold3%7Csize332484_214) | 2 | GT4 (http://www.cazy.org/GT4.html)(183-335) | GT4 (http://www.cazy.org/GT4.html) | N | N |
| scaffold3\|size332484_218 (domain.php?jobid=2020040800435&gene=scaffold3%7Csize332484_218) | 2 | GT4 (http://www.cazy.org/GT4.html)(221-375) | GT4 (http://www.cazy.org/GT4.html) | N | N |
| scaffold3\|size332484_220 (domain.php?jobid=2020040800435&gene=scaffold3%7Csize332484_220) | 3 | GT11 (http://www.cazy.org/GT11.html)(1-282) | GT11 (http://www.cazy.org/GT11.html) | GT11 (http://www.cazy.org/GT11.html) | N |
| scaffold3\|size332484_222 (domain.php?jobid=2020040800435&gene=scaffold3%7Csize332484_222) | 2 | GT11 (http://www.cazy.org/GT11.html)(10-259) | GT11 (http://www.cazy.org/GT11.html) | N | N |
| scaffold3\|size332484_224 (domain.php?jobid=2020040800435&gene=scaffold3%7Csize332484_224) | 2 | GT17 (http://www.cazy.org/GT17.html)(2-279) | GT17 (http://www.cazy.org/GT17.html) | N | N |
| scaffold3\|size332484_226 (domain.php?jobid=2020040800435&gene=scaffold3%7Csize332484_226) | 1 | GT2_Glycos_transf_2 (http://www.cazy.org/GT2_Glycos_transf_2.html)(4-115) | N | N | N |
| scaffold3\|size332484_229 (domain.php?jobid=2020040800435&gene=scaffold3%7Csize332484_229) | 2 | GT2_Glycos_transf_2 (http://www.cazy.org/GT2_Glycos_transf_2.html)(10-123) | GT2 (http://www.cazy.org/GT2.html) | N | N |
| scaffold3\|size332484_231 (domain.php?jobid=2020040800435&gene=scaffold3%7Csize332484_231) | 3 | GT11 (http://www.cazy.org/GT11.html)(1-296) | GT11 (http://www.cazy.org/GT11.html) | GT11 (http://www.cazy.org/GT11.html) | N |
| scaffold3\|size332484_233 (domain.php?jobid=2020040800435&gene=scaffold3%7Csize332484_233) | 1 | GH105 (http://www.cazy.org/GH105.html)(45-368) | N | N | N |
| scaffold3\|size332484_241 (domain.php?jobid=2020040800435&gene=scaffold3%7Csize332484_241) | 3 | GH30_1 (http://www.cazy.org/GH30_1.html)(31-443) | GH30_1 (http://www.cazy.org/GH30_1.html) | GH30 (http://www.cazy.org/GH30.html) | N |
| scaffold3\|size332484_25 | 1 | N | AA1 (http://www.cazy.org/AA1.html) | N | N |
| scaffold3\|size332484_54 | 1 | N | GT4 (http://www.cazy.org/GT4.html) | N | N |
| scaffold3\|size332484_61 (domain.php?jobid=2020040800435&gene=scaffold3%7Csize332484_61) | 3 | CE10 (http://www.cazy.org/CE10.html)(82-231)+GH43_35 (http://www.cazy.org/GH43_35.html)(467-768) | GH43_35 (http://www.cazy.org/GH43_35.html) | GH43 (http://www.cazy.org/GH43.html) | N |

| Gene ID | # of Tools | HMMER | DIAMOND | Hotpep | Signal Peptide |
|---|---|---|---|---|---|
| scaffold3\|size332484_63 | 1 | N | GH11 (http://www.cazy.org/GH11.html) | N | N |
| scaffold3\|size332484_65 (domain.php?jobid=2020040800435&gene=scaffold3%7Csize332484_65) | 3 | GH67 (http://www.cazy.org/GH67.html)(8-658) | GH67 (http://www.cazy.org/GH67.html) | GH67 (http://www.cazy.org/GH67.html) | N |
| scaffold3\|size332484_67 (domain.php?jobid=2020040800435&gene=scaffold3%7Csize332484_67) | 3 | GH115 (http://www.cazy.org/GH115.html)(8-645) | GH115 (http://www.cazy.org/GH115.html) | GH115 (http://www.cazy.org/GH115.html) | N |
| scaffold3\|size332484_71 (domain.php?jobid=2020040800435&gene=scaffold3%7Csize332484_71) | 3 | GH3 (http://www.cazy.org/GH3.html)(33-272) | GH3 (http://www.cazy.org/GH3.html) | GH3 (http://www.cazy.org/GH3.html) | N |
| scaffold3\|size332484_76 (domain.php?jobid=2020040800435&gene=scaffold3%7Csize332484_76) | 2 | GH25 (http://www.cazy.org/GH25.html)(43-212) | GH25 (http://www.cazy.org/GH25.html) | N | Y (1-30) |
| scaffold3\|size332484_90 | 1 | N | CBM48 (http://www.cazy.org/CBM48.html)+GH13_9 (http://www.cazy.org/GH13_9.html) | N | N |
| scaffold4\|size280552_126 (domain.php?jobid=2020040800435&gene=scaffold4%7Csize280552_126) | 3 | GH5_2 (http://www.cazy.org/GH5_2.html)(55-291)+GT2_Glyco_tranf_2_3 (http://www.cazy.org/GT2_Glyco_tranf_2_3.html)(499-738) | GH5_2 (http://www.cazy.org/GH5_2.html)+GT2 (http://www.cazy.org/GT2.html) | GH5 (http://www.cazy.org/GH5.html)+GT2 (http://www.cazy.org/GT2.html) | Y (1-30) |
| scaffold4\|size280552_165 (domain.php?jobid=2020040800435&gene=scaffold4%7Csize280552_165) | 1 | CE1 (http://www.cazy.org/CE1.html)(59-208) | N | N | N |
| scaffold4\|size280552_175 (domain.php?jobid=2020040800435&gene=scaffold4%7Csize280552_175) | 3 | GH51 (http://www.cazy.org/GH51.html)(3-500) | GH51 (http://www.cazy.org/GH51.html) | GH51 (http://www.cazy.org/GH51.html) | N |
| scaffold4\|size280552_206 (domain.php?jobid=2020040800435&gene=scaffold4%7Csize280552_206) | 1 | GT32 (http://www.cazy.org/GT32.html)(138-214) | N | N | N |
| scaffold4\|size280552_227 (domain.php?jobid=2020040800435&gene=scaffold4%7Csize280552_227) | 1 | CE10 (http://www.cazy.org/CE10.html)(46-284) | N | N | N |
| scaffold4\|size280552_232 (domain.php?jobid=2020040800435&gene=scaffold4%7Csize280552_232) | 1 | CE10 (http://www.cazy.org/CE10.html)(25-275) | N | N | N |
| scaffold4\|size280552_265 (domain.php?jobid=2020040800435&gene=scaffold4%7Csize280552_265) | 3 | GH3 (http://www.cazy.org/GH3.html)(685-918) | GH3 (http://www.cazy.org/GH3.html) | GH3 (http://www.cazy.org/GH3.html) | N |
| scaffold4\|size280552_266 (domain.php?jobid=2020040800435&gene=scaffold4%7Csize280552_266) | 3 | GH3 (http://www.cazy.org/GH3.html)(31-249) | GH3 (http://www.cazy.org/GH3.html) | GH3 (http://www.cazy.org/GH3.html) | N |
| scaffold4\|size280552_272 (domain.php?jobid=2020040800435&gene=scaffold4%7Csize280552_272) | 3 | GH53 (http://www.cazy.org/GH53.html)(58-421) | CBM61 (http://www.cazy.org/CBM61.html)+GH53 (http://www.cazy.org/GH53.html) | GH53 (http://www.cazy.org/GH53.html) | N |
| scaffold4\|size280552_35 (domain.php?jobid=2020040800435&gene=scaffold4%7Csize280552_35) | 1 | GH25 (http://www.cazy.org/GH25.html)(49-219) | N | N | N |
| scaffold4\|size280552_74 (domain.php?jobid=2020040800435&gene=scaffold4%7Csize280552_74) | 2 | GH16 (http://www.cazy.org/GH16.html)(71-318)+CBM4 (http://www.cazy.org/CBM4.html)(349-486)+CBM4 (http://www.cazy.org/CBM4.html)(658-793) | CBM4 (http://www.cazy.org/CBM4.html)+GH16 (http://www.cazy.org/GH16.html) | N | Y (1-44) |
| scaffold5\|size256508_1 (domain.php?jobid=2020040800435&gene=scaffold5%7Csize256508_1) | 1 | GH18 (http://www.cazy.org/GH18.html)(72-363) | N | N | N |

| Gene ID | # of Tools | HMMER | DIAMOND | Hotpep | Signal Peptide |
|---|---|---|---|---|---|
| scaffold5\|size256508_181 (domain.php?jobid=2020040800435&gene=scaffold5%7Csize256508_181) | 1 | CE10 (http://www.cazy.org/CE10.html)(76-300) | N | N | N |
| scaffold5\|size256508_183 (domain.php?jobid=2020040800435&gene=scaffold5%7Csize256508_183) | 2 | GT1 (http://www.cazy.org/GT1.html)(193-409) | GT1 (http://www.cazy.org/GT1.html) | N | N |
| scaffold5\|size256508_220 | 1 | N | GH13_30 (http://www.cazy.org/GH13_30.html) | N | N |
| scaffold5\|size256508_227 (domain.php?jobid=2020040800435&gene=scaffold5%7Csize256508_227) | 3 | GH2 (http://www.cazy.org/GH2.html)(41-536) | GH2 (http://www.cazy.org/GH2.html) | GH2 (http://www.cazy.org/GH2.html) | N |
| scaffold5\|size256508_238 (domain.php?jobid=2020040800435&gene=scaffold5%7Csize256508_238) | 2 | GH18 (http://www.cazy.org/GH18.html)(226-555) | GH18 (http://www.cazy.org/GH18.html) | N | N |
| scaffold5\|size256508_240 | 1 | N | GT2 (http://www.cazy.org/GT2.html) | N | N |
| scaffold5\|size256508_3 (domain.php?jobid=2020040800435&gene=scaffold5%7Csize256508_3) | 2 | GT2_Glycos_transf_2 (http://www.cazy.org/GT2_Glycos_transf_2.html)(242-418) | GT2 (http://www.cazy.org/GT2.html) | N | N |
| scaffold5\|size256508_4 (domain.php?jobid=2020040800435&gene=scaffold5%7Csize256508_4) | 1 | CE4 (http://www.cazy.org/CE4.html)(816-952) | N | N | Y (1-24) |
| scaffold6\|size244034_105 (domain.php?jobid=2020040800435&gene=scaffold6%7Csize244034_105) | 1 | CE10 (http://www.cazy.org/CE10.html)(17-254) | N | N | N |
| scaffold6\|size244034_128 (domain.php?jobid=2020040800435&gene=scaffold6%7Csize244034_128) | 3 | GH13_39 (http://www.cazy.org/GH13_39.html)(173-553) | CBM34 (http://www.cazy.org/CBM34.html)+GH13 (http://www.cazy.org/GH13.html) | GH13 (http://www.cazy.org/GH13.html)+CBM34 (http://www.cazy.org/CBM34.html) | N |
| scaffold6\|size244034_159 (domain.php?jobid=2020040800435&gene=scaffold6%7Csize244034_159) | 3 | GT35 (http://www.cazy.org/GT35.html)(97-818) | GT35 (http://www.cazy.org/GT35.html) | GT35 (http://www.cazy.org/GT35.html) | N |
| scaffold6\|size244034_171 (domain.php?jobid=2020040800435&gene=scaffold6%7Csize244034_171) | 3 | GH43_26 (http://www.cazy.org/GH43_26.html)(11-313) | GH43_26 (http://www.cazy.org/GH43_26.html) | GH43 (http://www.cazy.org/GH43.html) | N |
| scaffold6\|size244034_172 (domain.php?jobid=2020040800435&gene=scaffold6%7Csize244034_172) | 2 | GH43_4 (http://www.cazy.org/GH43_4.html)(54-402) | GH43_4 (http://www.cazy.org/GH43_4.html) | N | Y (1-27) |
| scaffold6\|size244034_173 (domain.php?jobid=2020040800435&gene=scaffold6%7Csize244034_173) | 3 | GH51 (http://www.cazy.org/GH51.html)(2-500) | GH51 (http://www.cazy.org/GH51.html) | GH51 (http://www.cazy.org/GH51.html) | N |
| scaffold6\|size244034_215 (domain.php?jobid=2020040800435&gene=scaffold6%7Csize244034_215) | 2 | GH9 (http://www.cazy.org/GH9.html)(143-487) | GH9 (http://www.cazy.org/GH9.html) | N | Y (1-26) |
| scaffold6\|size244034_39 (domain.php?jobid=2020040800435&gene=scaffold6%7Csize244034_39) | 3 | CBM48 (http://www.cazy.org/CBM48.html)(26-110)+GH13_9 (http://www.cazy.org/GH13_9.html)(182-482) | CBM48 (http://www.cazy.org/CBM48.html)+GH13_9 (http://www.cazy.org/GH13_9.html) | GH13 (http://www.cazy.org/GH13.html)+CBM48 (http://www.cazy.org/CBM48.html) | N |
| scaffold6\|size244034_66 (domain.php?jobid=2020040800435&gene=scaffold6%7Csize244034_66) | 3 | GT28 (http://www.cazy.org/GT28.html)(193-342) | GT28 (http://www.cazy.org/GT28.html) | GT28 (http://www.cazy.org/GT28.html) | N |
| scaffold7\|size202071_104 (domain.php?jobid=2020040800435&gene=scaffold7%7Csize202071_104) | 1 | GT2_Glycos_transf_2 (http://www.cazy.org/GT2_Glycos_transf_2.html)(6-180) | N | N | N |

| Gene ID | # of Tools | HMMER | DIAMOND | Hotpep | Signal Peptide |
|---|---|---|---|---|---|
| scaffold7\|size202071_112 (domain.php?jobid=2020040800435&gene=scaffold7%7Csize202071_112) | 3 | GT2_Glycos_transf_2 (http://www.cazy.org/GT2_Glycos_transf_2.html)(6-160) | GT2 (http://www.cazy.org/GT2.html) | GT2 (http://www.cazy.org/GT2.html) | N |
| scaffold7\|size202071_131 (domain.php?jobid=2020040800435&gene=scaffold7%7Csize202071_131) | 2 | GH25 (http://www.cazy.org/GH25.html)(1003-1183) | GH25 (http://www.cazy.org/GH25.html) | N | N |
| scaffold7\|size202071_132 (domain.php?jobid=2020040800435&gene=scaffold7%7Csize202071_132) | 1 | GT2_Glycos_transf_2 (http://www.cazy.org/GT2_Glycos_transf_2.html)(11-135) | N | N | N |
| scaffold7\|size202071_140 (domain.php?jobid=2020040800435&gene=scaffold7%7Csize202071_140) | 3 | GT2_Glycos_transf_2 (http://www.cazy.org/GT2_Glycos_transf_2.html)(260-446) | GT2 (http://www.cazy.org/GT2.html) | GT2 (http://www.cazy.org/GT2.html) | N |
| scaffold7\|size202071_141 (domain.php?jobid=2020040800435&gene=scaffold7%7Csize202071_141) | 3 | GT2_Glycos_transf_2 (http://www.cazy.org/GT2_Glycos_transf_2.html)(120-288)+GT2_Glycos_transf_2 (http://www.cazy.org/GT2_Glycos_transf_2.html)(390-584) | GT2 (http://www.cazy.org/GT2.html) | GT2 (http://www.cazy.org/GT2.html) | N |
| scaffold7\|size202071_148 (domain.php?jobid=2020040800435&gene=scaffold7%7Csize202071_148) | 2 | GT2_Glycos_transf_2 (http://www.cazy.org/GT2_Glycos_transf_2.html)(68-235) | GT2 (http://www.cazy.org/GT2.html) | N | N |
| scaffold7\|size202071_149 (domain.php?jobid=2020040800435&gene=scaffold7%7Csize202071_149) | 3 | GT2_Glycos_transf_2 (http://www.cazy.org/GT2_Glycos_transf_2.html)(5-193) | GT2 (http://www.cazy.org/GT2.html) | GT2 (http://www.cazy.org/GT2.html) | N |
| scaffold7\|size202071_15 (domain.php?jobid=2020040800435&gene=scaffold7%7Csize202071_15) | 3 | GH28 (http://www.cazy.org/GH28.html)(106-459) | GH28 (http://www.cazy.org/GH28.html) | GH28 (http://www.cazy.org/GH28.html) | N |
| scaffold7\|size202071_151 (domain.php?jobid=2020040800435&gene=scaffold7%7Csize202071_151) | 1 | GT2_Glycos_transf_2 (http://www.cazy.org/GT2_Glycos_transf_2.html)(6-168) | N | N | N |
| scaffold7\|size202071_154 (domain.php?jobid=2020040800435&gene=scaffold7%7Csize202071_154) | 2 | GT2_Glycos_transf_2 (http://www.cazy.org/GT2_Glycos_transf_2.html)(5-179) | GT2 (http://www.cazy.org/GT2.html) | N | N |
| scaffold7\|size202071_16 (domain.php?jobid=2020040800435&gene=scaffold7%7Csize202071_16) | 3 | CE8 (http://www.cazy.org/CE8.html)(4-312) | CE8 (http://www.cazy.org/CE8.html) | CE8 (http://www.cazy.org/CE8.html) | N |
| scaffold7\|size202071_168 (domain.php?jobid=2020040800435&gene=scaffold7%7Csize202071_168) | 1 | GT4 (http://www.cazy.org/GT4.html)(175-322) | N | N | N |
| scaffold7\|size202071_46 (domain.php?jobid=2020040800435&gene=scaffold7%7Csize202071_46) | 1 | GT83 (http://www.cazy.org/GT83.html)(6-380) | N | N | N |
| scaffold7\|size202071_47 (domain.php?jobid=2020040800435&gene=scaffold7%7Csize202071_47) | 1 | GT2_Glycos_transf_2 (http://www.cazy.org/GT2_Glycos_transf_2.html)(4-162) | N | N | N |
| scaffold7\|size202071_49 (domain.php?jobid=2020040800435&gene=scaffold7%7Csize202071_49) | 1 | GT83 (http://www.cazy.org/GT83.html)(13-379) | N | N | N |
| scaffold7\|size202071_53 (domain.php?jobid=2020040800435&gene=scaffold7%7Csize202071_53) | 3 | GT2_Glycos_transf_2 (http://www.cazy.org/GT2_Glycos_transf_2.html)(5-162) | GT2 (http://www.cazy.org/GT2.html) | GT2 (http://www.cazy.org/GT2.html) | N |
| scaffold7\|size202071_56 (domain.php?jobid=2020040800435&gene=scaffold7%7Csize202071_56) | 2 | GT26 (http://www.cazy.org/GT26.html)(72-243) | N | GT26 (http://www.cazy.org/GT26.html) | N |
| scaffold7\|size202071_64 | 1 | N | GT4 (http://www.cazy.org/GT4.html) | N | N |

| Gene ID | # of Tools | HMMER | DIAMOND | Hotpep | Signal Peptide |
|---|---|---|---|---|---|
| scaffold7\|size202071_65 (domain.php?jobid=2020040800435&gene=scaffold7%7Csize202071_65) | 1 | GH109 (http://www.cazy.org/GH109.html)(2-111) | N | N | N |
| scaffold7\|size202071_71 (domain.php?jobid=2020040800435&gene=scaffold7%7Csize202071_71) | 1 | GT4 (http://www.cazy.org/GT4.html)(226-370) | N | N | N |
| scaffold8\|size198603_106 (domain.php?jobid=2020040800435&gene=scaffold8%7Csize198603_106) | 2 | GH25 (http://www.cazy.org/GH25.html)(166-348) | GH25 (http://www.cazy.org/GH25.html) | N | N |
| scaffold8\|size198603_143 (domain.php?jobid=2020040800435&gene=scaffold8%7Csize198603_143) | 2 | GH13_11 (http://www.cazy.org/GH13_11.html)(291-482) | GH13_11 (http://www.cazy.org/GH13_11.html) | N | N |
| scaffold8\|size198603_15 (domain.php?jobid=2020040800435&gene=scaffold8%7Csize198603_15) | 3 | GT5 (http://www.cazy.org/GT5.html)(3-476) | GT5 (http://www.cazy.org/GT5.html) | GT5 (http://www.cazy.org/GT5.html) | N |
| scaffold8\|size198603_61 (domain.php?jobid=2020040800435&gene=scaffold8%7Csize198603_61) | 3 | GH16 (http://www.cazy.org/GH16.html)(244-470)+CBM4 (http://www.cazy.org/CBM4.html)(502-639)+CBM4 (http://www.cazy.org/CBM4.html)(813-947) | CBM4 (http://www.cazy.org/CBM4.html)+GH16 (http://www.cazy.org/GH16.html) | GH16 (http://www.cazy.org/GH16.html) | Y (1-34) |
| scaffold8\|size198603_79 | 1 | N | GT4 (http://www.cazy.org/GT4.html) | N | N |
| scaffold8\|size198603_93 (domain.php?jobid=2020040800435&gene=scaffold8%7Csize198603_93) | 3 | GT2_Glycos_transf_2 (http://www.cazy.org/GT2_Glycos_transf_2.html)(7-172) | GT2 (http://www.cazy.org/GT2.html) | GT2 (http://www.cazy.org/GT2.html) | N |
| scaffold9\|size67573_5 (domain.php?jobid=2020040800435&gene=scaffold9%7Csize67573_5) | 3 | GH13_11 (http://www.cazy.org/GH13_11.html)(206-560) | CBM48 (http://www.cazy.org/CBM48.html)+GH13_11 (http://www.cazy.org/GH13_11.html) | GH13 (http://www.cazy.org/GH13.html) | N |

Showing 1 to 161 of 161 entries

First Previous 1 Next Last

Copyright 2017 © YIN LAB (http://bcb.unl.edu), UNL (http://www.unl.edu). All rights reserved. Designed by Tanner Yohe and Le Huang. Maintained by Yanbin Yin. (http://bcb.unl.edu/dbCAN2/about.php)

### dbCAN meta server: automated CAZyme annotation

(index.php)

Home (index.php) | Annotate (blast.php) | Download (download/) | Help (help.php) | About us (about.php)

**Cite us: NAR/gky418 (https://doi.org/10.1093/nar/gky418) and gks479 (https://doi.org/10.1093/nar/gks479)**

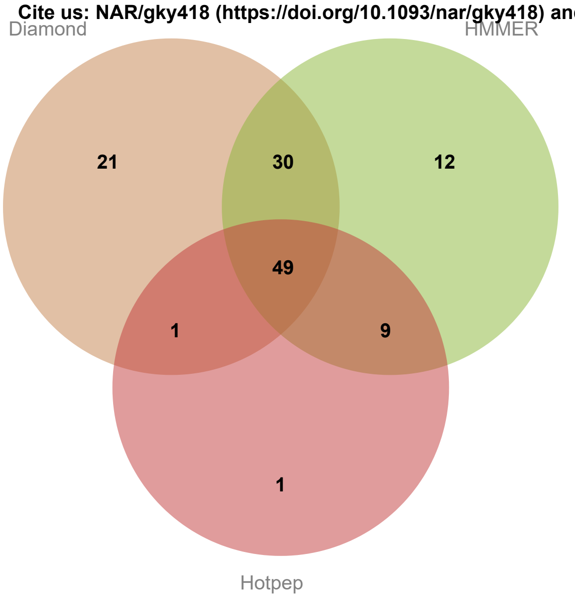


Result of job: 2020040823022

Overview | HMMER | DIAMOND | Hotpep | CGC-Finder

**Download SignalP output (data/blast/2020040823022/signalp.out) Download Prodigal predictions (data/blast/2020040823022/uniInput) Download this table (keep those with # of Tools >=2 will give you best result; and use dbCAN domain assignment is recommended) (data/blast/2020040823022/overview.txt) ? (help.php#over)**

**Show** All ▼ **entries** **Search:**

| Gene ID | # of Tools | HMMER | DIAMOND | Hotpep | Signal Peptide |
|---|---|---|---|---|---|
| scaffold10\|size136614_15 (domain.php?jobid=2020040823022&gene=scaffold10%7Csize136614_15) | 2 | GT2_Glycos_transf_2 (http://www.cazy.org/GT2_Glycos_transf_2.html)(5-178) | GT2 (http://www.cazy.org/GT2.html) | N | N |
| scaffold10\|size136614_18 (domain.php?jobid=2020040823022&gene=scaffold10%7Csize136614_18) | 2 | GT2_Glycos_transf_2 (http://www.cazy.org/GT2_Glycos_transf_2.html)(5-176) | GT2 (http://www.cazy.org/GT2.html) | N | N |
| scaffold10\|size136614_19 (domain.php?jobid=2020040823022&gene=scaffold10%7Csize136614_19) | 2 | GT2_Glycos_transf_2 (http://www.cazy.org/GT2_Glycos_transf_2.html)(68-238) | GT2 (http://www.cazy.org/GT2.html) | N | N |
| scaffold10\|size136614_21 (domain.php?jobid=2020040823022&gene=scaffold10%7Csize136614_21) | 3 | GT2_Glycos_transf_2 (http://www.cazy.org/GT2_Glycos_transf_2.html)(122-290)+GT2_Glycos_transf_2 (http://www.cazy.org/GT2_Glycos_transf_2.html)(391-586) | GT2 (http://www.cazy.org/GT2.html) | GT2 (http://www.cazy.org/GT2.html) | N |
| scaffold10\|size136614_22 (domain.php?jobid=2020040823022&gene=scaffold10%7Csize136614_22) | 3 | GT2_Glycos_transf_2 (http://www.cazy.org/GT2_Glycos_transf_2.html)(260-445) | GT2 (http://www.cazy.org/GT2.html) | GT2 (http://www.cazy.org/GT2.html) | N |
| scaffold10\|size136614_31 (domain.php?jobid=2020040823022&gene=scaffold10%7Csize136614_31) | 2 | GH25 (http://www.cazy.org/GH25.html)(1149-1329) | GH25 (http://www.cazy.org/GH25.html) | N | N |
| scaffold10\|size136614_46 (domain.php?jobid=2020040823022&gene=scaffold10%7Csize136614_46) | 2 | GT2_Glycos_transf_2 (http://www.cazy.org/GT2_Glycos_transf_2.html)(5-147) | GT2 (http://www.cazy.org/GT2.html) | N | N |
| scaffold10\|size136614_50 | 1 | N | GT0 (http://www.cazy.org/GT0.html) | N | N |

| Gene ID | # of Tools | HMMER | DIAMOND | Hotpep | Signal Peptide |
|---|---|---|---|---|---|
| scaffold10\|size136614_52 | 1 | N | GT4 (http://www.cazy.org/GT4.html) | N | N |
| scaffold10\|size136614_65 | 1 | N | GT4 (http://www.cazy.org/GT4.html) | N | N |
| scaffold10\|size136614_66 | 1 | N | GT4 (http://www.cazy.org/GT4.html) | N | N |
| scaffold10\|size136614_80 (domain.php?jobid=2020040823022&gene=scaffold10%7Csize136614_80) | 3 | GT2_Glycos_transf_2 (http://www.cazy.org/GT2_Glycos_transf_2.html)(103-270) | GT2 (http://www.cazy.org/GT2.html)+GT4 (http://www.cazy.org/GT4.html) | GT2 (http://www.cazy.org/GT2.html)+GT4 (http://www.cazy.org/GT4.html) | N |
| scaffold10\|size136614_85 (domain.php?jobid=2020040823022&gene=scaffold10%7Csize136614_85) | 2 | GT4 (http://www.cazy.org/GT4.html)(189-331) | N | GT4 (http://www.cazy.org/GT4.html) | N |
| scaffold10\|size136614_86 (domain.php?jobid=2020040823022&gene=scaffold10%7Csize136614_86) | 2 | N | GT4 (http://www.cazy.org/GT4.html) | GT4 (http://www.cazy.org/GT4.html) | N |
| scaffold10\|size136614_94 (domain.php?jobid=2020040823022&gene=scaffold10%7Csize136614_94) | 1 | GT4 (http://www.cazy.org/GT4.html)(298-443) | N | N | N |
| scaffold10\|size136614_95 (domain.php?jobid=2020040823022&gene=scaffold10%7Csize136614_95) | 1 | GT2_Glycos_transf_2 (http://www.cazy.org/GT2_Glycos_transf_2.html)(5-122) | N | N | N |
| scaffold11\|size83008_30 (domain.php?jobid=2020040823022&gene=scaffold11%7Csize83008_30) | 2 | CE4 (http://www.cazy.org/CE4.html)(3-111) | N | CE4 (http://www.cazy.org/CE4.html) | N |
| scaffold11\|size83008_33 (domain.php?jobid=2020040823022&gene=scaffold11%7Csize83008_33) | 3 | GH8 (http://www.cazy.org/GH8.html)(52-376) | GH8 (http://www.cazy.org/GH8.html) | GH8 (http://www.cazy.org/GH8.html) | N |
| scaffold11\|size83008_78 (domain.php?jobid=2020040823022&gene=scaffold11%7Csize83008_78) | 2 | CE4 (http://www.cazy.org/CE4.html)(3-111) | N | CE4 (http://www.cazy.org/CE4.html) | N |
| scaffold11\|size83008_81 (domain.php?jobid=2020040823022&gene=scaffold11%7Csize83008_81) | 3 | GH8 (http://www.cazy.org/GH8.html)(52-376) | GH8 (http://www.cazy.org/GH8.html) | GH8 (http://www.cazy.org/GH8.html) | N |
| scaffold12\|size68416_13 (domain.php?jobid=2020040823022&gene=scaffold12%7Csize68416_13) | 2 | GT2_Glycos_transf_2 (http://www.cazy.org/GT2_Glycos_transf_2.html)(5-132) | GT2 (http://www.cazy.org/GT2.html) | N | N |
| scaffold12\|size68416_19 (domain.php?jobid=2020040823022&gene=scaffold12%7Csize68416_19) | 3 | GT26 (http://www.cazy.org/GT26.html)(286-457) | GT26 (http://www.cazy.org/GT26.html) | GT26 (http://www.cazy.org/GT26.html) | N |
| scaffold12\|size68416_3 (domain.php?jobid=2020040823022&gene=scaffold12%7Csize68416_3) | 3 | GT2_Glycos_transf_2 (http://www.cazy.org/GT2_Glycos_transf_2.html)(6-160) | GT2 (http://www.cazy.org/GT2.html) | GT2 (http://www.cazy.org/GT2.html) | N |
| scaffold12\|size68416_60 (domain.php?jobid=2020040823022&gene=scaffold12%7Csize68416_60) | 3 | CE10 (http://www.cazy.org/CE10.html)(82-236)+GH43_35 (http://www.cazy.org/GH43_35.html)(475-777) | GH43_35 (http://www.cazy.org/GH43_35.html) | GH43 (http://www.cazy.org/GH43.html) | N |
| scaffold13\|size34950_16 | 1 | N | AA1 (http://www.cazy.org/AA1.html) | N | N |
| scaffold14\|size25486_19 | 1 | N | GH28 (http://www.cazy.org/GH28.html) | N | N |
| scaffold14\|size25486_5 (domain.php?jobid=2020040823022&gene=scaffold14%7Csize25486_5) | 1 | GT4 (http://www.cazy.org/GT4.html)(200-357) | N | N | N |
| scaffold14\|size25486_8 (domain.php?jobid=2020040823022&gene=scaffold14%7Csize25486_8) | 1 | GT2_Glycos_transf_2 (http://www.cazy.org/GT2_Glycos_transf_2.html)(7-108) | N | N | N |

| Gene ID | # of Tools | HMMER | DIAMOND | Hotpep | Signal Peptide |
|---|---|---|---|---|---|
| scaffold1\|size697872_163 (domain.php?jobid=2020040823022&gene=scaffold1%7Csize697872_163) | 2 | GH3 (http://www.cazy.org/GH3.html)(193-385) | GH3 (http://www.cazy.org/GH3.html) | N | N |
| scaffold1\|size697872_164 | 1 | N | GH13_11 (http://www.cazy.org/GH13_11.html) | N | N |
| scaffold1\|size697872_167 (domain.php?jobid=2020040823022&gene=scaffold1%7Csize697872_167) | 3 | GT2_Glycos_transf_2 (http://www.cazy.org/GT2_Glycos_transf_2.html)(5-108) | GT2 (http://www.cazy.org/GT2.html) | GT2 (http://www.cazy.org/GT2.html) | N |
| scaffold1\|size697872_208 (domain.php?jobid=2020040823022&gene=scaffold1%7Csize697872_208) | 3 | GH13_20 (http://www.cazy.org/GH13_20.html)(34-320) | GH13 (http://www.cazy.org/GH13.html) | GH13 (http://www.cazy.org/GH13.html) | N |
| scaffold1\|size697872_397 (domain.php?jobid=2020040823022&gene=scaffold1%7Csize697872_397) | 3 | GH36 (http://www.cazy.org/GH36.html)(63-575) | GH36 (http://www.cazy.org/GH36.html) | GH36 (http://www.cazy.org/GH36.html) | N |
| scaffold1\|size697872_450 (domain.php?jobid=2020040823022&gene=scaffold1%7Csize697872_450) | 3 | GH3 (http://www.cazy.org/GH3.html)(31-249) | GH3 (http://www.cazy.org/GH3.html) | GH3 (http://www.cazy.org/GH3.html) | N |
| scaffold1\|size697872_451 (domain.php?jobid=2020040823022&gene=scaffold1%7Csize697872_451) | 3 | GH3 (http://www.cazy.org/GH3.html)(695-889) | GH3 (http://www.cazy.org/GH3.html) | GH3 (http://www.cazy.org/GH3.html) | N |
| scaffold1\|size697872_526 (domain.php?jobid=2020040823022&gene=scaffold1%7Csize697872_526) | 3 | GH154 (http://www.cazy.org/GH154.html)(19-383) | GH154 (http://www.cazy.org/GH154.html) | GH154 (http://www.cazy.org/GH154.html) | N |
| scaffold1\|size697872_527 (domain.php?jobid=2020040823022&gene=scaffold1%7Csize697872_527) | 3 | GH88 (http://www.cazy.org/GH88.html)(50-372) | GH88 (http://www.cazy.org/GH88.html) | GH88 (http://www.cazy.org/GH88.html) | N |
| scaffold1\|size697872_528 (domain.php?jobid=2020040823022&gene=scaffold1%7Csize697872_528) | 3 | PL33_1 (http://www.cazy.org/PL33_1.html)(405-565) | PL33_1 (http://www.cazy.org/PL33_1.html) | PL0 (http://www.cazy.org/PL0.html)+PL33 (http://www.cazy.org/PL33.html) | N |
| scaffold1\|size697872_538 | 1 | N | GT13 (http://www.cazy.org/GT13.html) | N | N |
| scaffold1\|size697872_594 (domain.php?jobid=2020040823022&gene=scaffold1%7Csize697872_594) | 3 | GH5_37 (http://www.cazy.org/GH5_37.html)(10-324) | GH5_37 (http://www.cazy.org/GH5_37.html) | GH5 (http://www.cazy.org/GH5.html) | N |
| scaffold1\|size697872_595 (domain.php?jobid=2020040823022&gene=scaffold1%7Csize697872_595) | 3 | GH13_36 (http://www.cazy.org/GH13_36.html)(27-362) | GH13 (http://www.cazy.org/GH13.html) | GH13 (http://www.cazy.org/GH13.html) | N |
| scaffold1\|size697872_596 (domain.php?jobid=2020040823022&gene=scaffold1%7Csize697872_596) | 2 | CE2 (http://www.cazy.org/CE2.html)(130-344) | N | CE2 (http://www.cazy.org/CE2.html) | N |
| scaffold1\|size697872_600 (domain.php?jobid=2020040823022&gene=scaffold1%7Csize697872_600) | 2 | CE9 (http://www.cazy.org/CE9.html)(3-372) | N | CE9 (http://www.cazy.org/CE9.html) | N |
| scaffold1\|size697872_603 (domain.php?jobid=2020040823022&gene=scaffold1%7Csize697872_603) | 3 | GT35 (http://www.cazy.org/GT35.html)(94-768) | GT35 (http://www.cazy.org/GT35.html) | GT35 (http://www.cazy.org/GT35.html) | N |
| scaffold1\|size697872_628 (domain.php?jobid=2020040823022&gene=scaffold1%7Csize697872_628) | 3 | GH94 (http://www.cazy.org/GH94.html)(2-792) | GH94 (http://www.cazy.org/GH94.html) | GH94 (http://www.cazy.org/GH94.html) | N |
| scaffold1\|size697872_93 (domain.php?jobid=2020040823022&gene=scaffold1%7Csize697872_93) | 3 | CE4 (http://www.cazy.org/CE4.html)(112-215) | CE4 (http://www.cazy.org/CE4.html) | CE4 (http://www.cazy.org/CE4.html) | N |

| Gene ID | # of Tools | HMMER | DIAMOND | Hotpep | Signal Peptide |
|---|---|---|---|---|---|
| scaffold2\|size580642_141 (domain.php?jobid=2020040823022&gene=scaffold2%7Csize580642_141) | 2 | GH25 (http://www.cazy.org/GH25.html)(166-348) | GH25 (http://www.cazy.org/GH25.html) | N | N |
| scaffold2\|size580642_213 (domain.php?jobid=2020040823022&gene=scaffold2%7Csize580642_213) | 2 | GH13_11 (http://www.cazy.org/GH13_11.html)(292-483) | GH13_11 (http://www.cazy.org/GH13_11.html) | N | N |
| scaffold2\|size580642_271 (domain.php?jobid=2020040823022&gene=scaffold2%7Csize580642_271) | 2 | GH120 (http://www.cazy.org/GH120.html)(299-388) | GH120 (http://www.cazy.org/GH120.html) | N | N |
| scaffold2\|size580642_349 (domain.php?jobid=2020040823022&gene=scaffold2%7Csize580642_349) | 2 | CBM48 (http://www.cazy.org/CBM48.html)(127-213)+GH13_9 (http://www.cazy.org/GH13_9.html)(279-572) | CBM48 (http://www.cazy.org/CBM48.html)+GH13_9 (http://www.cazy.org/GH13_9.html) | N | N |
| scaffold2\|size580642_378 (domain.php?jobid=2020040823022&gene=scaffold2%7Csize580642_378) | 3 | GT2_Glycos_transf_2 (http://www.cazy.org/GT2_Glycos_transf_2.html)(7-173) | GT2 (http://www.cazy.org/GT2.html) | GT2 (http://www.cazy.org/GT2.html) | N |
| scaffold2\|size580642_392 | 1 | N | GT4 (http://www.cazy.org/GT4.html) | N | N |
| scaffold2\|size580642_416 (domain.php?jobid=2020040823022&gene=scaffold2%7Csize580642_416) | 1 | GH109 (http://www.cazy.org/GH109.html)(2-116) | N | N | N |
| scaffold2\|size580642_496 (domain.php?jobid=2020040823022&gene=scaffold2%7Csize580642_496) | 2 | CE1 (http://www.cazy.org/CE1.html)(129-361) | N | CE1 (http://www.cazy.org/CE1.html) | N |
| scaffold2\|size580642_497 (domain.php?jobid=2020040823022&gene=scaffold2%7Csize580642_497) | 2 | GH13_18 (http://www.cazy.org/GH13_18.html)(40-400) | GH13_18 (http://www.cazy.org/GH13_18.html) | N | N |
| scaffold2\|size580642_499 (domain.php?jobid=2020040823022&gene=scaffold2%7Csize580642_499) | 2 | CE1 (http://www.cazy.org/CE1.html)(39-262) | N | CE1 (http://www.cazy.org/CE1.html) | N |
| scaffold2\|size580642_511 (domain.php?jobid=2020040823022&gene=scaffold2%7Csize580642_511) | 3 | GT5 (http://www.cazy.org/GT5.html)(3-476) | GT5 (http://www.cazy.org/GT5.html) | GT5 (http://www.cazy.org/GT5.html) | N |
| scaffold2\|size580642_6 (domain.php?jobid=2020040823022&gene=scaffold2%7Csize580642_6) | 3 | GT2_Glycos_transf_2 (http://www.cazy.org/GT2_Glycos_transf_2.html)(3-128) | GT2 (http://www.cazy.org/GT2.html) | GT2 (http://www.cazy.org/GT2.html) | N |
| scaffold2\|size580642_69 (domain.php?jobid=2020040823022&gene=scaffold2%7Csize580642_69) | 3 | GH3 (http://www.cazy.org/GH3.html)(109-341) | GH3 (http://www.cazy.org/GH3.html) | GH3 (http://www.cazy.org/GH3.html) | Y (1-31) |
| scaffold2\|size580642_70 (domain.php?jobid=2020040823022&gene=scaffold2%7Csize580642_70) | 2 | GT4 (http://www.cazy.org/GT4.html)(181-332) | GT4 (http://www.cazy.org/GT4.html) | N | N |
| scaffold3\|size380913_147 (domain.php?jobid=2020040823022&gene=scaffold3%7Csize380913_147) | 3 | GH10 (http://www.cazy.org/GH10.html)(74-345) | GH10 (http://www.cazy.org/GH10.html) | GH10 (http://www.cazy.org/GH10.html) | N |
| scaffold3\|size380913_212 (domain.php?jobid=2020040823022&gene=scaffold3%7Csize380913_212) | 1 | CBM2 (http://www.cazy.org/CBM2.html)(479-561)+CBM2 (http://www.cazy.org/CBM2.html)(586-673) | N | N | Y (1-35) |
| scaffold3\|size380913_241 (domain.php?jobid=2020040823022&gene=scaffold3%7Csize380913_241) | 3 | GT2_Glycos_transf_2 (http://www.cazy.org/GT2_Glycos_transf_2.html)(5-166) | GT2 (http://www.cazy.org/GT2.html) | GT2 (http://www.cazy.org/GT2.html) | N |
| scaffold3\|size380913_242 (domain.php?jobid=2020040823022&gene=scaffold3%7Csize380913_242) | 2 | GT2_Glycos_transf_2 (http://www.cazy.org/GT2_Glycos_transf_2.html)(3-124) | GT2 (http://www.cazy.org/GT2.html) | N | N |

| Gene ID | # of Tools | HMMER | DIAMOND | Hotpep | Signal Peptide |
|---|---|---|---|---|---|
| scaffold3\|size380913_247 | 1 | N | GT2 (http://www.cazy.org/GT2.html) | N | N |
| scaffold3\|size380913_248 (domain.php?jobid=2020040823022&gene=scaffold3%7Csize380913_248) | 2 | GT4 (http://www.cazy.org/GT4.html)(224-390) | GT4 (http://www.cazy.org/GT4.html) | N | N |
| scaffold3\|size380913_249 (domain.php?jobid=2020040823022&gene=scaffold3%7Csize380913_249) | 2 | GT4 (http://www.cazy.org/GT4.html)(197-341) | GT4 (http://www.cazy.org/GT4.html) | N | N |
| scaffold3\|size380913_251 (domain.php?jobid=2020040823022&gene=scaffold3%7Csize380913_251) | 2 | GT2_Glycos_transf_2 (http://www.cazy.org/GT2_Glycos_transf_2.html)(9-176) | GT2 (http://www.cazy.org/GT2.html) | N | N |
| scaffold3\|size380913_256 | 1 | N | GT0 (http://www.cazy.org/GT0.html) | N | N |
| scaffold3\|size380913_257 (domain.php?jobid=2020040823022&gene=scaffold3%7Csize380913_257) | 2 | GT2_Glycos_transf_2 (http://www.cazy.org/GT2_Glycos_transf_2.html)(5-163) | GT2 (http://www.cazy.org/GT2.html) | N | N |
| scaffold3\|size380913_258 | 1 | N | GT100 (http://www.cazy.org/GT100.html) | N | N |
| scaffold3\|size380913_266 (domain.php?jobid=2020040823022&gene=scaffold3%7Csize380913_266) | 2 | GT2_Glycos_transf_2 (http://www.cazy.org/GT2_Glycos_transf_2.html)(9-142) | GT2 (http://www.cazy.org/GT2.html) | N | N |
| scaffold3\|size380913_277 (domain.php?jobid=2020040823022&gene=scaffold3%7Csize380913_277) | 2 | GH25 (http://www.cazy.org/GH25.html)(68-241) | N | GH25 (http://www.cazy.org/GH25.html) | N |
| scaffold3\|size380913_286 (domain.php?jobid=2020040823022&gene=scaffold3%7Csize380913_286) | 2 | GT4 (http://www.cazy.org/GT4.html)(192-337) | GT4 (http://www.cazy.org/GT4.html) | N | N |
| scaffold3\|size380913_290 (domain.php?jobid=2020040823022&gene=scaffold3%7Csize380913_290) | 2 | GT4 (http://www.cazy.org/GT4.html)(225-369) | GT4 (http://www.cazy.org/GT4.html) | N | N |
| scaffold3\|size380913_291 (domain.php?jobid=2020040823022&gene=scaffold3%7Csize380913_291) | 2 | GT4 (http://www.cazy.org/GT4.html)(188-304) | GT4 (http://www.cazy.org/GT4.html) | N | N |
| scaffold3\|size380913_293 (domain.php?jobid=2020040823022&gene=scaffold3%7Csize380913_293) | 2 | GT4 (http://www.cazy.org/GT4.html)(180-336) | GT4 (http://www.cazy.org/GT4.html) | N | N |
| scaffold3\|size380913_298 (domain.php?jobid=2020040823022&gene=scaffold3%7Csize380913_298) | 2 | GT4 (http://www.cazy.org/GT4.html)(224-376) | GT4 (http://www.cazy.org/GT4.html) | N | N |
| scaffold3\|size380913_303 (domain.php?jobid=2020040823022&gene=scaffold3%7Csize380913_303) | 3 | GT11 (http://www.cazy.org/GT11.html)(1-280) | GT11 (http://www.cazy.org/GT11.html) | GT11 (http://www.cazy.org/GT11.html) | N |
| scaffold3\|size380913_305 (domain.php?jobid=2020040823022&gene=scaffold3%7Csize380913_305) | 2 | GT11 (http://www.cazy.org/GT11.html)(10-256) | GT11 (http://www.cazy.org/GT11.html) | N | N |
| scaffold3\|size380913_307 (domain.php?jobid=2020040823022&gene=scaffold3%7Csize380913_307) | 1 | GT2_Glycos_transf_2 (http://www.cazy.org/GT2_Glycos_transf_2.html)(4-106) | N | N | N |
| scaffold3\|size380913_308 (domain.php?jobid=2020040823022&gene=scaffold3%7Csize380913_308) | 2 | GT17 (http://www.cazy.org/GT17.html)(2-282) | GT17 (http://www.cazy.org/GT17.html) | N | N |

| Gene ID | # of Tools | HMMER | DIAMOND | Hotpep | Signal Peptide |
|---|---|---|---|---|---|
| scaffold3\|size380913_311 (domain.php?jobid=2020040823022&gene=scaffold3%7Csize380913_311) | 2 | GT2_Glycos_transf_2 (http://www.cazy.org/GT2_Glycos_transf_2.html)(10-119) | GT2 (http://www.cazy.org/GT2.html) | N | N |
| scaffold3\|size380913_313 (domain.php?jobid=2020040823022&gene=scaffold3%7Csize380913_313) | 3 | GT11 (http://www.cazy.org/GT11.html)(1-300) | GT11 (http://www.cazy.org/GT11.html) | GT11 (http://www.cazy.org/GT11.html) | N |
| scaffold3\|size380913_315 (domain.php?jobid=2020040823022&gene=scaffold3%7Csize380913_315) | 1 | GH105 (http://www.cazy.org/GH105.html)(118-430) | N | N | N |
| scaffold3\|size380913_48 | 1 | N | CBM50 (http://www.cazy.org/CBM50.html) | N | Y (1-23) |
| scaffold3\|size380913_59 | 1 | N | GT4 (http://www.cazy.org/GT4.html) | N | N |
| scaffold3\|size380913_63 | 1 | N | GH11 (http://www.cazy.org/GH11.html) | N | N |
| scaffold3\|size380913_65 (domain.php?jobid=2020040823022&gene=scaffold3%7Csize380913_65) | 3 | GH67 (http://www.cazy.org/GH67.html)(55-645) | GH67 (http://www.cazy.org/GH67.html) | GH67 (http://www.cazy.org/GH67.html) | N |
| scaffold3\|size380913_66 (domain.php?jobid=2020040823022&gene=scaffold3%7Csize380913_66) | 3 | GH3 (http://www.cazy.org/GH3.html)(32-272) | GH3 (http://www.cazy.org/GH3.html) | GH3 (http://www.cazy.org/GH3.html) | N |
| scaffold3\|size380913_69 (domain.php?jobid=2020040823022&gene=scaffold3%7Csize380913_69) | 2 | GH25 (http://www.cazy.org/GH25.html)(42-211) | GH25 (http://www.cazy.org/GH25.html) | N | Y (1-34) |
| scaffold3\|size380913_79 (domain.php?jobid=2020040823022&gene=scaffold3%7Csize380913_79) | 3 | GH2 (http://www.cazy.org/GH2.html)(43-525) | GH2 (http://www.cazy.org/GH2.html) | GH2 (http://www.cazy.org/GH2.html) | N |
| scaffold3\|size380913_90 | 1 | N | GH13_30 (http://www.cazy.org/GH13_30.html) | N | N |
| scaffold4\|size330350_12 | 1 | N | GT2 (http://www.cazy.org/GT2.html) | N | N |
| scaffold4\|size330350_14 (domain.php?jobid=2020040823022&gene=scaffold4%7Csize330350_14) | 2 | GH18 (http://www.cazy.org/GH18.html)(255-554) | GH18 (http://www.cazy.org/GH18.html) | N | N |
| scaffold4\|size330350_187 (domain.php?jobid=2020040823022&gene=scaffold4%7Csize330350_187) | 1 | N | N | GT2 (http://www.cazy.org/GT2.html) | N |
| scaffold4\|size330350_22 (domain.php?jobid=2020040823022&gene=scaffold4%7Csize330350_22) | 3 | GH2 (http://www.cazy.org/GH2.html)(30-901) | GH2 (http://www.cazy.org/GH2.html) | GH2 (http://www.cazy.org/GH2.html) | N |
| scaffold4\|size330350_227 (domain.php?jobid=2020040823022&gene=scaffold4%7Csize330350_227) | 3 | GT51 (http://www.cazy.org/GT51.html)(86-265) | GT51 (http://www.cazy.org/GT51.html) | GT51 (http://www.cazy.org/GT51.html) | N |
| scaffold4\|size330350_237 (domain.php?jobid=2020040823022&gene=scaffold4%7Csize330350_237) | 3 | GH94 (http://www.cazy.org/GH94.html)(2-813) | GH94 (http://www.cazy.org/GH94.html) | GH94 (http://www.cazy.org/GH94.html) | N |
| scaffold4\|size330350_257 (domain.php?jobid=2020040823022&gene=scaffold4%7Csize330350_257) | 3 | GH77 (http://www.cazy.org/GH77.html)(20-504) | GH77 (http://www.cazy.org/GH77.html) | GH77 (http://www.cazy.org/GH77.html) | N |
| scaffold4\|size330350_28 | 1 | N | CBM48 (http://www.cazy.org/CBM48.html)+GH13_9 (http://www.cazy.org/GH13_9.html) | N | N |
| scaffold4\|size330350_281 | 1 | N | GH6 (http://www.cazy.org/GH6.html) | N | N |

| Gene ID | # of Tools | HMMER | DIAMOND | Hotpep | Signal Peptide |
|---|---|---|---|---|---|
| scaffold5\|size287620_137 (domain.php?jobid=2020040823022&gene=scaffold5%7Csize287620_137) | 2 | PL9_2 (http://www.cazy.org/PL9_2.html)(53-402) | N | PL9 (http://www.cazy.org/PL9.html) | Y (1-51) |
| scaffold5\|size287620_240 | 1 | N | GT2 (http://www.cazy.org/GT2.html) | N | N |
| scaffold5\|size287620_245 (domain.php?jobid=2020040823022&gene=scaffold5%7Csize287620_245) | 3 | GT2_Glycos_transf_2 (http://www.cazy.org/GT2_Glycos_transf_2.html)(5-144) | GT2 (http://www.cazy.org/GT2.html) | GT2 (http://www.cazy.org/GT2.html) | N |
| scaffold5\|size287620_272 (domain.php?jobid=2020040823022&gene=scaffold5%7Csize287620_272) | 3 | GT2_Glycos_transf_2 (http://www.cazy.org/GT2_Glycos_transf_2.html)(9-154) | GT2 (http://www.cazy.org/GT2.html) | GT2 (http://www.cazy.org/GT2.html) | N |
| scaffold5\|size287620_4 | 1 | N | GH6 (http://www.cazy.org/GH6.html) | N | N |
| scaffold5\|size287620_79 (domain.php?jobid=2020040823022&gene=scaffold5%7Csize287620_79) | 3 | GH35 (http://www.cazy.org/GH35.html)(17-363) | GH35 (http://www.cazy.org/GH35.html) | GH35 (http://www.cazy.org/GH35.html) | N |
| scaffold6\|size182103_10 (domain.php?jobid=2020040823022&gene=scaffold6%7Csize182103_10) | 3 | GT35 (http://www.cazy.org/GT35.html)(97-818) | GT35 (http://www.cazy.org/GT35.html) | GT35 (http://www.cazy.org/GT35.html) | N |
| scaffold6\|size182103_127 (domain.php?jobid=2020040823022&gene=scaffold6%7Csize182103_127) | 3 | CBM48 (http://www.cazy.org/CBM48.html)(26-110)+GH13_9 (http://www.cazy.org/GH13_9.html)(182-482) | CBM48 (http://www.cazy.org/CBM48.html)+GH13_9 (http://www.cazy.org/GH13_9.html) | GH13 (http://www.cazy.org/GH13.html)+CBM48 (http://www.cazy.org/CBM48.html) | N |
| scaffold6\|size182103_34 (domain.php?jobid=2020040823022&gene=scaffold6%7Csize182103_34) | 3 | GH13_39 (http://www.cazy.org/GH13_39.html)(173-553) | CBM34 (http://www.cazy.org/CBM34.html)+GH13 (http://www.cazy.org/GH13.html) | GH13 (http://www.cazy.org/GH13.html)+CBM34 (http://www.cazy.org/CBM34.html) | N |
| scaffold6\|size182103_96 (domain.php?jobid=2020040823022&gene=scaffold6%7Csize182103_96) | 3 | GH51 (http://www.cazy.org/GH51.html)(8-504) | GH51 (http://www.cazy.org/GH51.html) | GH51 (http://www.cazy.org/GH51.html) | N |
| scaffold6\|size182103_99 (domain.php?jobid=2020040823022&gene=scaffold6%7Csize182103_99) | 3 | GT28 (http://www.cazy.org/GT28.html)(193-341) | GT28 (http://www.cazy.org/GT28.html) | GT28 (http://www.cazy.org/GT28.html) | N |
| scaffold7\|size159935_108 (domain.php?jobid=2020040823022&gene=scaffold7%7Csize159935_108) | 3 | GH3 (http://www.cazy.org/GH3.html)(558-766) | GH3 (http://www.cazy.org/GH3.html) | GH3 (http://www.cazy.org/GH3.html) | N |
| scaffold7\|size159935_62 (domain.php?jobid=2020040823022&gene=scaffold7%7Csize159935_62) | 3 | GH35 (http://www.cazy.org/GH35.html)(10-321) | GH35 (http://www.cazy.org/GH35.html) | GH35 (http://www.cazy.org/GH35.html) | N |
| scaffold7\|size159935_63 (domain.php?jobid=2020040823022&gene=scaffold7%7Csize159935_63) | 3 | GH2 (http://www.cazy.org/GH2.html)(6-560) | GH2 (http://www.cazy.org/GH2.html) | GH2 (http://www.cazy.org/GH2.html) | N |
| scaffold8\|size157267_145 (domain.php?jobid=2020040823022&gene=scaffold8%7Csize157267_145) | 1 | CE13 (http://www.cazy.org/CE13.html)(64-258) | N | N | N |
| scaffold8\|size157267_148 (domain.php?jobid=2020040823022&gene=scaffold8%7Csize157267_148) | 3 | GH120 (http://www.cazy.org/GH120.html)(297-387) | GH120 (http://www.cazy.org/GH120.html) | GH120 (http://www.cazy.org/GH120.html) | N |
| scaffold8\|size157267_6 (domain.php?jobid=2020040823022&gene=scaffold8%7Csize157267_6) | 3 | GH13_11 (http://www.cazy.org/GH13_11.html)(208-561) | CBM48 (http://www.cazy.org/CBM48.html)+GH13_11 (http://www.cazy.org/GH13_11.html) | GH13 (http://www.cazy.org/GH13.html) | N |
| scaffold9\|size138208_1 (domain.php?jobid=2020040823022&gene=scaffold9%7Csize138208_1) | 1 | CBM2 (http://www.cazy.org/CBM2.html)(688-768) | N | N | N |
| scaffold9\|size138208_19 (domain.php?jobid=2020040823022&gene=scaffold9%7Csize138208_19) | 1 | CBM32 (http://www.cazy.org/CBM32.html)(14-113) | N | N | N |

| Gene ID | # of Tools | HMMER | DIAMOND | Hotpep | Signal Peptide |
|---|---|---|---|---|---|
| scaffold9\|size138208_2 (domain.php?jobid=2020040823022&gene=scaffold9%7Csize138208_2) | 1 | CBM2 (http://www.cazy.org/CBM2.html)(362-436) | N | N | Y (1-28) |
| scaffold9\|size138208_51 (domain.php?jobid=2020040823022&gene=scaffold9%7Csize138208_51) | 2 | GH9 (http://www.cazy.org/GH9.html)(152-491) | GH9 (http://www.cazy.org/GH9.html) | N | Y (1-32) |

Showing 1 to 123 of 123 entries

First Previous 1 Next Last

Copyright 2017 © YIN LAB (http://bcb.unl.edu), UNL (http://www.unl.edu). All rights reserved. Designed by Tanner Yohe and Le Huang. Maintained by Yanbin Yin. (http://bcb.unl.edu/dbCAN2/about.php)

dbCAN meta server: automated CAZyme annotation

(index.php)

Home (index.php) | Annotate (blast.php) | Download (download/) | Help (help.php) | About us (about.php)

**Cite us: NAR/gky418 (https://doi.org/10.1093/nar/gky418) and gks479 (https://doi.org/10.1093/nar/gks479)**

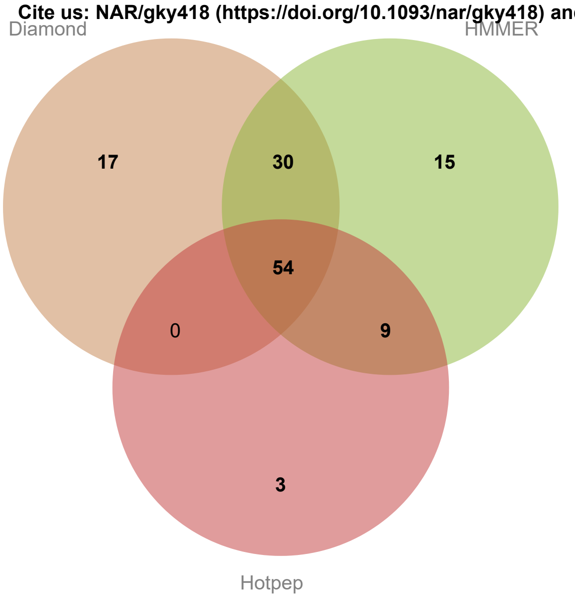


Result of job: 2020040810636

Overview HMMER DIAMOND Hotpep CGC-Finder

**Download SignalP output (data/blast/2020040810636/signalp.out) Download Prodigal predictions (data/blast/2020040810636/uniInput) Download this table (keep those with # of Tools >=2 will give you best result; and use dbCAN domain assignment is recommended) (data/blast/2020040810636/overview.txt) ? (help.php#over)**

**Show** All ▼ **entries** **Search:**

| Gene ID | # of Tools | HMMER | DIAMOND | Hotpep | Signal Peptide |
|---|---|---|---|---|---|
| scaffold10\|size141526_112 (domain.php?jobid=2020040810636&gene=scaffold10%7Csize141526_112) | 1 | CBM32 (http://www.cazy.org/CBM32.html)(12-113) | N | N | N |
| scaffold10\|size141526_131 (domain.php?jobid=2020040810636&gene=scaffold10%7Csize141526_131) | 1 | CBM2 (http://www.cazy.org/CBM2.html)(361-435) | N | N | Y (1-29) |
| scaffold10\|size141526_132 (domain.php?jobid=2020040810636&gene=scaffold10%7Csize141526_132) | 1 | CBM2 (http://www.cazy.org/CBM2.html)(593-673) | N | N | N |
| scaffold10\|size141526_36 (domain.php?jobid=2020040810636&gene=scaffold10%7Csize141526_36) | 1 | GT2_Glycos_transf_2 (http://www.cazy.org/GT2_Glycos_transf_2.html)(4-160) | N | N | N |
| scaffold10\|size141526_79 (domain.php?jobid=2020040810636&gene=scaffold10%7Csize141526_79) | 2 | GH9 (http://www.cazy.org/GH9.html)(154-488) | GH9 (http://www.cazy.org/GH9.html) | N | Y (1-32) |
| scaffold11\|size84195_26 (domain.php?jobid=2020040810636&gene=scaffold11%7Csize84195_26) | 3 | GH8 (http://www.cazy.org/GH8.html)(52-376) | GH8 (http://www.cazy.org/GH8.html) | GH8 (http://www.cazy.org/GH8.html) | N |
| scaffold11\|size84195_28 (domain.php?jobid=2020040810636&gene=scaffold11%7Csize84195_28) | 2 | CE4 (http://www.cazy.org/CE4.html)(3-112) | N | CE4 (http://www.cazy.org/CE4.html) | N |
| scaffold11\|size84195_74 (domain.php?jobid=2020040810636&gene=scaffold11%7Csize84195_74) | 3 | GH8 (http://www.cazy.org/GH8.html)(52-376) | GH8 (http://www.cazy.org/GH8.html) | GH8 (http://www.cazy.org/GH8.html) | N |

| Gene ID | # of Tools | HMMER | DIAMOND | Hotpep | Signal Peptide |
|---|---|---|---|---|---|
| scaffold11\|size84195_76 (domain.php?jobid=2020040810636&gene=scaffold11%7Csize84195_76) | 2 | CE4 (http://www.cazy.org/CE4.html)(3-112) | N | CE4 (http://www.cazy.org/CE4.html) | N |
| scaffold12\|size77156_6 (domain.php?jobid=2020040810636&gene=scaffold12%7Csize77156_6) | 3 | GH13_11 (http://www.cazy.org/GH13_11.html)(208-561) | CBM48 (http://www.cazy.org/CBM48.html)+GH13_11 (http://www.cazy.org/GH13_11.html) | GH13 (http://www.cazy.org/GH13.html) | N |
| scaffold13\|size65078_10 (domain.php?jobid=2020040810636&gene=scaffold13%7Csize65078_10) | 3 | CE10 (http://www.cazy.org/CE10.html)(82-236)+GH43_35 (http://www.cazy.org/GH43_35.html)(475-776) | GH43_35 (http://www.cazy.org/GH43_35.html) | GH43 (http://www.cazy.org/GH43.html) | N |
| scaffold13\|size65078_31 | 1 | N | GH6 (http://www.cazy.org/GH6.html) | N | N |
| scaffold13\|size65078_47 (domain.php?jobid=2020040810636&gene=scaffold13%7Csize65078_47) | 3 | GH77 (http://www.cazy.org/GH77.html)(20-505) | GH77 (http://www.cazy.org/GH77.html) | GH77 (http://www.cazy.org/GH77.html) | N |
| scaffold14\|size51804_13 (domain.php?jobid=2020040810636&gene=scaffold14%7Csize51804_13) | 2 | GT2_Glycos_transf_2 (http://www.cazy.org/GT2_Glycos_transf_2.html)(11-138) | GT2 (http://www.cazy.org/GT2.html) | N | N |
| scaffold14\|size51804_18 (domain.php?jobid=2020040810636&gene=scaffold14%7Csize51804_18) | 3 | GT26 (http://www.cazy.org/GT26.html)(286-457) | GT26 (http://www.cazy.org/GT26.html) | GT26 (http://www.cazy.org/GT26.html) | N |
| scaffold14\|size51804_3 (domain.php?jobid=2020040810636&gene=scaffold14%7Csize51804_3) | 3 | GT2_Glycos_transf_2 (http://www.cazy.org/GT2_Glycos_transf_2.html)(6-160) | GT2 (http://www.cazy.org/GT2.html) | GT2 (http://www.cazy.org/GT2.html) | N |
| scaffold1\|size609713_123 (domain.php?jobid=2020040810636&gene=scaffold1%7Csize609713_123) | 3 | GH13_20 (http://www.cazy.org/GH13_20.html)(34-320) | GH13 (http://www.cazy.org/GH13.html) | GH13 (http://www.cazy.org/GH13.html) | N |
| scaffold1\|size609713_18 (domain.php?jobid=2020040810636&gene=scaffold1%7Csize609713_18) | 3 | CE4 (http://www.cazy.org/CE4.html)(114-217) | CE4 (http://www.cazy.org/CE4.html) | CE4 (http://www.cazy.org/CE4.html) | N |
| scaffold1\|size609713_327 (domain.php?jobid=2020040810636&gene=scaffold1%7Csize609713_327) | 3 | GH3 (http://www.cazy.org/GH3.html)(31-249) | GH3 (http://www.cazy.org/GH3.html) | GH3 (http://www.cazy.org/GH3.html) | N |
| scaffold1\|size609713_328 (domain.php?jobid=2020040810636&gene=scaffold1%7Csize609713_328) | 3 | GH3 (http://www.cazy.org/GH3.html)(695-889) | GH3 (http://www.cazy.org/GH3.html) | GH3 (http://www.cazy.org/GH3.html) | N |
| scaffold1\|size609713_402 (domain.php?jobid=2020040810636&gene=scaffold1%7Csize609713_402) | 3 | GH154 (http://www.cazy.org/GH154.html)(19-383) | GH154 (http://www.cazy.org/GH154.html) | GH154 (http://www.cazy.org/GH154.html) | N |
| scaffold1\|size609713_403 (domain.php?jobid=2020040810636&gene=scaffold1%7Csize609713_403) | 3 | GH88 (http://www.cazy.org/GH88.html)(50-372) | GH88 (http://www.cazy.org/GH88.html) | GH88 (http://www.cazy.org/GH88.html) | N |
| scaffold1\|size609713_404 (domain.php?jobid=2020040810636&gene=scaffold1%7Csize609713_404) | 3 | PL33_1 (http://www.cazy.org/PL33_1.html)(403-571) | PL33_1 (http://www.cazy.org/PL33_1.html) | PL33 (http://www.cazy.org/PL33.html) | N |
| scaffold1\|size609713_414 | 1 | N | GT13 (http://www.cazy.org/GT13.html) | N | N |
| scaffold1\|size609713_463 (domain.php?jobid=2020040810636&gene=scaffold1%7Csize609713_463) | 3 | GH5_37 (http://www.cazy.org/GH5_37.html)(10-324) | GH5_37 (http://www.cazy.org/GH5_37.html) | GH5 (http://www.cazy.org/GH5.html) | N |
| scaffold1\|size609713_464 (domain.php?jobid=2020040810636&gene=scaffold1%7Csize609713_464) | 3 | GH13_36 (http://www.cazy.org/GH13_36.html)(27-362) | GH13 (http://www.cazy.org/GH13.html) | GH13 (http://www.cazy.org/GH13.html) | N |
| scaffold1\|size609713_465 (domain.php?jobid=2020040810636&gene=scaffold1%7Csize609713_465) | 2 | CE2 (http://www.cazy.org/CE2.html)(130-344) | N | CE2 (http://www.cazy.org/CE2.html) | N |

| Gene ID | # of Tools | HMMER | DIAMOND | Hotpep | Signal Peptide |
|---|---|---|---|---|---|
| scaffold1\|size609713_471 (domain.php?jobid=2020040810636&gene=scaffold1%7Csize609713_471) | 2 | CE9 (http://www.cazy.org/CE9.html)(3-372) | N | CE9 (http://www.cazy.org/CE9.html) | N |
| scaffold1\|size609713_474 (domain.php?jobid=2020040810636&gene=scaffold1%7Csize609713_474) | 3 | GT35 (http://www.cazy.org/GT35.html)(94-768) | GT35 (http://www.cazy.org/GT35.html) | GT35 (http://www.cazy.org/GT35.html) | N |
| scaffold1\|size609713_509 (domain.php?jobid=2020040810636&gene=scaffold1%7Csize609713_509) | 3 | GH94 (http://www.cazy.org/GH94.html)(2-792) | GH94 (http://www.cazy.org/GH94.html) | GH94 (http://www.cazy.org/GH94.html) | N |
| scaffold1\|size609713_79 (domain.php?jobid=2020040810636&gene=scaffold1%7Csize609713_79) | 2 | GH3 (http://www.cazy.org/GH3.html)(194-385) | GH3 (http://www.cazy.org/GH3.html) | N | N |
| scaffold1\|size609713_80 | 1 | N | GH13_11 (http://www.cazy.org/GH13_11.html) | N | N |
| scaffold1\|size609713_83 (domain.php?jobid=2020040810636&gene=scaffold1%7Csize609713_83) | 3 | GT2_Glycos_transf_2 (http://www.cazy.org/GT2_Glycos_transf_2.html)(5-108) | GT2 (http://www.cazy.org/GT2.html) | GT2 (http://www.cazy.org/GT2.html) | N |
| scaffold2\|size463030_149 | 1 | N | GH6 (http://www.cazy.org/GH6.html) | N | N |
| scaffold2\|size463030_229 (domain.php?jobid=2020040810636&gene=scaffold2%7Csize463030_229) | 3 | GH35 (http://www.cazy.org/GH35.html)(16-363) | GH35 (http://www.cazy.org/GH35.html) | GH35 (http://www.cazy.org/GH35.html) | N |
| scaffold2\|size463030_267 (domain.php?jobid=2020040810636&gene=scaffold2%7Csize463030_267) | 2 | GH25 (http://www.cazy.org/GH25.html)(47-220) | N | GH25 (http://www.cazy.org/GH25.html) | N |
| scaffold2\|size463030_281 (domain.php?jobid=2020040810636&gene=scaffold2%7Csize463030_281) | 2 | PL9_2 (http://www.cazy.org/PL9_2.html)(35-384) | N | PL9 (http://www.cazy.org/PL9.html) | Y (1-33) |
| scaffold2\|size463030_36 (domain.php?jobid=2020040810636&gene=scaffold2%7Csize463030_36) | 3 | GH3 (http://www.cazy.org/GH3.html)(559-766) | GH3 (http://www.cazy.org/GH3.html) | GH3 (http://www.cazy.org/GH3.html) | N |
| scaffold2\|size463030_379 | 1 | N | GT2 (http://www.cazy.org/GT2.html) | N | N |
| scaffold2\|size463030_384 (domain.php?jobid=2020040810636&gene=scaffold2%7Csize463030_384) | 3 | GT2_Glycos_transf_2 (http://www.cazy.org/GT2_Glycos_transf_2.html)(5-144) | GT2 (http://www.cazy.org/GT2.html) | GT2 (http://www.cazy.org/GT2.html) | N |
| scaffold2\|size463030_406 (domain.php?jobid=2020040810636&gene=scaffold2%7Csize463030_406) | 3 | GT2_Glycos_transf_2 (http://www.cazy.org/GT2_Glycos_transf_2.html)(7-121) | GT2 (http://www.cazy.org/GT2.html) | GT2 (http://www.cazy.org/GT2.html) | N |
| scaffold2\|size463030_414 (domain.php?jobid=2020040810636&gene=scaffold2%7Csize463030_414) | 1 | GT2_Glycos_transf_2 (http://www.cazy.org/GT2_Glycos_transf_2.html)(5-170) | N | N | N |
| scaffold2\|size463030_415 (domain.php?jobid=2020040810636&gene=scaffold2%7Csize463030_415) | 1 | GT8 (http://www.cazy.org/GT8.html)(3-262) | N | N | N |
| scaffold2\|size463030_416 (domain.php?jobid=2020040810636&gene=scaffold2%7Csize463030_416) | 3 | GH36 (http://www.cazy.org/GH36.html)(62-620) | GH36 (http://www.cazy.org/GH36.html) | GH36 (http://www.cazy.org/GH36.html) | N |
| scaffold2\|size463030_420 (domain.php?jobid=2020040810636&gene=scaffold2%7Csize463030_420) | 3 | GH36 (http://www.cazy.org/GH36.html)(12-707) | GH36 (http://www.cazy.org/GH36.html) | GH36 (http://www.cazy.org/GH36.html) | N |
| scaffold2\|size463030_430 | 1 | N | GH28 (http://www.cazy.org/GH28.html) | N | N |

| Gene ID | # of Tools | HMMER | DIAMOND | Hotpep | Signal Peptide |
|---|---|---|---|---|---|
| scaffold2\|size463030_77 (domain.php?jobid=2020040810636&gene=scaffold2%7Csize463030_77) | 3 | GH2 (http://www.cazy.org/GH2.html)(6-560) | GH2 (http://www.cazy.org/GH2.html) | GH2 (http://www.cazy.org/GH2.html) | N |
| scaffold2\|size463030_78 (domain.php?jobid=2020040810636&gene=scaffold2%7Csize463030_78) | 3 | GH35 (http://www.cazy.org/GH35.html)(10-321) | GH35 (http://www.cazy.org/GH35.html) | GH35 (http://www.cazy.org/GH35.html) | N |
| scaffold3\|size376888_103 (domain.php?jobid=2020040810636&gene=scaffold3%7Csize376888_103) | 3 | GH65 (http://www.cazy.org/GH65.html)(325-696) | GH65 (http://www.cazy.org/GH65.html) | GH65 (http://www.cazy.org/GH65.html) | N |
| scaffold3\|size376888_11 (domain.php?jobid=2020040810636&gene=scaffold3%7Csize376888_11) | 3 | GH120 (http://www.cazy.org/GH120.html)(299-388) | GH120 (http://www.cazy.org/GH120.html) | GH120 (http://www.cazy.org/GH120.html) | N |
| scaffold3\|size376888_146 (domain.php?jobid=2020040810636&gene=scaffold3%7Csize376888_146) | 2 | GH25 (http://www.cazy.org/GH25.html)(166-348) | GH25 (http://www.cazy.org/GH25.html) | N | N |
| scaffold3\|size376888_163 (domain.php?jobid=2020040810636&gene=scaffold3%7Csize376888_163) | 3 | GH13_31 (http://www.cazy.org/GH13_31.html)(28-372) | GH13_31 (http://www.cazy.org/GH13_31.html) | GH13 (http://www.cazy.org/GH13.html) | N |
| scaffold3\|size376888_165 (domain.php?jobid=2020040810636&gene=scaffold3%7Csize376888_165) | 3 | GH13_31 (http://www.cazy.org/GH13_31.html)(28-376) | GH13_31 (http://www.cazy.org/GH13_31.html) | GH13 (http://www.cazy.org/GH13.html) | N |
| scaffold3\|size376888_225 (domain.php?jobid=2020040810636&gene=scaffold3%7Csize376888_225) | 2 | GT4 (http://www.cazy.org/GT4.html)(181-332) | GT4 (http://www.cazy.org/GT4.html) | N | N |
| scaffold3\|size376888_226 (domain.php?jobid=2020040810636&gene=scaffold3%7Csize376888_226) | 3 | GH3 (http://www.cazy.org/GH3.html)(110-341) | GH3 (http://www.cazy.org/GH3.html) | GH3 (http://www.cazy.org/GH3.html) | Y (1-28) |
| scaffold3\|size376888_276 (domain.php?jobid=2020040810636&gene=scaffold3%7Csize376888_276) | 3 | GT2_Glycos_transf_2 (http://www.cazy.org/GT2_Glycos_transf_2.html)(3-124) | GT2 (http://www.cazy.org/GT2.html) | GT2 (http://www.cazy.org/GT2.html) | N |
| scaffold3\|size376888_69 (domain.php?jobid=2020040810636&gene=scaffold3%7Csize376888_69) | 2 | GH13_11 (http://www.cazy.org/GH13_11.html)(296-488) | GH13_11 (http://www.cazy.org/GH13_11.html) | N | N |
| scaffold4\|size299732_104 (domain.php?jobid=2020040810636&gene=scaffold4%7Csize299732_104) | 2 | GT4 (http://www.cazy.org/GT4.html)(224-376) | GT4 (http://www.cazy.org/GT4.html) | N | N |
| scaffold4\|size299732_109 (domain.php?jobid=2020040810636&gene=scaffold4%7Csize299732_109) | 3 | GT11 (http://www.cazy.org/GT11.html)(1-281) | GT11 (http://www.cazy.org/GT11.html) | GT11 (http://www.cazy.org/GT11.html) | N |
| scaffold4\|size299732_111 (domain.php?jobid=2020040810636&gene=scaffold4%7Csize299732_111) | 2 | GT11 (http://www.cazy.org/GT11.html)(10-256) | GT11 (http://www.cazy.org/GT11.html) | N | N |
| scaffold4\|size299732_113 (domain.php?jobid=2020040810636&gene=scaffold4%7Csize299732_113) | 1 | GT2_Glycos_transf_2 (http://www.cazy.org/GT2_Glycos_transf_2.html)(4-104) | N | N | N |
| scaffold4\|size299732_114 (domain.php?jobid=2020040810636&gene=scaffold4%7Csize299732_114) | 2 | GT17 (http://www.cazy.org/GT17.html)(2-282) | GT17 (http://www.cazy.org/GT17.html) | N | N |
| scaffold4\|size299732_117 (domain.php?jobid=2020040810636&gene=scaffold4%7Csize299732_117) | 1 | GT2_Glycos_transf_2 (http://www.cazy.org/GT2_Glycos_transf_2.html)(10-117) | N | N | N |

| Gene ID | # of Tools | HMMER | DIAMOND | Hotpep | Signal Peptide |
|---|---|---|---|---|---|
| scaffold4\|size299732_119 (domain.php?jobid=2020040810636&gene=scaffold4%7Csize299732_119) | 3 | GT11 (http://www.cazy.org/GT11.html)(1-300) | GT11 (http://www.cazy.org/GT11.html) | GT11 (http://www.cazy.org/GT11.html) | N |
| scaffold4\|size299732_121 (domain.php?jobid=2020040810636&gene=scaffold4%7Csize299732_121) | 1 | GH105 (http://www.cazy.org/GH105.html)(46-385) | N | N | N |
| scaffold4\|size299732_170 (domain.php?jobid=2020040810636&gene=scaffold4%7Csize299732_170) | 2 | GT26 (http://www.cazy.org/GT26.html)(59-231) | N | GT26 (http://www.cazy.org/GT26.html) | N |
| scaffold4\|size299732_171 (domain.php?jobid=2020040810636&gene=scaffold4%7Csize299732_171) | 2 | GT4 (http://www.cazy.org/GT4.html)(183-329) | GT4 (http://www.cazy.org/GT4.html) | N | N |
| scaffold4\|size299732_174 (domain.php?jobid=2020040810636&gene=scaffold4%7Csize299732_174) | 1 | GT2_Glycos_transf_2 (http://www.cazy.org/GT2_Glycos_transf_2.html)(3-174) | N | N | N |
| scaffold4\|size299732_182 (domain.php?jobid=2020040810636&gene=scaffold4%7Csize299732_182) | 1 | N | N | GT1 (http://www.cazy.org/GT1.html) | N |
| scaffold4\|size299732_183 (domain.php?jobid=2020040810636&gene=scaffold4%7Csize299732_183) | 1 | N | N | GT1 (http://www.cazy.org/GT1.html) | N |
| scaffold4\|size299732_185 (domain.php?jobid=2020040810636&gene=scaffold4%7Csize299732_185) | 2 | GT2_Glycos_transf_2 (http://www.cazy.org/GT2_Glycos_transf_2.html)(7-166) | GT2 (http://www.cazy.org/GT2.html) | N | N |
| scaffold4\|size299732_186 (domain.php?jobid=2020040810636&gene=scaffold4%7Csize299732_186) | 1 | GT2_Glycos_transf_2 (http://www.cazy.org/GT2_Glycos_transf_2.html)(6-139) | N | N | N |
| scaffold4\|size299732_213 (domain.php?jobid=2020040810636&gene=scaffold4%7Csize299732_213) | 1 | GT4 (http://www.cazy.org/GT4.html)(188-308) | N | N | N |
| scaffold4\|size299732_228 (domain.php?jobid=2020040810636&gene=scaffold4%7Csize299732_228) | 2 | GT2_Glycos_transf_2 (http://www.cazy.org/GT2_Glycos_transf_2.html)(5-147) | GT2 (http://www.cazy.org/GT2.html) | N | N |
| scaffold4\|size299732_23 (domain.php?jobid=2020040810636&gene=scaffold4%7Csize299732_23) | 1 | CBM2 (http://www.cazy.org/CBM2.html)(479-561)+CBM2 (http://www.cazy.org/CBM2.html)(586-673) | N | N | Y (1-35) |
| scaffold4\|size299732_243 (domain.php?jobid=2020040810636&gene=scaffold4%7Csize299732_243) | 2 | GH25 (http://www.cazy.org/GH25.html)(1116-1296) | GH25 (http://www.cazy.org/GH25.html) | N | N |
| scaffold4\|size299732_252 (domain.php?jobid=2020040810636&gene=scaffold4%7Csize299732_252) | 3 | GT2_Glycos_transf_2 (http://www.cazy.org/GT2_Glycos_transf_2.html)(260-446) | GT2 (http://www.cazy.org/GT2.html) | GT2 (http://www.cazy.org/GT2.html) | N |
| scaffold4\|size299732_253 (domain.php?jobid=2020040810636&gene=scaffold4%7Csize299732_253) | 3 | GT2_Glycos_transf_2 (http://www.cazy.org/GT2_Glycos_transf_2.html)(122-290)+GT2_Glycos_transf_2 (http://www.cazy.org/GT2_Glycos_transf_2.html)(391-586) | GT2 (http://www.cazy.org/GT2.html) | GT2 (http://www.cazy.org/GT2.html) | N |
| scaffold4\|size299732_255 (domain.php?jobid=2020040810636&gene=scaffold4%7Csize299732_255) | 2 | GT2_Glycos_transf_2 (http://www.cazy.org/GT2_Glycos_transf_2.html)(68-238) | GT2 (http://www.cazy.org/GT2.html) | N | N |

| Gene ID | # of Tools | HMMER | DIAMOND | Hotpep | Signal Peptide |
|---|---|---|---|---|---|
| scaffold4\|size299732_256 (domain.php?jobid=2020040810636&gene=scaffold4%7Csize299732_256) | 2 | GT2_Glycos_transf_2 (http://www.cazy.org/GT2_Glycos_transf_2.html)(5-173) | GT2 (http://www.cazy.org/GT2.html) | N | N |
| scaffold4\|size299732_259 (domain.php?jobid=2020040810636&gene=scaffold4%7Csize299732_259) | 2 | GT2_Glycos_transf_2 (http://www.cazy.org/GT2_Glycos_transf_2.html)(5-178) | GT2 (http://www.cazy.org/GT2.html) | N | N |
| scaffold4\|size299732_52 (domain.php?jobid=2020040810636&gene=scaffold4%7Csize299732_52) | 3 | GT2_Glycos_transf_2 (http://www.cazy.org/GT2_Glycos_transf_2.html)(5-166) | GT2 (http://www.cazy.org/GT2.html) | GT2 (http://www.cazy.org/GT2.html) | N |
| scaffold4\|size299732_53 (domain.php?jobid=2020040810636&gene=scaffold4%7Csize299732_53) | 2 | GT2_Glycos_transf_2 (http://www.cazy.org/GT2_Glycos_transf_2.html)(3-125) | GT2 (http://www.cazy.org/GT2.html) | N | N |
| scaffold4\|size299732_58 | 1 | N | GT2 (http://www.cazy.org/GT2.html) | N | N |
| scaffold4\|size299732_59 (domain.php?jobid=2020040810636&gene=scaffold4%7Csize299732_59) | 2 | GT4 (http://www.cazy.org/GT4.html)(224-390) | GT4 (http://www.cazy.org/GT4.html) | N | N |
| scaffold4\|size299732_60 (domain.php?jobid=2020040810636&gene=scaffold4%7Csize299732_60) | 2 | GT4 (http://www.cazy.org/GT4.html)(184-328) | GT4 (http://www.cazy.org/GT4.html) | N | N |
| scaffold4\|size299732_62 (domain.php?jobid=2020040810636&gene=scaffold4%7Csize299732_62) | 2 | GT2_Glycos_transf_2 (http://www.cazy.org/GT2_Glycos_transf_2.html)(9-175) | GT2 (http://www.cazy.org/GT2.html) | N | N |
| scaffold4\|size299732_67 | 1 | N | GT0 (http://www.cazy.org/GT0.html) | N | N |
| scaffold4\|size299732_68 (domain.php?jobid=2020040810636&gene=scaffold4%7Csize299732_68) | 2 | GT2_Glycos_transf_2 (http://www.cazy.org/GT2_Glycos_transf_2.html)(5-164) | GT2 (http://www.cazy.org/GT2.html) | N | N |
| scaffold4\|size299732_69 | 1 | N | GT100 (http://www.cazy.org/GT100.html) | N | N |
| scaffold4\|size299732_77 (domain.php?jobid=2020040810636&gene=scaffold4%7Csize299732_77) | 2 | GT2_Glycos_transf_2 (http://www.cazy.org/GT2_Glycos_transf_2.html)(7-139) | GT2 (http://www.cazy.org/GT2.html) | N | N |
| scaffold4\|size299732_92 (domain.php?jobid=2020040810636&gene=scaffold4%7Csize299732_92) | 2 | GT4 (http://www.cazy.org/GT4.html)(192-337) | GT4 (http://www.cazy.org/GT4.html) | N | N |
| scaffold4\|size299732_96 (domain.php?jobid=2020040810636&gene=scaffold4%7Csize299732_96) | 2 | GT4 (http://www.cazy.org/GT4.html)(225-369) | GT4 (http://www.cazy.org/GT4.html) | N | N |
| scaffold4\|size299732_97 (domain.php?jobid=2020040810636&gene=scaffold4%7Csize299732_97) | 2 | GT4 (http://www.cazy.org/GT4.html)(188-304) | GT4 (http://www.cazy.org/GT4.html) | N | N |
| scaffold4\|size299732_99 (domain.php?jobid=2020040810636&gene=scaffold4%7Csize299732_99) | 2 | GT4 (http://www.cazy.org/GT4.html)(180-337) | GT4 (http://www.cazy.org/GT4.html) | N | N |
| scaffold5\|size271805_10 (domain.php?jobid=2020040810636&gene=scaffold5%7Csize271805_10) | 3 | GH94 (http://www.cazy.org/GH94.html)(2-813) | GH94 (http://www.cazy.org/GH94.html) | GH94 (http://www.cazy.org/GH94.html) | N |
| scaffold5\|size271805_20 (domain.php?jobid=2020040810636&gene=scaffold5%7Csize271805_20) | 3 | GT51 (http://www.cazy.org/GT51.html)(87-266) | GT51 (http://www.cazy.org/GT51.html) | GT51 (http://www.cazy.org/GT51.html) | N |
| scaffold5\|size271805_219 | 1 | N | CBM48 (http://www.cazy.org/CBM48.html)+GH13_9 (http://www.cazy.org/GH13_9.html) | N | N |
| scaffold5\|size271805_225 (domain.php?jobid=2020040810636&gene=scaffold5%7Csize271805_225) | 3 | GH2 (http://www.cazy.org/GH2.html)(28-901) | GH2 (http://www.cazy.org/GH2.html) | GH2 (http://www.cazy.org/GH2.html) | N |

| Gene ID | # of Tools | HMMER | DIAMOND | Hotpep | Signal Peptide |
|---|---|---|---|---|---|
| scaffold5\|size271805_233 (domain.php?jobid=2020040810636&gene=scaffold5%7Csize271805_233) | 2 | GH18 (http://www.cazy.org/GH18.html)(255-556) | GH18 (http://www.cazy.org/GH18.html) | N | N |
| scaffold5\|size271805_235 | 1 | N | GT2 (http://www.cazy.org/GT2.html) | N | N |
| scaffold5\|size271805_60 (domain.php?jobid=2020040810636&gene=scaffold5%7Csize271805_60) | 1 | N | N | GT2 (http://www.cazy.org/GT2.html) | N |
| scaffold6\|size266804_103 (domain.php?jobid=2020040810636&gene=scaffold6%7Csize266804_103) | 3 | GT2_Glycos_transf_2 (http://www.cazy.org/GT2_Glycos_transf_2.html)(7-171) | GT2 (http://www.cazy.org/GT2.html) | GT2 (http://www.cazy.org/GT2.html) | N |
| scaffold6\|size266804_117 | 1 | N | GT4 (http://www.cazy.org/GT4.html) | N | N |
| scaffold6\|size266804_138 (domain.php?jobid=2020040810636&gene=scaffold6%7Csize266804_138) | 1 | GH109 (http://www.cazy.org/GH109.html)(2-116) | N | N | N |
| scaffold6\|size266804_210 (domain.php?jobid=2020040810636&gene=scaffold6%7Csize266804_210) | 2 | CE1 (http://www.cazy.org/CE1.html)(129-361) | N | CE1 (http://www.cazy.org/CE1.html) | N |
| scaffold6\|size266804_211 (domain.php?jobid=2020040810636&gene=scaffold6%7Csize266804_211) | 2 | GH13_18 (http://www.cazy.org/GH13_18.html)(40-400) | GH13_18 (http://www.cazy.org/GH13_18.html) | N | N |
| scaffold6\|size266804_213 (domain.php?jobid=2020040810636&gene=scaffold6%7Csize266804_213) | 2 | CE1 (http://www.cazy.org/CE1.html)(36-262) | N | CE1 (http://www.cazy.org/CE1.html) | N |
| scaffold6\|size266804_225 (domain.php?jobid=2020040810636&gene=scaffold6%7Csize266804_225) | 3 | GT5 (http://www.cazy.org/GT5.html)(3-476) | GT5 (http://www.cazy.org/GT5.html) | GT5 (http://www.cazy.org/GT5.html) | N |
| scaffold6\|size266804_73 (domain.php?jobid=2020040810636&gene=scaffold6%7Csize266804_73) | 2 | CBM48 (http://www.cazy.org/CBM48.html)(127-213)+GH13_9 (http://www.cazy.org/GH13_9.html)(280-572) | CBM48 (http://www.cazy.org/CBM48.html)+GH13_9 (http://www.cazy.org/GH13_9.html) | N | N |
| scaffold6\|size266804_91 (domain.php?jobid=2020040810636&gene=scaffold6%7Csize266804_91) | 3 | GH39 (http://www.cazy.org/GH39.html)(14-435) | GH39 (http://www.cazy.org/GH39.html) | GH39 (http://www.cazy.org/GH39.html) | N |
| scaffold7\|size217497_10 (domain.php?jobid=2020040810636&gene=scaffold7%7Csize217497_10) | 3 | GT35 (http://www.cazy.org/GT35.html)(97-818) | GT35 (http://www.cazy.org/GT35.html) | GT35 (http://www.cazy.org/GT35.html) | N |
| scaffold7\|size217497_108 (domain.php?jobid=2020040810636&gene=scaffold7%7Csize217497_108) | 3 | GH51 (http://www.cazy.org/GH51.html)(8-504) | GH51 (http://www.cazy.org/GH51.html) | GH51 (http://www.cazy.org/GH51.html) | N |
| scaffold7\|size217497_111 (domain.php?jobid=2020040810636&gene=scaffold7%7Csize217497_111) | 3 | GT28 (http://www.cazy.org/GT28.html)(193-342) | GT28 (http://www.cazy.org/GT28.html) | GT28 (http://www.cazy.org/GT28.html) | N |
| scaffold7\|size217497_139 (domain.php?jobid=2020040810636&gene=scaffold7%7Csize217497_139) | 3 | CBM48 (http://www.cazy.org/CBM48.html)(26-110)+GH13_9 (http://www.cazy.org/GH13_9.html)(182-482) | CBM48 (http://www.cazy.org/CBM48.html)+GH13_9 (http://www.cazy.org/GH13_9.html) | GH13 (http://www.cazy.org/GH13.html) | N |
| scaffold7\|size217497_34 (domain.php?jobid=2020040810636&gene=scaffold7%7Csize217497_34) | 3 | GH13_39 (http://www.cazy.org/GH13_39.html)(173-553) | CBM34 (http://www.cazy.org/CBM34.html)+GH13 (http://www.cazy.org/GH13.html) | GH13 (http://www.cazy.org/GH13.html)+CBM34 (http://www.cazy.org/CBM34.html) | N |
| scaffold8\|size203148_102 (domain.php?jobid=2020040810636&gene=scaffold8%7Csize203148_102) | 3 | GH2 (http://www.cazy.org/GH2.html)(43-525) | GH2 (http://www.cazy.org/GH2.html) | GH2 (http://www.cazy.org/GH2.html) | N |

| Gene ID | # of Tools | HMMER | DIAMOND | Hotpep | Signal Peptide |
|---|---|---|---|---|---|
| scaffold8\|size203148_110 | 1 | N | GH13_30 (http://www.cazy.org/GH13_30.html) | N | N |
| scaffold8\|size203148_147 (domain.php?jobid=2020040810636&gene=scaffold8%7Csize203148_147) | 3 | GH10 (http://www.cazy.org/GH10.html)(74-345) | GH10 (http://www.cazy.org/GH10.html) | GH10 (http://www.cazy.org/GH10.html) | N |
| scaffold8\|size203148_16 | 1 | N | AA1 (http://www.cazy.org/AA1.html) | N | N |
| scaffold8\|size203148_74 | 1 | N | CBM50 (http://www.cazy.org/CBM50.html) | N | Y (1-23) |
| scaffold8\|size203148_85 | 1 | N | GT4 (http://www.cazy.org/GT4.html) | N | N |
| scaffold8\|size203148_89 | 1 | N | GH11 (http://www.cazy.org/GH11.html) | N | N |
| scaffold8\|size203148_91 (domain.php?jobid=2020040810636&gene=scaffold8%7Csize203148_91) | 3 | GH67 (http://www.cazy.org/GH67.html)(8-664) | GH67 (http://www.cazy.org/GH67.html) | GH67 (http://www.cazy.org/GH67.html) | N |
| scaffold8\|size203148_92 (domain.php?jobid=2020040810636&gene=scaffold8%7Csize203148_92) | 3 | GH3 (http://www.cazy.org/GH3.html)(32-272) | GH3 (http://www.cazy.org/GH3.html) | GH3 (http://www.cazy.org/GH3.html) | N |
| scaffold8\|size203148_95 (domain.php?jobid=2020040810636&gene=scaffold8%7Csize203148_95) | 2 | GH25 (http://www.cazy.org/GH25.html)(42-211) | GH25 (http://www.cazy.org/GH25.html) | N | Y (1-34) |
| scaffold9\|size160854_151 (domain.php?jobid=2020040810636&gene=scaffold9%7Csize160854_151) | 1 | CE13 (http://www.cazy.org/CE13.html)(64-258) | N | N | N |
| scaffold9\|size160854_154 (domain.php?jobid=2020040810636&gene=scaffold9%7Csize160854_154) | 3 | GH120 (http://www.cazy.org/GH120.html)(297-387) | GH120 (http://www.cazy.org/GH120.html) | GH120 (http://www.cazy.org/GH120.html) | N |

Showing 1 to 128 of 128 entries

First Previous 1 Next Last

Copyright 2017 © YIN LAB (http://bcb.unl.edu), UNL (http://www.unl.edu). All rights reserved. Designed by Tanner Yohe and Le Huang. Maintained by Yanbin Yin. (http://bcb.unl.edu/dbCAN2/about.php)
