## Supplementary file 9 for "Genomic architecture of three newly isolated unclassified *Butyrivibrio* species elucidate their potential role in the rumen ecosystem"

**Table S9.** Source of Horizontal gene transfer (HGT) in *Butyrivibrio* sp. strains from other relative rumen bacteria

| **HGT genes** | **Source** | **Sequence similarity (%)** |
| --- | --- | --- |
| Strain CB08 |  |  |
| Mobile element | Unidentified plasmid plasmid GF1-2_000012F Sequence ID:  [CP021585.1](https://www.ncbi.nlm.nih.gov/nucleotide/CP021585.1?report=genbank&log$=nuclalign&blast_rank=1&RID=C2D3MJ6X016) | 74 |
| Plasmid genes, *parA* | *Clostridium* sp. SY8519 DNA, complete genome  Sequence ID: [AP012212.1](https://www.ncbi.nlm.nih.gov/nucleotide/AP012212.1?report=genbank&log$=nuclalign&blast_rank=1&RID=C2DAEZSC016) | 90 |
| *parB* | *Butyrivibrio proteoclasticus* B316 chromosome 1, complete sequence  Sequence ID: [CP001810.1](https://www.ncbi.nlm.nih.gov/nucleotide/CP001810.1?report=genbank&log$=nuclalign&blast_rank=1&RID=C2DEWDD1014) | 70 |
| Plasmid replication DNA binding-factor 1 | *Butyrivibrio proteoclasticus* B316 chromosome 1, complete sequence  Sequence ID: [CP001810.1](https://www.ncbi.nlm.nih.gov/nucleotide/CP001810.1?report=genbank&log$=nuclalign&blast_rank=1&RID=C2DEWDD1014) | 79 |
| Strain XB500-5 |  |  |
| Mobile element | *Butyrivibrio proteoclasticus* B316 chromosome 2, complete sequence  Sequence ID: [CP001811.1](https://www.ncbi.nlm.nih.gov/nucleotide/CP001811.1?report=genbank&log$=nuclalign&blast_rank=1&RID=C2DPK61J014) | 73 |
| *parA* | *Butyrivibrio hungatei* strain MB2003 chromosome I, complete sequence  Sequence ID: [CP017831.1](https://www.ncbi.nlm.nih.gov/nucleotide/CP017831.1?report=genbank&log$=nuclalign&blast_rank=1&RID=C2DUVYZG014) | 80 |
| *parB* | *Butyrivibrio hungatei* strain MB2003 chromosome I, complete sequence  Sequence ID: [CP017831.1](https://www.ncbi.nlm.nih.gov/nucleotide/CP017831.1?report=genbank&log$=nuclalign&blast_rank=1&RID=C2DWUTVP014) | 79 |
| Plasmid replication DNA binding-factor 1 | *Butyrivibrio hungatei* strain MB2003 chromosome I, complete sequence  Sequence ID: [CP017831.1](https://www.ncbi.nlm.nih.gov/nucleotide/CP017831.1?report=genbank&log$=nuclalign&blast_rank=1&RID=C2DZ9CJ7016) | 78 |
| Strain X503 |  |  |
| *parA* | *Clostridium* sp. SY8519 DNA, complete genome  Sequence ID: [AP012212.1](https://www.ncbi.nlm.nih.gov/nucleotide/AP012212.1?report=genbank&log$=nuclalign&blast_rank=1&RID=C2E2K3K5014) | 72 |
| *parB* | *Butyrivibrio* fibrisolvens 16/4 draft genome  Sequence ID: [FP929036.1](https://www.ncbi.nlm.nih.gov/nucleotide/FP929036.1?report=genbank&log$=nuclalign&blast_rank=1&RID=C2E57CBZ016) | 69 |
| Plasmid replication DNA binding-factor 1 | *Butyrivibrio* hungatei strain MB2003 chromosome I, complete sequence  Sequence ID: [CP017831.1](https://www.ncbi.nlm.nih.gov/nucleotide/CP017831.1?report=genbank&log$=nuclalign&blast_rank=1&RID=C2E6W5DR014) | 78 |
