## Supplementary file 10 for "Genomic architecture of three newly isolated unclassified *Butyrivibrio* species elucidate their potential role in the rumen ecosystem"

**Table S10.** PHAST output, prediction of phage sequences in *Butyrivibrio* spp. strains CB08, XB500-5 and X503.

|  | Region length | Completeness (score) | Region position | Total protein no. | Phage hit protein no. | Possible phage (hit genes count) |
| --- | --- | --- | --- | --- | --- | --- |
| CB08 | 19.7Kb | Intact (110) | 334826-354600 | 21 | 16 | PHAGE_Paenib_PG1_NC_021558(4),PHAGE_Lactob_Ld3_NC_025421(3),PHAGE_Clostr_phiCD506_NC_028838(3),PHAGE_Lactob_c5_NC_019449(3),PHAGE_Lactob_LLKu_NC_022989(3),PHAGE_Paenib_Harrison_NC_028746(2),PHAGE_Paenib_phiIBB_Pl23_NC_021865(2),PHAGE_Clostr_phiCD6356_NC_015262(2),PHAGE_Lactob_Ld17_NC_025420(2),PHAGE_Clostr_phiMMP04_NC_019422(2),PHAGE_Clostr_phiCD481_1_NC_028951(2),PHAGE_Bacter_Diva_NC_028788(2),PHAGE_Clostr_phiCDHM13_NC_029116(2),PHAGE_Paenib_Vegas_NC_028767(2),PHAGE_Bacill_BM5_NC_029069(2),PHAGE_Paenib_HB10c2_NC_028758(2),PHAGE_Bacter_Rani_NC_029084(2),PHAGE_Entero_EFRM31_NC_015270(1),PHAGE_Paenib_Fern_NC_028851(1),PHAGE_Mycoba_Gizmo_NC_021346(1),PHAGE_Pseudo_YuA_NC_010116(1),PHAGE_Lactob_iA2_NC_028830(1),PHAGE_Entero_EFAP_1_NC_012419(1),PHAGE_Strept_Dp_1_NC_015274(1),PHAGE_Bacill_G_NC_023719(1),PHAGE_Entero_IME_EF4_NC_023551(1),PHAGE_Clostr_phiCDHM11_NC_029001(1),PHAGE_Strept_phiSASD1_NC_014229(1),PHAGE_Lactob_PLE2_NC_031036(1),PHAGE_Clostr_c_st_NC_007581(1),PHAGE_Bacill_SP_15_NC_031245(1),PHAGE_Mycoba_ScottMcG_NC_011269(1),PHAGE_Lactob_Ld25A_NC_025415(1),PHAGE_Lactoc_BM13_NC_021861(1),PHAGE_Entero_EfaCPT1_NC_025465(1),PHAGE_Strept_PH10_NC_012756(1),PHAGE_Staphy_phiBU01_NC_026016(1),PHAGE_Lactob_phiLdb_NC_022762(1),PHAGE_Pseudo_MP1412_NC_018282(1),PHAGE_Bacill_1_NC_009737(1),PHAGE_Mycoba_Myrna_NC_011273(1),PHAGE_Entero_phiFL4A_NC_013644(1),PHAGE_Lactob_PL_1_NC_022757(1),PHAGE_Mycoba_Pleione_NC_023737(1),PHAGE_Geobac_GBSV1_NC_008376(1) |
|  | 46.6Kb | Questionable (90) | 227847-274533 | 34 | 19 | PHAGE_Gordon_Zirinka_NC_031097(7),PHAGE_Geobac_E2_NC_009552(4),PHAGE_Lactoc_bIL285_NC_002666(4),PHAGE_Bacill_1_NC_009737(4),PHAGE_Salmon_118970_sal3_NC_031940(3),PHAGE_Geobac_GBSV1_NC_008376(3),PHAGE_Clostr_phiSM101_NC_008265(2),PHAGE_Shigel_SfII_NC_021857(2),PHAGE_Bacill_WBeta_NC_007734(1),PHAGE_Salmon_ST64B_NC_004313(1),PHAGE_Bacill_SP_15_NC_031245(1),PHAGE_Lactob_JCL1032_NC_019456(1),PHAGE_Bacill_PBC1_NC_017976(1),PHAGE_Stx2_vB_EcoP_24B_NC_027984(1),PHAGE_Bacill_Fah_NC_007814(1),PHAGE_Entero_Min27_NC_010237(1),PHAGE_Klebsi_phiKO2_NC_005857(1),PHAGE_Staphy_StB20_like_NC_028821(1),PHAGE_Mycoba_MarQuardt_NC_028798(1),PHAGE_Bacill_phBC6A51_NC_004820(1),PHAGE_Strept_5093_NC_012753(1),PHAGE_Staphy_StB20_NC_019915(1),PHAGE_Bacill_phIS3501_NC_019502(1),PHAGE_Burkho_KS9_NC_013055(1),PHAGE_Lactoc_BK5_T_NC_002796(1),PHAGE_Entero_933W_NC_000924(1),PHAGE_Bacill_Gamma_NC_007458(1),PHAGE_Stx2_c_86_NC_008464(1),PHAGE_Clostr_phiCT19406C_NC_029006(1),PHAGE_Bacill_BM5_NC_029069(1),PHAGE_Clostr_phi8074_B1_NC_019924(1),PHAGE_Strept_PH10_NC_012756(1),PHAGE_Bacill_phi105_NC_004167(1),PHAGE_Entero_SfI_NC_027339(1),PHAGE_Plankt_PaV_LD_NC_016564(1),PHAGE_Yersin_PY54_NC_005069(1),PHAGE_Clostr_phiCT9441A_NC_029022(1),PHAGE_Bacter_Lily_NC_028841(1),PHAGE_Strept_P9_NC_009819(1),PHAGE_Clostr_vB_CpeS_CP51_NC_021325(1) |
|  | 27.6Kb | Incomplete (20) | 257834-285506 | 27 | 13 | PHAGE_Bacill_BCJA1c_NC_006557(5),PHAGE_Entero_EFC_1_NC_025453(5),PHAGE_Paenib_Vegas_NC_028767(5),PHAGE_Erwini_vB_EamM_Phobos_NC_031043(4),PHAGE_Clostr_phiCP39_O_NC_011318(1),PHAGE_Psychr_pOW20_A_NC_020841(1),PHAGE_Brevib_Jimmer1_NC_029104(1),PHAGE_Staphy_187_NC_007047(1),PHAGE_Staphy_SA13_NC_021863(1),PHAGE_Entero_mEp235_NC_019708(1),PHAGE_Thermu_P2345_NC_009803(1),PHAGE_Aeropy_1_NC_028268(1),PHAGE_Lactoc_ul36_NC_004066(1),PHAGE_Thermu_P7426_NC_009804(1),PHAGE_Entero_HK629_NC_019711(1),PHAGE_Natria_PhiCh1_NC_004084(1),PHAGE_Bdello_phi1422_NC_019525(1),PHAGE_Strept_EJ_1_NC_005294(1),PHAGE_Strept_TP1604_NC_028818(1),PHAGE_Bacill_phIS3501_NC_019502(1),PHAGE_Deep_s_D6E_NC_019544(1),PHAGE_Verruc_P8625_NC_029047(1),PHAGE_Strept_YDN12_NC_028974(1),PHAGE_Staphy_phiETA2_NC_008798(1),PHAGE_Brevib_Abouo_NC_029029(1),PHAGE_Aurant_AmM_1_NC_027334(1),PHAGE_Strept_PH15_NC_010945(1),PHAGE_Rhizob_16_3_NC_011103(1),PHAGE_Brevib_Osiris_NC_028969(1),PHAGE_Halovi_HGTV_1_NC_021328(1),PHAGE_Staphy_phiETA3_NC_008799(1),PHAGE_Pseudo_PaBG_NC_022096(1),PHAGE_Lactob_Lj771_NC_010179(1),PHAGE_Geobac_GBSV1_NC_008376(1),PHAGE_Staphy_SA97_NC_029010(1) |
|  | 24Kb | Incomplete (50) | 1180017-1204063 | 33 | 13 | PHAGE_Clostr_phiSM101_NC_008265(2),PHAGE_Staphy_phiPV83_NC_002486(2),PHAGE_Bacill_G_NC_023719(2),PHAGE_Clostr_phiCD211_NC_029048(2),PHAGE_Clostr_PhiS63_NC_017978(2),PHAGE_Staphy_Stau2_NC_030933(1),PHAGE_Synech_Syn5_NC_009531(1),PHAGE_Staphy_80alpha_NC_009526(1),PHAGE_Deep_s_D6E_NC_019544(1),PHAGE_Staphy_vB_SauM_Romulus_NC_020877(1),PHAGE_Staphy_85_NC_007050(1),PHAGE_Staphy_SPbeta_like_NC_029119(1),PHAGE_Thermu_P2345_NC_009803(1),PHAGE_Lactob_jlb1_NC_024206(1),PHAGE_Staphy_53_NC_007049(1),PHAGE_Staphy_2638A_NC_007051(1),PHAGE_Thermu_P7426_NC_009804(1),PHAGE_Staphy_phi2958PVL_NC_011344(1),PHAGE_Staphy_StauST398_4_NC_023499(1),PHAGE_Staphy_vB_SauM_Remus_NC_022090(1),PHAGE_Staphy_SA11_NC_019511(1),PHAGE_Brevib_Sundance_NC_028749(1),PHAGE_Lister_LP_101_NC_024387(1),PHAGE_Staphy_StauST398_3_NC_021332(1),PHAGE_Stx2_c_1717_NC_011357(1),PHAGE_Paenib_Tripp_NC_028930(1),PHAGE_Lactob_phiJB_NC_022775(1),PHAGE_Spirop_SVTS2_NC_001270(1) |
|  | 23.7Kb | Incomplete (20) | 3440891-3464670 | 11 | 6 | PHAGE_Gordon_Bowser_NC_030930(3),PHAGE_Strept_9872_NC_031094(3),PHAGE_Clostr_phiC2_NC_009231(1),PHAGE_Brevib_Jimmer1_NC_029104(1),PHAGE_Klebsi_phiKO2_NC_005857(1),PHAGE_Paenib_Vegas_NC_028767(1),PHAGE_Strept_315.1_NC_004584(1),PHAGE_Brevib_Osiris_NC_028969(1),PHAGE_Brevib_Davies_NC_022980(1),PHAGE_Clostr_phiMMP03_NC_028959(1),PHAGE_Natria_PhiCh1_NC_004084(1),PHAGE_Entero_EF62phi_NC_017732(1),PHAGE_Brevib_Sundance_NC_028749(1),PHAGE_Mycoba_Adler_NC_023591(1),PHAGE_Clostr_CDMH1_NC_024144(1),PHAGE_Clostr_phiMMP01_NC_028883(1),PHAGE_Clostr_phiCD211_NC_029048(1) |
| X503 | 12.5Kb | Incomplete (20) | 78081-90667 | 10 | 8 | PHAGE_Cronob_vB_CsaM_GAP32_NC_019401(2),PHAGE_Bacill_G_NC_023719(2),PHAGE_Entero_vB_KleM_RaK2_NC_019526(1),PHAGE_Cellul_phi38:1_NC_021796(1),PHAGE_Halocy_JM_2012_NC_017975(1),PHAGE_Bacill_AR9_NC_031039(1),PHAGE_Pseudo_YuA_NC_010116(1),PHAGE_Ralsto_RSL1_NC_010811(1),PHAGE_Lister_LP_048_NC_024359(1),PHAGE_Pseudo_MP1412_NC_018282(1),PHAGE_Halovi_HGTV_1_NC_021328(1),PHAGE_Plankt_PaV_LD_NC_016564(1),PHAGE_Bacill_0305phi8_36_NC_009760(1),PHAGE_Cellul_phi14:2_NC_021806(1),PHAGE_Bacill_SPbeta_NC_001884(1),PHAGE_Lister_LMSP_25_NC_024360(1),PHAGE_Staphy_P108_NC_025426(1),PHAGE_Bacill_SP_15_NC_031245(1) |
|  | 16.8Kb | Incomplete (20) | 2183618-2200457 | 14 | 8 | PHAGE_Synech_S_CRM01_NC_015569(1),PHAGE_Bacill_SP_15_NC_031245(1),PHAGE_Clostr_phiC2_NC_009231(1),PHAGE_Pseudo_phiPsa374_NC_023601(1),PHAGE_Strept_phiC31_NC_001978(1),PHAGE_Thermu_P2345_NC_009803(1),PHAGE_Cyanop_KBS_P_1A_NC_020865(1),PHAGE_Caulob_Cr30_NC_025422(1),PHAGE_Thermu_P7426_NC_009804(1),PHAGE_Clostr_phiMMP03_NC_028959(1),PHAGE_Cronob_S13_NC_028773(1),PHAGE_Natria_PhiCh1_NC_004084(1),PHAGE_Halovi_HGTV_1_NC_021328(1),PHAGE_Strept_Jay2Jay_NC_029098(1),PHAGE_Prochl_P_SSP10_NC_020835(1),PHAGE_Mycoba_Myrna_NC_011273(1),PHAGE_Pseudo_PaBG_NC_022096(1),PHAGE_Clostr_phiMMP01_NC_028883(1),PHAGE_Cyanop_P_SSP2_NC_016656(1),PHAGE_Cyanop_9515_10a_NC_016657(1),PHAGE_Clostr_c_st_NC_007581(1),PHAGE_Bacill_phBC6A52_NC_004821(1),PHAGE_Pseudo_VCM_NC_029065(1) |
|  | 8.3Kb | Incomplete (20) | 2549934-2558270 | 8 | 6 | PHAGE_Mycoba_Troll4_NC_011285(1),PHAGE_Clostr_phiCT453B_NC_029004(1),PHAGE_Clostr_phiCD119_NC_007917(1),PHAGE_Bacill_Finn_NC_020480(1),PHAGE_Mycoba_PBI1_NC_008198(1),PHAGE_Entero_c_1_NC_019706(1),PHAGE_Salmon_SP_004_NC_021774(1),PHAGE_Burkho_BcepC6B_NC_005887(1),PHAGE_Clostr_phiCT19406C_NC_029006(1),PHAGE_Mycoba_Gumball_NC_011290(1),PHAGE_Bacill_IEBH_NC_011167(1),PHAGE_Entero_phi80_NC_021190(1),PHAGE_Brocho_BL3_NC_015254(1),PHAGE_Brevib_Davies_NC_022980(1),PHAGE_Bacill_Eoghan_NC_020477(1),PHAGE_Clostr_phiCDHM19_NC_028996(1),PHAGE_Bacill_0305phi8_36_NC_009760(1),PHAGE_Bacill_250_NC_029024(1),PHAGE_Bacill_Blastoid_NC_022773(1),PHAGE_Strept_315.2_NC_004585(1),PHAGE_Bacill_SPbeta_NC_001884(1),PHAGE_Paenib_Tripp_NC_028930(1),PHAGE_Vibrio_X29_NC_024369(1),PHAGE_Mycoba_PLot_NC_008200(1),PHAGE_Geobac_GBSV1_NC_008376(1) |
| XB00-5 | 23Kb | Incomplete (20) | 1269771-1292792 | 9 | 7 | PHAGE_Clostr_phiCD119_NC_007917(1),PHAGE_Bacill_Finn_NC_020480(1),PHAGE_Staphy_StauST398_2_NC_021323(1),PHAGE_Bacill_Andromeda_NC_020478(1),PHAGE_Clostr_phiC2_NC_009231(1),PHAGE_Clostr_phiCD505_NC_028764(1),PHAGE_Staphy_Ipla35_NC_011612(1),PHAGE_Brevib_Abouo_NC_029029(1),PHAGE_Mycoba_Squirty_NC_026588(1),PHAGE_Lister_A500_NC_009810(1),PHAGE_Staphy_vB_SauS_phi2_NC_028862(1),PHAGE_Crocei_P2559Y_NC_023614(1),PHAGE_Brevib_Davies_NC_022980(1),PHAGE_Synech_ACG_2014f_NC_026927(1),PHAGE_Entero_phiFL4A_NC_013644(1),PHAGE_Bacill_SPbeta_NC_001884(1),PHAGE_Helico_KHP30_NC_019928(1),PHAGE_Clostr_phiMMP02_NC_019421(1),PHAGE_Clostr_phiMMP01_NC_028883(1),PHAGE_Lactob_phiJB_NC_022775(1),PHAGE_Helico_phiHP33_NC_016568(1),PHAGE_Helico_KHP40_NC_019931(1),PHAGE_Gordon_Nymphadora_NC_031061(1),PHAGE_Staphy_3A_NC_007053(1),PHAGE_Clostr_phiCD211_NC_029048(1),PHAGE_Geobac_GBSV1_NC_008376(1) |
|  | 9.3Kb | Incomplete (10) | 1573399-1582744 | 8 | 7 | PHAGE_Synech_ACG_2014f_NC_026927(4),PHAGE_Synech_S_SKS1_NC_020851(2),PHAGE_Bacill_SP_15_NC_031245(1),PHAGE_Entero_phi92_NC_023693(1),PHAGE_Mycoba_ScottMcG_NC_011269(1),PHAGE_Escher_phAPEC8_NC_020079(1),PHAGE_Pseudo_phiPsa374_NC_023601(1),PHAGE_Bacill_SPbeta_NC_001884(1),PHAGE_Pseudo_PaBG_NC_022096(1),PHAGE_Bacill_G_NC_023719(1),PHAGE_Pseudo_YuA_NC_010116(1),PHAGE_Pseudo_VCM_NC_029065(1),PHAGE_Prochl_P_SSM2_NC_006883(1),PHAGE_Mycoba_Tonenili_NC_031080(1),PHAGE_Pseudo_MP1412_NC_018282(1) |
|  | 6.9Kb | Incomplete (10) | 2468172-2475151 | 7 | 6 | PHAGE_Synech_S_CAM7_NC_031927(4),PHAGE_Synech_S_SKS1_NC_020851(2),PHAGE_Salmon_ST160_NC_014900(1),PHAGE_Staphy_phiN315_NC_004740(1),PHAGE_Oenoco_phiS13_NC_023560(1),PHAGE_Lactoc_P087_NC_012663(1),PHAGE_Cronob_ENT47670_NC_019927(1),PHAGE_Oenoco_phi9805_NC_023559(1),PHAGE_Cellul_phiST_NC_020842(1),PHAGE_Oenoco_phiS11_NC_023571(1),PHAGE_Prochl_P_SSM2_NC_006883(1) |

**CB08**

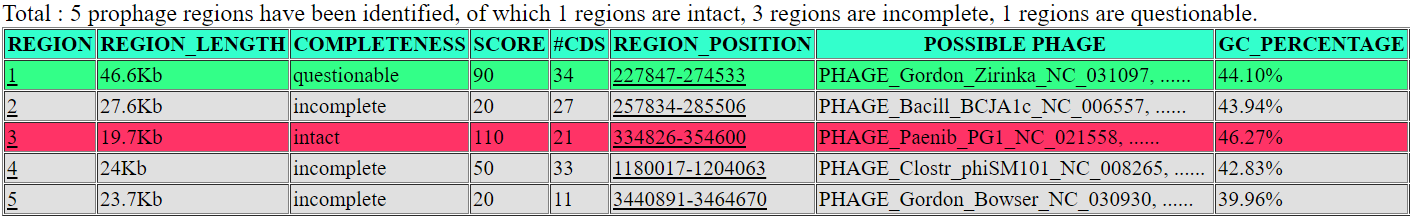

**X503**

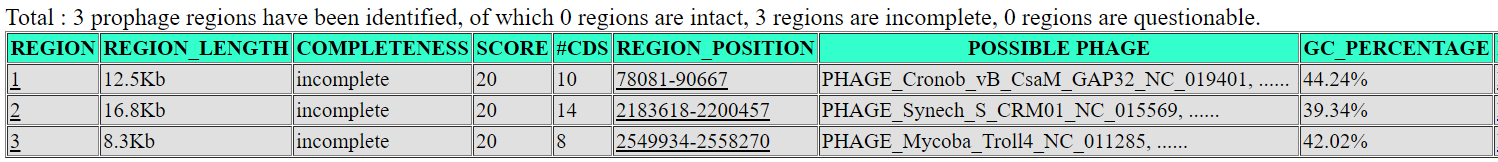

**XB500-5**

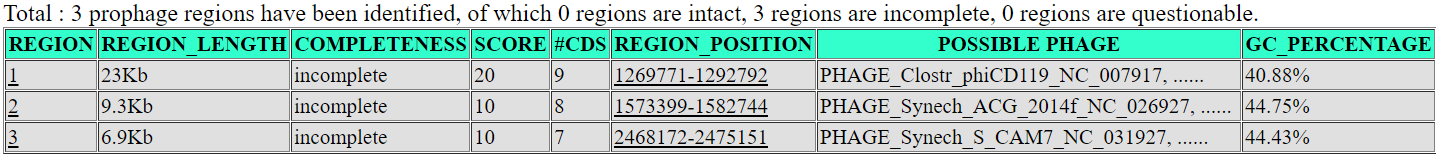
