## Supplementary file legends for "Genomic architecture of three newly isolated unclassified *Butyrivibrio* species elucidate their potential role in the rumen ecosystem"

**Supplementary files legends**

**Supplementary file 1.** Comprehensive genome annotation reports obtained from PATRIC webserver for *Butyrivibrio* spp. strains CB08, XB500-5 and X503.

**Supplementary file 2a.** Circular view of genome sequence of *Butyrivibrio* spp. strains CB08, XB500-5 and X503 obtained from CG viewer; position (Mb) of contigs is labelled as black, forward and reverse coding sequence are marked as green and red respectively whereas blue mark denotes noncoding region; GC content is black graph against yellow.

**Supplementary file 2b.** Comparative circular view of genome sequence of *Butyrivibrio* spp. strains CB08, XB500-5 and X503 with nearest type strains *B. hungatei* DSM 14810^T^ and *B.* *proteoclasticus* B316 ^T^ obtained from BRIG.

**Supplementary file 3.** Summary of 31 bacterial single copy genes detected in the three genomes of *Butyrivibrio* sp. CB08, XB500-5 and X503. Detection of all the 31 single copy genes in the three genomes indicates the completeness of the genome sequencing.

**Supplementary file 4a (Table S4a).** ANI value for pair wise genome comparisons between *Butyrivibrio* spp. (strains CB08, XB500-5 and X503) and type strains *Butyrivibrio* *hungatei* DSM 14810^T^, *Butyrivibrio* *proteoclasticus* B316^T^. The ANI values were inferred using three different ANI calculators. All values are in %; SD = standard deviation.

**Supplementary file 4b (Table S4b).** Comparative account of dDDH, 16S rDNA sequence similarity and GC difference between *Butyrivibrio* spp. (strains CB08, XB500-5 and X503) and type strains *Butyrivibrio* *hungatei* DSM 14810^T^, *Butyrivibrio* *proteoclasticus* B316^T^. All values are in %. Value of dDDH >60%, 16S rDNA sequence similarity >98% and GC difference <1% are shaded in grey.

**Supplementary file 5.** Comprehensive TYGS server reports for *Butyrivibrio* spp. strains CB08, XB500-5 and X503, indicating their novelty.

**Supplementary file 6.** Genome alignment of *Butyrivibrio* spp. strains CB08, XB500-5 and X503 showing similarities within them. The genome alignments and visualization were done using proMauve in PATRIC server (<https://www.patricbrc.org/app/GenomeAlignment>).

**Supplementary file 7.** Distribution of proteins in various COG categories of *Butyrivibrio* spp. strains CB08, XB500-5 and X503 and nearest type strains *B. hungatei* DSM 14810^T^ and *B.* *proteoclasticus* B316 ^T^.

**Supplementary file 8.** Report generated for CAZyme prediction from genomes of *Butyrivibrio* spp. strains CB08, XB500-5 and X503 using dbCAN server.

**Supplementary file 9 (Table S9).** Source of horizontal gene transfer (HGT) in Butyrivibrio spp. strains CB08, XB500-5 and X503 from other relative rumen bacteria. The HGT were predicted using IslandViewer4 ([https://www.pathogenomics.sfu.ca/islandviewer](https://www.pathogenomics.sfu.ca/islandviewer/)), prediction methods used: Integrated, IslandPath-DIMOB and SIGI-HMM.

**Supplementary file 10.** Prediction of phage viral protein coding sequencing in the three Butyrivibrio spp. strains CB08, XB500-5 and X503. The results were obtained from PHAST webserver ([http://phast.wishartlab.com](http://phast.wishartlab.com/)).
